# Deep Learning Reveals How Cells Pull, Buckle, and Navigate Tissue-Like Environments

**DOI:** 10.1101/2022.10.24.513423

**Authors:** Abinash Padhi, Arka Daw, Medha Sawhney, Maahi M. Talukder, Atharva Agashe, Mehran Mohammad Hossein Pour, Mohammad Jafari, Guy M. Genin, Farid Alisafaei, Sohan Kale, Anuj Karpatne, Amrinder S. Nain

## Abstract

Cells move within tissues by pulling on and reshaping their fibrous surroundings. Measuring the associated forces has been a fundamental challenge in cell biology. Here, we develop deep-learning-enabled live-cell fiber-force microscopy (DLFM), which computes forces produced by living cells in real time as they interact with tissue-like fiber networks. DLFM combines basic phase microscopy with novel deep learning to simultaneously track cell movement and fiber deformation without disruptive fluorescent labels or chemical modifications. This allowed us to measure forces in real-time situations that were previously impossible to study, revealing an intricate mechanical landscape: cells generate ten-fold changes in force as they change shape during migration, create force-dipoles during cell-cell interactions, and dramatically alter their force patterns during stem cell differentiation. Through integrated experiments and mathematical modeling, we discovered that cells in fibrous environments form force-generating adhesions throughout their body, strikingly different from the edge-only adhesions seen in traditional petri dish experiments. Results clarify cytoskeletal pathways by which cells adapt force-generating machinery to navigate the fibrous architecture of tissues.

## Introduction

Cells in solid tissues interact with a fibrous extracellular matrix (ECM) through matrix adhesions that remain poorly characterized, hindering our understanding of how forces regulate cell function in physiological environments ^1–7^. Quantifying the dynamic mechanical feedback between cells and surrounding fibrous networks has proved challenging due to difficulties in simultaneously resolving deformations of living cells and ECM in systems where cells are cultured on a deformable fibrillar background. Furthermore, a lack of robust inverse methods to rapidly infer spatially-resolved cellular tractions from matrix deformations has hindered the investigation of cell-ECM mechanics. To address these limitations, we developed a novel deep learning-enabled technique that measures forces exerted by unlabeled living cells on fibrous matrices in real time. Our approach, termed deep-learning-enabled live-cell fiber-force microscopy (DLFM), utilizes a deep-learning model trained to enhance the contrast of low-resolution, unlabeled cells against the deforming fiber background, which allows the extraction of cell-induced ECM displacements, which are converted to mechanical tractions using inverse finite-element methods.

The mechanobiological questions that motivate this work are the questions of why cell-matrix adhesions in a fibrous system differ so fundamentally from the focal adhesions observed in cells on flat substrates such as Petri dishes and how these focal adhesions can remain stable. Focal adhesions typically form on the periphery of cells on flat, continuous substrates, with stability requiring shear forces on the focal adhesions ^8^, as has been established through traction microscopy techniques that estimate these forces from how cells deform their substrates ^9–14^. However, cells in fibrous matrices have shapes and matrix adhesions that differ fundamentally, with major adhesions often appearing proximal to the ends of protrusions ^15–24^. We previously extended these inverse traction estimation approaches to cells in 3D hydrogels ^25^ and to fibrous matrices ^16,26^, but these approaches have thus far failed to elucidate answers to these questions. Challenges are that the 3D hydrogel systems, which allow for excellent spatial resolution, have focal adhesions that differ from those seen in fibrous systems. In contrast, models for estimating tractions on fibrous systems have accuracy and spatial resolution that are inadequate for tracking focal adhesion dynamics. Regular arrays of well-defined fibers have been used to recapitulate critical aspects of 3D cell mechanobiology ^15,24,27–30^. However, simultaneously tracking deformations of cells and the fibers they cover is a computer vision challenge that has not yet been solved.

Simultaneously tracking deforming objects, such as cells, on a deforming background is a central challenge for quantitative image analysis in deep learning. Numerous deep learning approaches can track deforming structures on static backgrounds ^31–35^. For the problem in which both the semi-transparent cells and fibrous, non-uniform background deform, segmentation is possible, but not at the level of resolution needed for tracking mechanical fields ^36^, and all existing methods face challenges in this regard. Deep learning-based methods require extensive labeled data ^37^. Optical flow methods rely heavily on apparent motion, which can be ambiguous when objects and backgrounds share transparency and deformation characteristics ^38^. Active contour models suffer from issues of segmentation for partially transparent objects ^39^. Generative adversarial networks (GANs) are computationally intensive to train and may be unstable when trying to capture the nuances of transparency and deformation ^40^. Integrating physics-based models can often reduce the complexity of an artificial intelligence algorithm ^41^. Still, these have much less utility in cases such as tracking cells, where the underlying biophysics is much less clear. However, integrating phase-contrast microscopy with deep learning-based techniques, especially GANs, has allowed "de-crappification" of otherwise "crappy" images in other contexts ^42^, including prediction of where fluorescent labels would appear within and around an unlabeled cell imaged using only phase contrast microscopy ^31,43,44^. We, therefore, developed a GAN and trained on a system of parallel labeled training image datasets, thereby sidestepping issues of cell translucency and simultaneous foreground and background deformation. We applied the resulting technique in conjunction with a chemomechanical model of interactions between cells and fibers to identify how tractions at the cell-matrix interface regulate the internal matrix adhesions characteristic of cells in fibrous networks and discover dramatic changes in forces during critical processes of migration and stem cell differentiation.

## Results

### Non-uniform, out-of-plane stresses enable adhesions to form away from the cell periphery

To understand how cells exert forces on their surroundings, we first examined how they form connections with the extracellular matrix (ECM). Cells generate forces through structures focal adhesions, molecular assemblies that physically anchor the cell to the ECM. While previous studies using flat surfaces (2D hydrogels) showed that these adhesions form primarily at the cell edge, a fundamentally different pattern emerged in our tissue-like fibrous environments. These were evident in human mesenchymal stem cells (hMSCs) migrating on networks of orthogonal fibers, which instead presented adhesions throughout the cell surface, far from the periphery (**Fig. 1B** and **Supplementary Movie1**).

**Figure 1.**
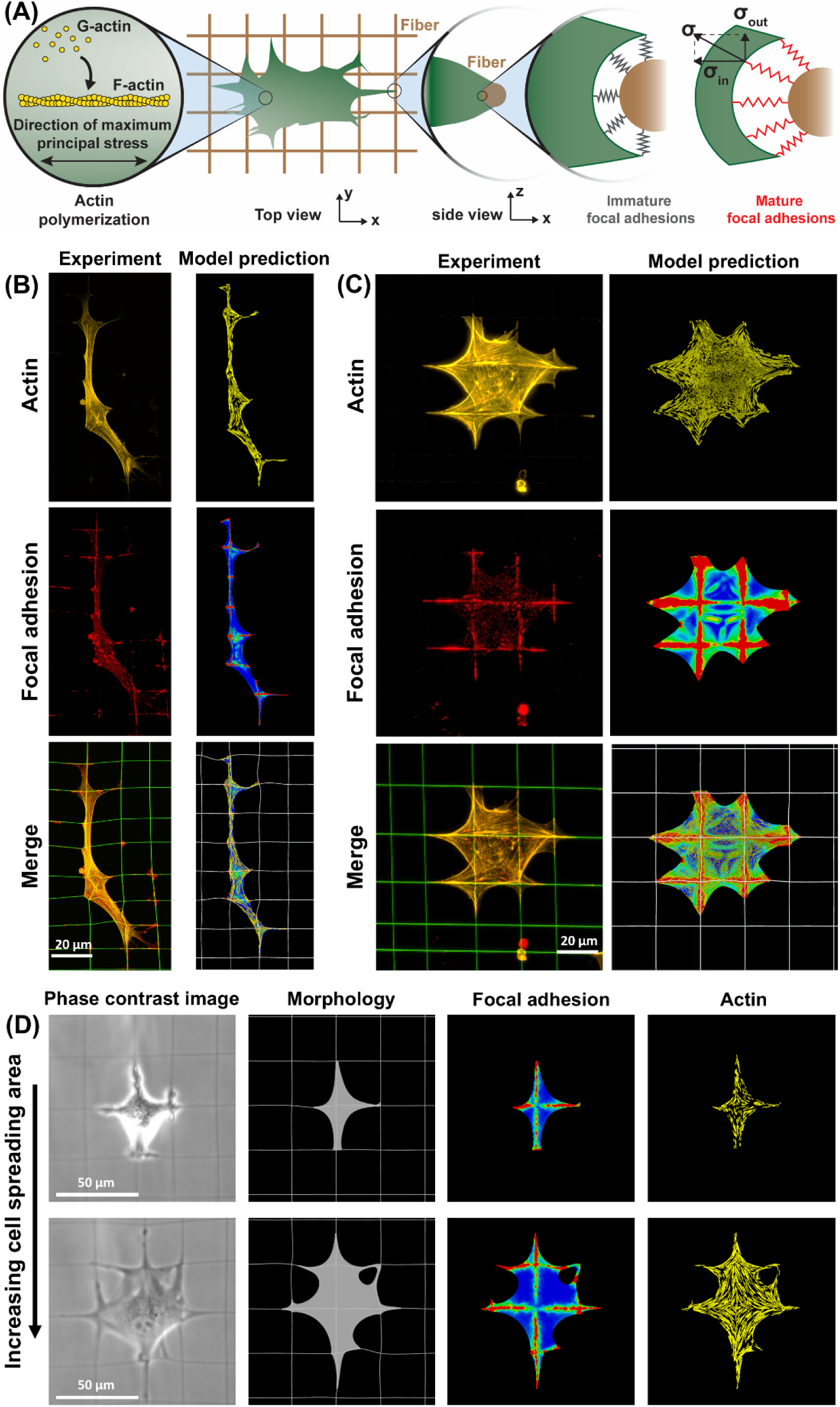
Tensile stresses generated within the cytoskeleton and at the cell-matrix interface regulate the formation of actomyosin structures and mature focal adhesions. (A) Using an active contractile cell model, we simulated cell contraction on a crosshatch fiber network. Simulations began with two initial conditions: (i) uniform, isotropic cell contraction, with a uniform distribution of myosin and actin densities throughout the cell in all directions, and (ii) a uniform and compliant layer of immature focal adhesions with weak cell-fiber connections. After that, cell contraction generated non-uniform tension within the cytoskeleton and at the cell-matrix interface, depending on cell morphology and spreading, promoting actomyosin expression and actin fiber formation in the direction of maximum principal cytoskeletal stress. Tension at the cell-fiber interface matured focal adhesions connecting the cell to the fibers. (B-C) Simulation results for two representative cells, one elongated and one well-spread. The top panels depict the direction of maximum principal stress in the cytoskeleton, indicating the orientation of actin fiber formation in the model. The middle panels show a heatmap of maximum principal stress at the cell-fiber interface, demonstrating the formation of mature focal adhesions in the model. Model predictions for the formation of actin and focal adhesions were validated against staining for actin (phalloidin) and paxillin, respectively. These results confirmed that actin forms along the direction of maximum principal stress, and unlike focal adhesions on 2D hydrogels, which typically form at the cell periphery, focal adhesions in fibrous networks can also form at locations distant from the cell edge. Notably, the mechanical stresses generated by cell contraction regulate both actin organization and focal adhesion formation. (D) Our model showed that these mechanical stresses — and, consequently, the formation of focal adhesions and actin networks — are highly dynamic, changing significantly as cells alter their spreading area and shape. Scale bars: (B, C) 20 µm, (D) 50 µm.

We used a chemomechanical cell model to investigate the mechanisms behind this distinct focal adhesion formation in fibrous ECMs, previously validated on 2D nonfibrous ECMs (**Supplementary Fig. 1**) ^45^. The model incorporates key cytoskeletal components involved in adhesion, including myosin, actin, microtubules, and focal adhesions (**Supplementary Notes 1-2** and Materials and Methods). Central to the model is the hypothesis that actomyosin contractility and focal adhesion formation are regulated by mechanical tension generated within the cytoskeleton and at the cell-ECM interface, respectively.

Simulations began with two initial conditions: (i) uniform and isotropic cell contraction, reflecting a uniform initial distribution of myosin and actin densities, both spatially and in terms of orientation distribution, and (ii) a uniform, compliant layer of focal adhesions, modeling immature adhesions and weak cell-ECM connections (**Fig. 1A**). Starting from these conditions, the model predicted that cell contraction would stretch immature focal adhesions (**Fig. 1A**). Prolonged stretching matured the adhesions to a stiffer state, creating a stronger link between the cell and ECM that resisted further contraction and upregulated cytoskeletal tension. This tension promoted actomyosin contractility,^46^ stiffening actin along the direction of maximum principal stress and representing actin filament formation (**Fig. 1A**).

Consistent with our experimental observations, the simulations showed that mature focal adhesions were not limited to the cell periphery and could form at a considerable distance from the cell periphery along nanofibers (**Figs. 1B-C**). The model revealed that the formation of these non-peripheral focal adhesions was regulated by out-of-plane stresses exerted on nanofibers. Suppressing this by restricting cells to 2D nonfibrous ECM or by allowing only in-plane stresses at the cell-fiber interface in the simulations caused focal adhesions to migrate to the cell periphery (**Supplementary Fig. 2**), as seen on traditional 2D nonfibrous substrates^45,47,48^.

The model also predicted that actin fibers would form in the directions of maximum principal stress. We found close agreement between predicted orientations and experimental images (**Fig. 1B-C and Supplementary Figs. 3-4**), further validating the model and supporting our hypothesis on the central role of mechanical tension in directing cell-ECM interactions. However, our simulations demonstrated that mechanical stress — and consequently, the formation of focal adhesions and actin networks — is highly dynamic, changing continuously as cells adjust their spreading area and shape during migration (**Fig. 1D**). Yet, no existing approach allows for real-time measurement of cell-generated stress on fibrous ECMs. This limitation motivated us to develop the first real-time technique to measure cell traction forces in fibrous matrices.

### Estimation of cellular forces from phase contrast images

The initial step in quantifying cellular forces requires measuring the deformation of ECM fibers induced by cell-generated forces. To resolve deformations of an initially orthogonal fiber network with hMSCs migrating on it, we developed a conditional generative adversarial network (cGAN) deep learning model for reconstituting fiber geometry from phase contrast images (**Fig. 2A**, **Supplementary movies M2, M3**). We developed a "pix2pix" cGAN to resolve the fiber deformations by training the model on paired phase contrast and fluorescence microscopy images of cells deforming fluorescently labeled fibers^49–51^ (**Fig. 3Ai**). Phase contrast images of fibers were obscured by cells, but fluorescence images were not (**Figs. 2Ai and 2Aii**). The cGAN’s generator, G: x → y, was trained to map a phase contrast input image x to a fluorescence image y. The cGAN’s discriminator, D, was trained to distinguish between real images y and reconstituted images G(x). After training the cGAN on a manually curated data set (**Supplementary Note 3**), it robustly reconstituted fluorescent images of deformed fiber networks from experimental phase-contrast images (**Fig. 3Aii, Supplementary movies M4, M5**).

**Figure 2.**
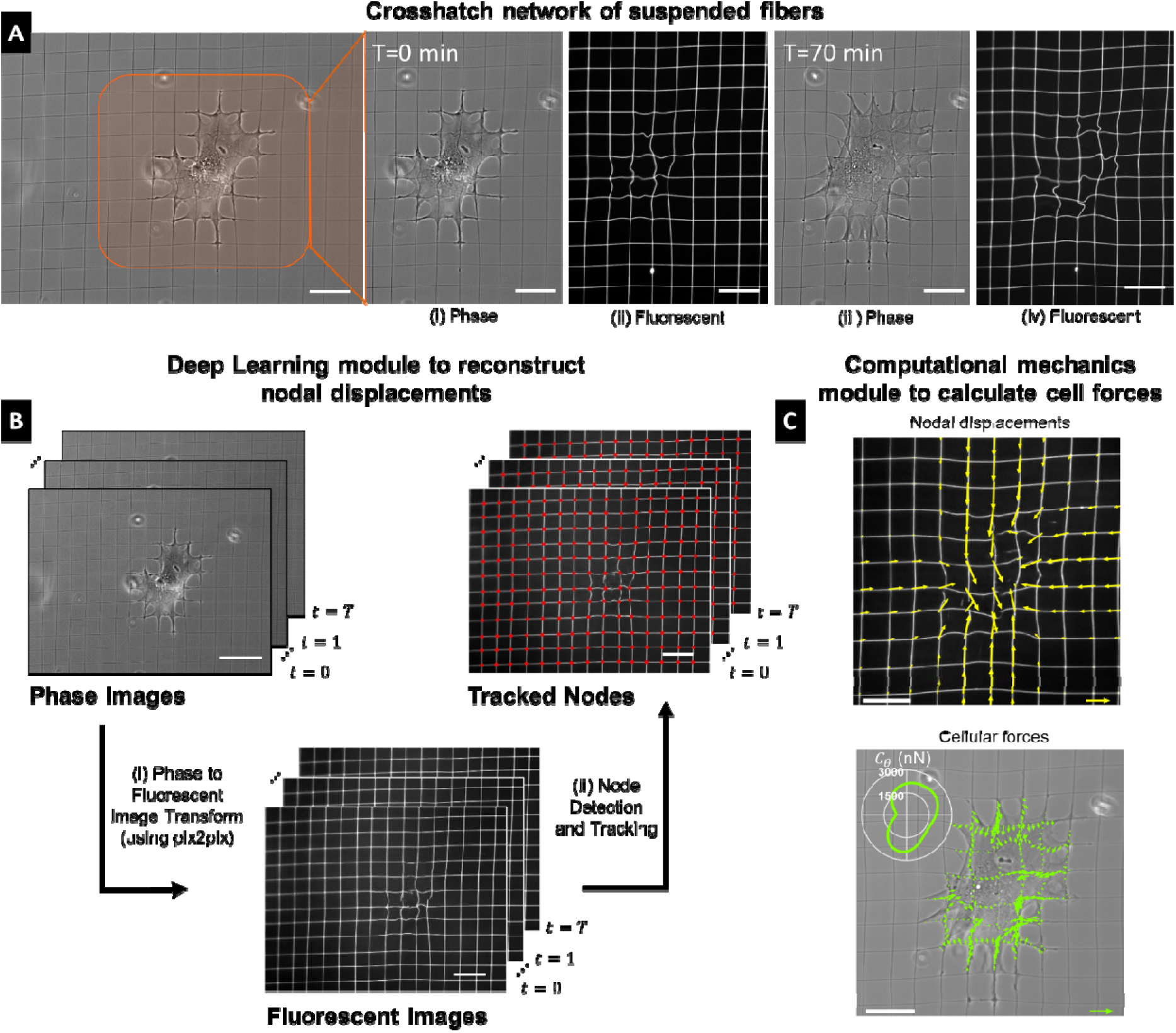
Cell force computation algorithm. (A) Cells migrate over suspended crosshatch fiber networks, deforming the underlying fiber network, as seen in experimentally obtained phase-contrast (i and iii) and corresponding fluorescence microscopy images (iii and iv). Experimental fluorescent images were used exclusively to train the deep learning modules. (B) (i) Phase contrast images were transformed into synthetically generated fluorescent images through a deep learning-based model (pix2pix). (ii) A deep learning-based object detection model, RetinaNet, and a bi-directional tracking algorithm identified nodal intersection points of the fiber network (red dots added for visualization). (C) Displacement fields were calculated using the extracted nodal points (yellow arrows). Cell forces computed from displacement fields were represented as line tension (green arrows). Force polarity maps ( , inset) captured the orientational distribution of cellular forces. Scale bars: 50μm, Yellow arrow: 5 µm, Green arrow: 35 mN/m

**Figure 3.**
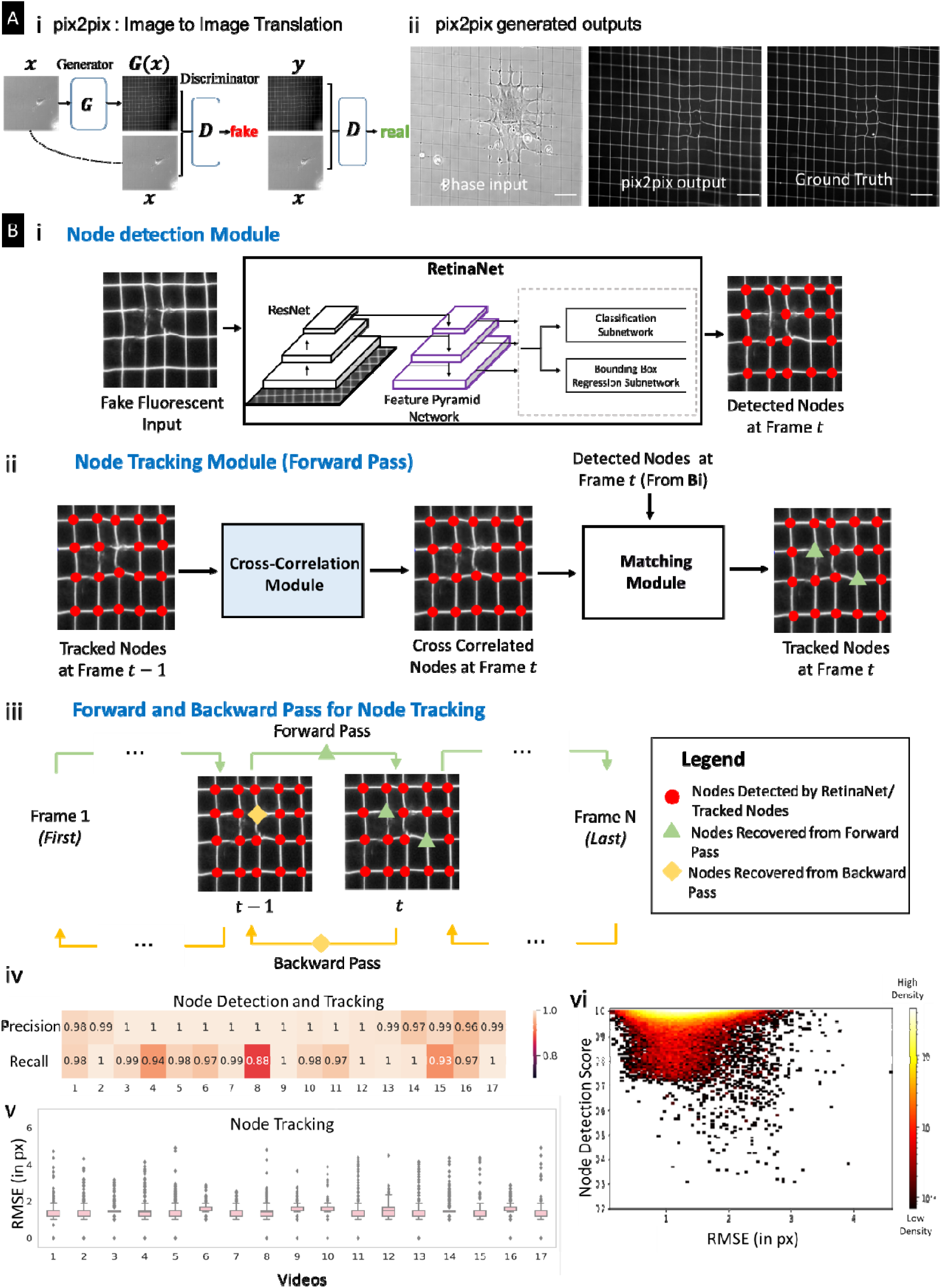
Deep-learning-based image reconstruction and feature extraction. (A) (i) Schematic representation of our pix2pix model, which generates fluorescent images from input phase-contrast experimental images (ii) An example of a synthetically generated fluorescent image obtained from a trained pix2pix model. Scale bar: 50 µm (B) (i) Node Detection Module that uses a deep-learning-based object detection model called *RetinaNet* to identify grid intersections/nodes from synthetically generated fluorescent images. (ii) An illustration of the forward pass of the node tracking algorithm, (iii) A schematic representation of the forward and backward passes of the proposed bi-directional Tracking algorithm, (iv) *Precision* and *Recall* values of the predicted nodes after node detection and tracking are close to ∼1 across all 17 videos, indicating close to ideal performance. (v) The distribution of Nodal Errors observed after node detection and tracking in each video shows that the average nodal error is less than 2 pixels, comparable to human measurement error. (vi) The scatter plot of node detection confidence scores vs nodal errors shows high confidence (greater than 0.7) of the RetinaNet for most of the predicted nodes with low nodal errors.

These reconstituted images were analyzed to identify fiber intersection "nodes" (red dots, **Fig. 3Bi**) and to track them across frames (**Fig. 2Bii**) using the deep learning-based object detection model *RetinaNet*^52^. To improve accuracy, we developed a bi-directional tracking algorithm. For each timeframe t_i_, a cross-correlation based forward nodal tracking algorithm (**Fig. 3Bii**) was used to estimate nodal mapping using positions from the previous timeframe t_i-1_. Candidate cross-correlated nodes were combined with nodes detected by *RetinaNet* using a matching algorithm (**Supplementary Fig. 7**) that mapped intersections from t_i-1_ to t_i_. This was repeated to map intersections from t_i+1_ back to t_i_, resulting in a complete bi-directional tracking algorithm (**Fig. 3Biii**; see Materials and Methods for details). Nodal positions were then used to calculate fiber deformations. An inverse problem was solved to estimate force fields from these fiber deformations (**Fig. 2C**).

To assess the effectiveness of node tracking, we analyzed synthetically generated images. Precision and recall^53^ were computed, with precision defined as the ratio of the number of correctly identified nodes to the total number of nodes identified and recall defined as the ratio of the number of nodes correctly identified to the actual number of nodes (ground truth). Precision and recall were both near the ideal value of 1 for seventeen videos analyzed (**Fig. 3Biv**), demonstrating adequate node detection and tracking.

The spatial accuracy of node tracking, defined as the average root mean squared error (RMSE) of the Euclidean distance between experimentally estimated and ground truth node positions (**Fig. 3Bv**), was below 2 pixels, on the order of expected human measurement error (**Supplementary Fig. 8**). RMSE was relatively small for nodes with a high "Node Detection" confidence score (>0.7) generated by *RetinaNet* and the RMSE across all 44,852 detected nodes (**Fig. 3Bvi**), prompting us to eliminate detected nodes with a confidence score less than 0.7 in subsequent analyses.

We needed a reference map of where the fibers would be without any cell-generated forces to calculate how much cells deformed the fibers. We created this reference map by taking advantage of the precise geometric pattern of our crosshatch fiber networks (**Supplementary Note 4, Supplementary Fig. 9**). This represents a key advantage over traditional force measurement methods, which require adding drugs to temporarily paralyze cells to obtain such reference measurements.

### Estimation of forces by an inverse mechanics formulation

We developed an inverse formulation to infer forces from the nodal displacements and deformed fiber shapes (see Methods). To validate this and test its accuracy, we generated nodal displacement data (u_m_) at grid intersections by applying representative force patterns to a computational model of the crosshatch fiber network (red arrows, **Fig. 4A**) and simulated measurement errors (**Supplementary Fig. 8**) by adding zero-mean Gaussian noise of varying strength (*σ_e_* = {0.0 μm, 0.1 μm, 0.2 μm, 0.5 μm}) to ***u***_*m*_. An example of the recovered force pattern obtained by solving the regularized inverse problem for *σ_e_* = 0.2 μm (green arrows, **Fig. 4A**) recovered force patterns with less than 10% error (**Fig. 4Bii**), where the forces are compared after summing within a circular patch of 20 μm radius. Averaged measurements of the force patterns (overall contractility C, see Methods) were recovered with even higher accuracy than pointwise measurements (**Supplementary Note 5**). A regularization parameter *β* (*i.e.,* parameter to penalize large, unphysical forces in the inverse solution) was chosen based on an L-curve (**Fig. 4Bi**) of reconstructed force magnitude ||**f**|| versus normalized matching errors for a range of *β* values. Although a typical L-curve displays a sharp increase in ||**f**|| as *β* approaches zero because of overfitting the solution to noise, this was absent in our L-curves because the image-based fitting term constrains crosshatch fiber network deformations to the fluorescent fiber shapes. The level of noise *σ_e_* in the measured nodal displacements was relatively low (vertical dotted lines, **Fig. 4Bi**). The inverse solver reduced normalized matching errors within this noise level (see **Supplementary Note 5** for detailed analysis and more validation examples).

**Figure 4.**
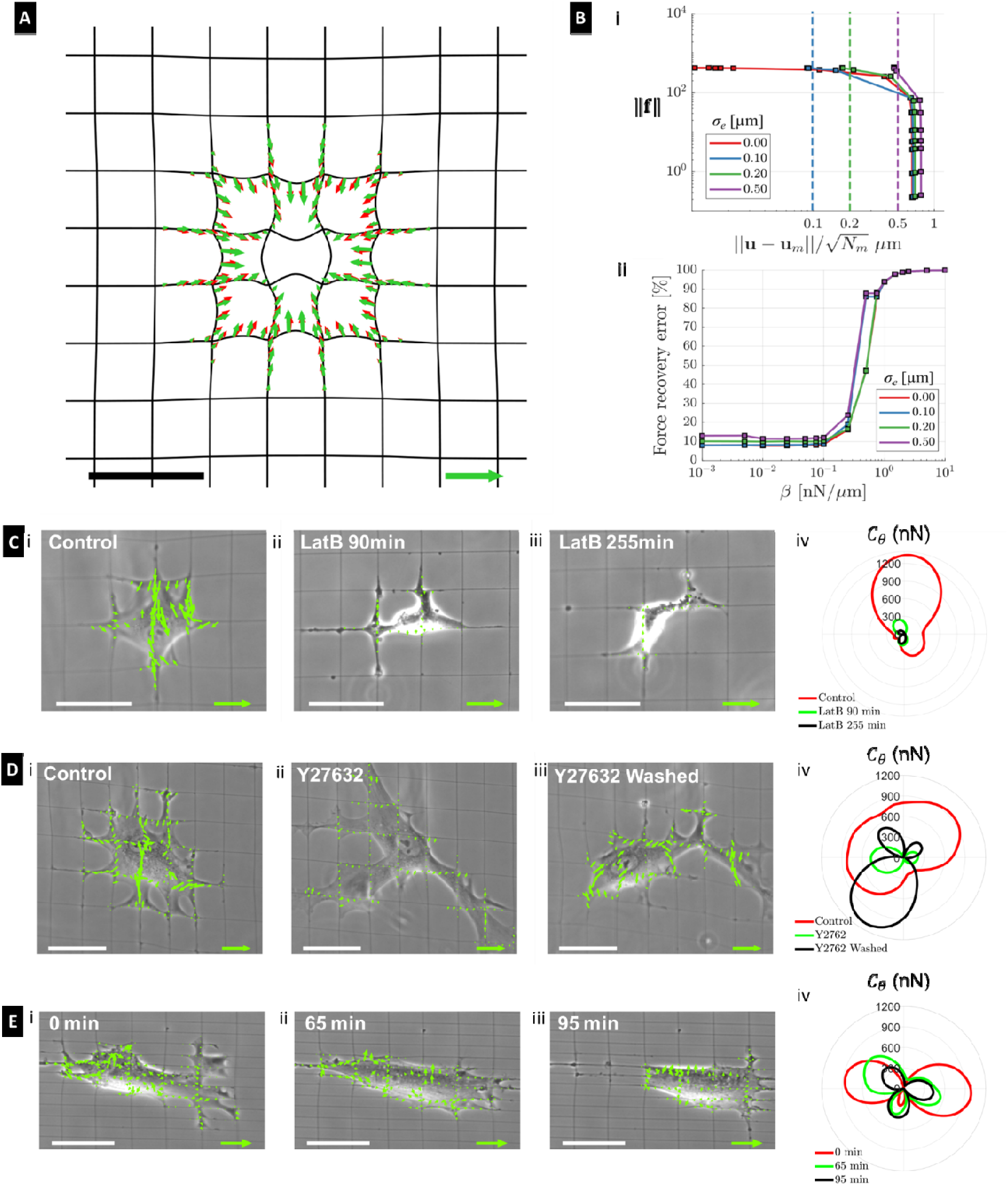
Inverse formulation to estimate forces on crosshatch fiber networks. (A) Validation of the inverse formulation with a representative radial contractile force pattern (red arrows) generating the displacements at grid intersections. The inverse problem is solved using with zero-mean Gaussian noise of standard deviation. The solution obtained for a typical noise strength μm is shown with green arrows. (B) Analysis of the validation example in (A) for different σ_e_. (i) The L-curves showing the magnitude of the force pattern ||**f**|| as a function of the displacement matching error ||**u - u**_*m*_|| normalized by the number of measured nodes *N_m_* for a range of regularization parameter *β* from 10 to 10^-3^ nN/μm. (ii) Recovery of forces as a function of *β* for different σ_e_. The forces are compared after summing up within a patch of 20 μm radius (see **Supplementary Note 5**). (C) Line tension map of the cell (i) under control condition, presence of Latrunculin B (actin inhibitor) after (ii) 90min, and (iii) 255min. (iv) Force polarity maps capturing the orientational distribution of cellular forces. (D) Line tension map of the cell (i) under control condition, (ii) in the presence of Y27632 drug (ROCK inhibitor), and (iii) after drug wash-off showing recovered contractility. (iv) Force polarity maps for cases i-iii. (E) i-iii. Line tension map of cells migrating in aligned morphology on rectangular crosshatch fibers tracked over tens of minutes. (iv) Force polarity maps capturing the horizontal alignment of cellular forces. Green tension scale arrow: (A) 25 mN/m, (C-E) 35 mN/m. Black/White scale bars: (A, C-E) 50 μm.

We applied these validated inverse methods to estimate forces exerted by cells migrating over crosshatch fiber networks. Phase contrast images were interpreted using our cGAN (**Fig. 3**) to generate a corresponding fluorescent image to calculate nodal deformations required for calculating cell forces (**Supplementary Figs. 9-10**). Estimated nodal forces scaled with the length of finite elements used to discretize the nanofibers and reported as line tensions (force per unit element length in mN/m, **Supplementary Fig. 10C**). Line tensions in the range of 5-25 mN/m were observed, which for a representative 10 µm adhesion patch on fibers results in 50-250 nN force, in agreement with other reports for cells on fibers ^24,54^, and on the same order as forces per adhesion patch for cells on soft gels (2-50 nN ^55,56^), fibrous networks (100-400 nN ^24,28^), and micropost arrays (1-80 nN ^55,57–59^). Force polarity maps of the contractility C_8_ (**Supplementary Fig. 10D**, methods) showed spatial organization of cellular contractility.

### Cellular contractility in dynamic ECM interactions and stem cell differentiation

To verify our method, we tested it in several biologically relevant scenarios. First, we verified that DLFM could detect changes in cell forces when we added drugs that target the cell’s force-generating machinery. When we disrupted the cell’s internal scaffolding using Latrunculin B, forces decreased as expected (**Fig. 4C, Supplementary movie M6**). Similarly, when we inhibited ROCK (Rho-associated kinase, a key regulator of cell contraction) using Y-27632, forces dropped within 30 minutes (**Fig. 4Di, ii, Supplementary movie M7**). When we removed these drugs, cells recovered and restored their original force patterns and magnitudes (**Fig. 4Diii, iv**).

Next, we inquired how contractility varied as cells migrated and changed shapes. Previously, we demonstrated that the spacing of fibers in crosshatch fiber networks controls cell shapes and migration direction ^30,60,61^. On rectangular (25 μm x 8 μm) grids (**Fig. 4E, Supplementary movie M8**), cells elongated and applied forces that were mainly concentrated at their leading and trailing ends (**Fig. 4Ei-iii**), with forces transverse to the fibers lower than those parallel to the fibers (**Fig. 4Eiv**), as expected for polarized cell migration ^64,65^. On a network with square (30 μm X 30 μm) grids transitioning into rectangular (30 μm x 15 μm) grids, cells spread in the square regions and elongated in the rectangular regions (**Fig. 5A, Supplementary movie M9**). As cells migrated across regions, a low and high contractility migration cycle was observed, with contractility reducing by an order of magnitude as cells transitioned from spread to elongated morphologies. The contractility of hMSCs was on the order of values reported on PDMS micropost arrays (1000 -5000 nN, depending on cell area ^48,66^) and polyacrylamide gels (∼600-6000 nN ^67^, depending on substrate stiffness). Cell-cell interactions led to net increases in contractility (**Fig. 5B** and **Supplementary movie M10**) as cells cycled between high and low contraction while changing shape during migration.

**Figure 5.**
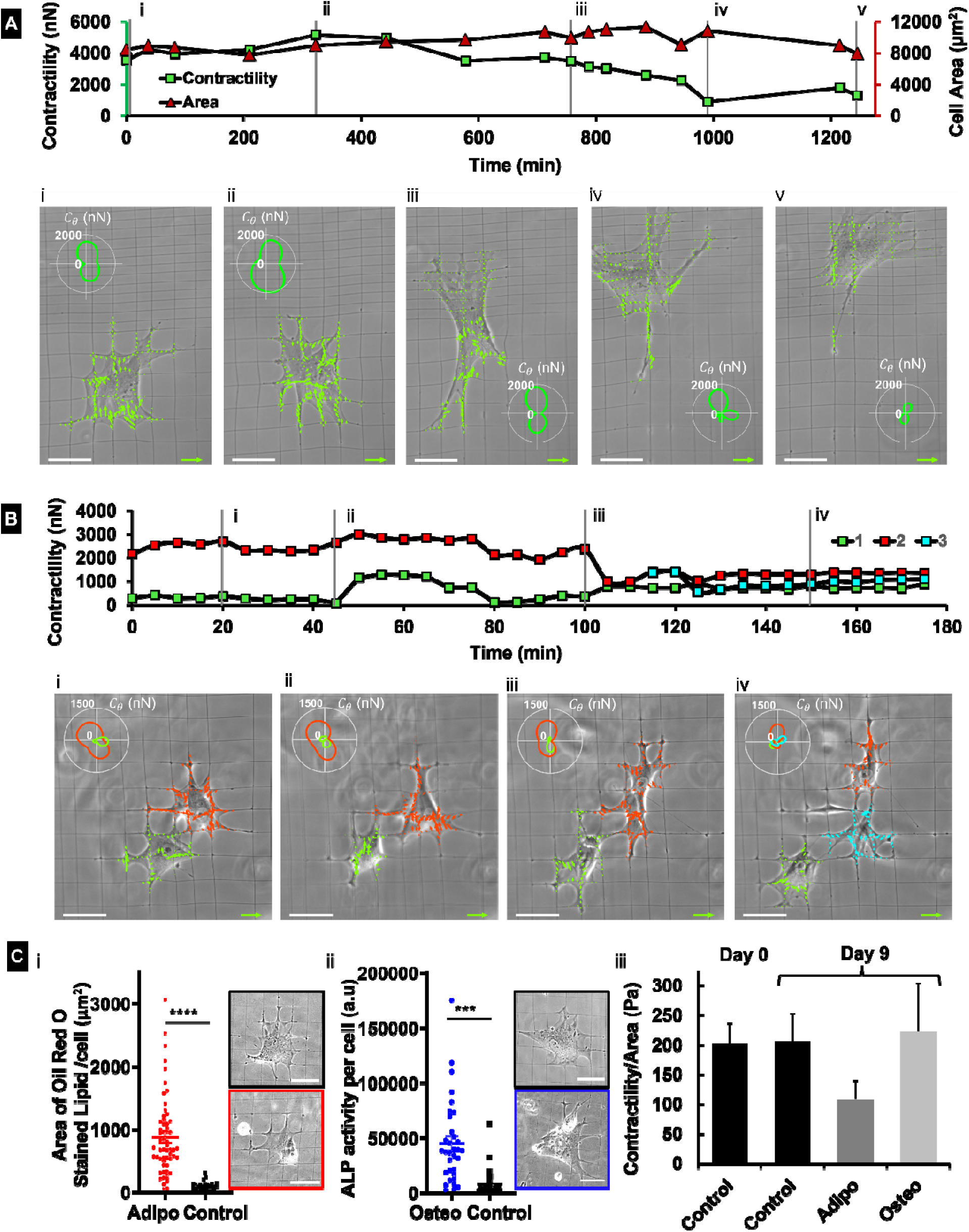
Forces in shape-controlled cell migration, cell interactions, and stem cell differentiation. (A) A single cell exerts low and high contractility cycles across the migration cycle as it transitions through different shapes. Line tension maps are shown for times marked as i-v of the cell migrating across regions of varying fiber arrangements in various shapes. Force polarity maps *C_θ_* (insets) capture the change of force polarity with cell morphology and motion. (B) Time evolution of contractility of two interacting cells and line tension maps for selected frames i-iv. Force polarity maps *C_θ_* (insets) depict the evolution of the force patterns of interacting cells along the direction of respective cell motion. (C) i. The total area covered by Oil Red O stained lipid droplets quantified for cells treated with adipogenic medium vs growth medium (control) confirms adipogenic differentiation. n=58 (N=5) for Adipogenic, n=25 (N=2) for Control. The image in the black box represents a cell on Day 0, and the image in the red box represents a cell on Day 9. ii. Alkaline phosphatase (ALP) activity of cells in osteogenic medium vs growth medium (control) expressed in arbitrary units indicates osteogenic differentiation. n=33 (N=4) for Osteogenic, n=32 (N=2) for Control. The image in the black box represents a cell on Day 0, and the image in the blue box represents a cell on Day 9. iii. Cells in the growth medium (control) maintain contractility magnitudes over 9 days. Cells undergoing adipogenic differentiation exhibit lower contractility than those in growth medium (control) and those undergoing osteogenic differentiation. n=15 (N=2) for Control Day-0, n=10 (N=5) for Adipogenic, n=6 (N=4) for Osteogenic and Control Day-9. n=number of cells, N=number of independent samples (2 sets of independent experiments were performed for all the data) Green tension scale arrow: (A, B) 35 mN/m. White scale bar: (A, B) 50μm.

Finally, we show how contractility affects stem cell differentiation in compliant fibrous environments. Stem cell fate is well known to be influenced by matrix elasticity ^68^ and mechanical tractions ^55,69,70^. However, the evolution of tractions during differentiation on a compliant fibrous ECM has yet to be reported. hMSCs treated with adipogenic differentiation medium for nine days (red box, **Fig. 5Ci**) reduced contractility over time, while hMSCs treated with osteogenic differentiation medium for nine days (blue box, **Fig. 5Cii**), both compared to control hMSCs that were in growth medium for nine days (**Fig. 5Ciii**). We confirmed differentiation by staining for lipid droplets (adipogenic differentiation) using Oil Red (**Fig. 5Ci**, **Supplementary Fig. 24**) and alkaline phosphatase activity (ALP, osteogenic differentiation, **Fig. 5Cii**, **Supplementary Fig. 25**). Results demonstrate our platform’s robustness to estimate single cell contractility across a range of biological conditions, including cell-cell interactions, response to drugs, migration, and differentiation.

## Discussion

Understanding how cells navigate through tissues has been limited by our inability to measure the forces they generate in fibrous environments. Here, we overcome this barrier with DLFM, which captured cellular forces in real time using only a standard microscope with phase contrast imaging. This allowed continuous monitoring of cells over extended periods without the limitations of fluorescence microscopy or the need to add potentially disruptive markers to cells. A key advantage of our approach is that we could determine the natural, undeformed state of the fiber network without paralyzing cells chemically, a requirement of traditional force measurement methods.^14,56,72^

The success of DLFM stems from two technical breakthroughs. First, our deep learning algorithms can track moving cells and the fibers they deform, a challenging computer vision problem that has long challenged the field. Second, our mathematical framework for calculating forces is robust against noise and large deformations. The forces we measure align well with previous studies using other methods, validating our approach.^29,69,73–75^

DLFM has enabled three key findings that advance our understanding of mechanobiology. First, we found that cells undergo force changes up to ten-fold as they change shape during migration. We could track these force dynamics in individual cells over time, something not possible with previous methods. Second, we discovered that stem cells show distinct patterns of force generation as they differentiate: forces drop early in fat cell formation but increase late in bone cell development. This finding further demonstrates DLFM’s ability to track unlabeled cells over many days. Third, we found that cells in fibrous environments form force-generating adhesions away from the periphery of cells. These are associated with the local transference of fiber tension to the cell cytoskeleton, demonstrating spatial resolution far exceeding the resolution of other fibrous traction systems.

### Conclusions

DLFM provides a powerful method for quantifying how cells exert forces dynamically as they move through tissue-like fibrous environments, offering new insight into cell behavior. By combining standard microscopy with deep learning, we could track living cells pulling and deforming fibers without needing fluorescent labels or other markers that might disturb their behavior. We demonstrated DLFM’s capabilities by measuring forces during key biological processes, revealing how cells adjust forces to navigate fibrous surroundings. Notably, we discovered that cells must form force-generating attachments throughout their surface to move through fiber networks, a departure from their behavior on flat surfaces. By capturing these dynamic mechanical interactions between cells and fibers, DLFM opens new possibilities for understanding cellular behavior in contexts ranging from morphogenesis to disease.

## Materials and Methods

### Crosshatch fiber network manufacturing

Suspended orthogonal fiber networks were generated without electrospinning using "spinneret-based tuned engineered polymers" (STEP)^76–78^. Briefly, nonelectrospun ∼250 nm diameter polystyrene (Scientific Polymer Products, Ontario, NY) fibers dissolved in xylene solution (Thermo Fisher Scientific, Waltham, MA) were suspended in two orthogonal layers with defined fiber spacings, typically a 25 x 25 µm square grid or a 25 x 5-8 µm rectangular grid. For GAN training data set generation, fibers were immersed in 4 µg/ml rhodamine-tagged fibronectin (Cytoskeleton Inc, CO).

### Cell culture and imaging

Human mesenchymal stem cells (hMSCs, Lonza) were cultured on fiber scaffolds in 5% CO_2_ at 37°C. Fluorescence and phase contrast imaging was performed using a Zeiss AxioObserver Z1 with a 20x objective. See **Supplementary Note 6** for details on differentiation media composition, protocols for Oil Red O, alkaline phosphatase staining, and analysis of the obtained RGB images (with camera Amscope MU300).

### Cell Staining and Imaging

Fiber solutions were dyed with BDP FL Maleimide dye (10 µl of 1 mg/ml, Lumiprobe) before spinning to get fluorescent fibers. Cells were seeded on the fiber scaffolds and were allowed to spread for 4 hours. Next, cells were fixed with 4% paraformaldehyde in PBS (Santa Cruz Chemicals) for 15 min. The cells were washed with PBS two times and then permeabilized with 0.1% Triton X -100 solution (Fisher Scientific) for 15 mins. Following two PBS washes, the cells were blocked with 5 % goat serum (Fisher Scientific) for 30 mins. The primary mouse monoclonal antibody for paxillin (Thermo Fisher) was diluted (1:100) in an antibody dilution buffer (0.1% BSA in PBS, Cytiva) and was added to the cells. Cells were incubated for 24 hours at 4°C. The following day, the cells were washed with PBS (3x, 5 mins each). Next, conjugated antibody Alexa Fluor 568 Phalloidin (1:100, Thermo Fisher) and secondary antibody Alexa Fluor 647 goat anti-mouse (Thermo Fisher) were diluted in the antibody dilution buffer and added to the cells. After one hour, the cells were washed with PBS (3x, 5 mins each). The sample was then covered in 2 mL of PBS for imaging. Cells were imaged using a Nikon AX confocal microscope with a 20x objective. Z-stacks were acquired at a step size of 0.75 μm. Images were processed in Nikon’s NIS elements software.

### Phase to fluorescent image translation using pix2pix

We used a pix2pix GAN model to translate the phase contrast images to fluorescent images. Details of the pix2pix model are provided in **Supplementary Note 3**. Our pix2pix model was trained on a pristine phase and fluorescent image pair set (**Supplementary Note 3**).

### Node detection and tracking

We used *RetinaNet* to detect grid intersections from the synthetically generated fluorescent images. Additional details of *RetinaNet* are provided in **Supplementary Note 3**. We further developed a bi-directional tracking algorithm to recover nodes that the RetinaNet did not detec*t*. Specifically, the tracking algorithm utilized information from the previous and future frames to recover nodes at a current frame. More details on the bi-directional tracking algorithm and the training and testing setups of node detection and tracking are provided in **Supplementary Note 3**.

### Finite element (FE) based inverse formulation

Nanofibers are modeled as beams undergoing elastic deformations with pretension arising from manufacturing ^28^. For the FE model of nanofibers mechanics, we used corotational beam formulation ^79^, which accounts for the geometrical nonlinearities due to large rigid body motions and small elastic strains in a corotating frame. The inverse problem is formulated as a regularization-based minimization problem constrained by the nanonet mechanics. The formulation is, therefore, similar in structure to classical traction force microscopy on soft gels ^62,80^. The inverse problem solver attempts to minimize the difference between measured and FE model deformations by tweaking the unknown nodal forces. The difference is quantitated using an error function with two contributions: (1) difference between nodal displacements at grid intersections in measured data and the FE model, and (2) misalignment between deformed fiber shapes in artificial fluorescent images and the FE model, similar to the active-contour segmentation approach ^81,82^.

Furthermore, a regularization term penalizes the magnitude of nodal forces to avoid overfitting the noise in measurements. We perform the minimization of the objective function (error + regularization) using the Broyden-Fletcher-Goldfarb-Shanno (BFGS) algorithm ^83^, where the gradient of the nonlinear implicitly-constrained minimization problem is obtained by using the adjoint-set method ^64–66^. The details of our inverse formulation, finite elemental model, and validation are provided in **Supplementary Note 5**.

### Contractility and force polarity maps

The estimated force pattern **f** is obtained as a set of nodal forces on the computational finite elemental mesh. To quantitate **f**, we extract the overall cell contractility as a biophysical measure of cell mechanics defined as ^62^

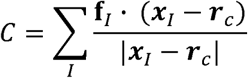

where **f**_*I*_ is the estimated force at node *I*, ***x***_*I*_ is the position vector of that node, and ***r***_*c*_ is the epicenter of the force pattern. The epicenter of the force pattern is defined as a point that minimizes the sum of cross-products of nodal force vectors with the position vector from the epicenter such that ***r***_*c*_ = argmin *W_c_* (***r***), where *W_c_* (***r***) = ∑_*I*_ |**f**_*I*_ × (***x***_*I*_ − ***r***)|.

Furthermore, the orientational organization of the force pattern is examined by evaluating the polar distribution of cellular contractility using

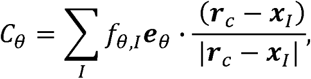

where ***e***_*θ*_ is a unit vector at an angle *θ* to the *x*-axis and *f_θ,I_* = min(**f**_*I*_ · ***e***_*θ*_, 0). For a contractile force pattern, *C_θ_* quantifies the contractility along e_*θ*_. The force polarity map is then obtained by plotting *C_θ_* in polar coordinates (Fig. 4C-E iv). This map visually reveals any preferential alignment of contractile forces along a given direction.

### Cell model

The computational model used in this study is a 3D finite element model ^26,45,84^. Details of the model can be found in **Supplementary Notes 1-2**.

The cell cytoskeleton in this model is represented as a continuous structure composed of representative volume elements (RVEs), each of which encompasses (i) the myosin molecular motors, (ii) the actin filament network, and (iii) the microtubule network. The interconnections between these elements are shown in **Supplementary Fig. 1**. The myosin element functions as an active agent, generating internal contractile forces whose magnitude increases in response to tension within the cytoskeleton. This increase in force occurs concurrently with the stiffening of the actin element in tension. These combined mechanisms collectively represent the promotion of cell actomyosin in response to tension.

Focal adhesions are modeled as a thin, initially soft layer, which undergoes local stiffening when tensile forces at a particular location exceed a specific threshold. This transformation establishes a physical link between the cell and the ECM at that location, representing the formation of mature focal adhesions. Treating focal adhesions as a thin 3D layer allows for generating both in-plane and out-of-plane stresses at the cell-fiber interface induced by cell contractility. Our simulations demonstrated that the out-of-plane component of these tensile stresses regulates the formation of focal adhesions observed away from the cell periphery in our experiments. This was confirmed by restricting the stresses generated at the cell-fiber interface to only in-plane stresses in our simulations, achieved by treating the focal adhesion as a 2D layer without thickness in the z-direction. This resulted in the formation of focal adhesions limited to the cell periphery (**Supplementary Fig. 2**), failing to replicate the experimentally observed focal adhesions within the cell body.

Our simulations were performed with a symmetry condition relative to the X-Y plane, reflecting our experimental findings that cells wrap around fibers ^67,85,86^. This wrapping resulted in a nearly symmetrical distribution of the cell cytoskeleton both above (positive Z-direction) and below (negative Z-direction) the fiber network. As a result, the out-of-plane contractility of cells generated an almost symmetrical stress field with respect to the X-Y plane in our simulations. Therefore, although the magnitude of these out-of-plane stresses can be significant, their symmetric distribution minimized the out-of-plane deformations of ECM fibers. This was consistent with our experimental results, where we observed that ECM fiber deformations primarily occurred in-plane, with instances of significant non-planar deformations being rare.

### Statistical analysis

Statistical analysis was performed using GraphPad Prism (GraphPad Software, La Jolla, CA, USA) software. Statistical comparisons among multiple groups were performed using one-way ANOVA and Tukey’s honest significant difference test. Pairwise statistical comparisons were performed using Student’s t-test. Error bars in scatter data plots indicate standard deviation. *,**,***,**** represent p< 0.05, 0.01, 0.001 and 0.0001 respectively.

## Acknowledgments

ASN, SK, and AK acknowledge funding support from the National Science Foundation (NSF, Grant No. 2107332). ASN acknowledges partial funding from the National Science Foundation (NSF, Grant No. 2422340 and 2119949) and the National Institute of Health (1R01 HL162822-01A1). ASN acknowledges the Institute of Critical Technologies and Science (ICTAS) and Macromolecules Innovative Institute (MII) at Virginia Tech for supporting this study. FA acknowledges startup funding provided by the NJIT. GMG acknowledges support from the National Institutes of Health (grants R01DK131177, R01AR077793, R01HL159094) and by the National Science Foundation through grants CMMI 1548571 (the NSF Science and Technology Center for Engineering Mechanobiology), DMR 2105150, and OIA-2219142.

## List of Supplementary Movies

1. **<u>Supplementary Movie M1</u>**: **Tension-dependent formation of actin fibers and focal adhesions.** Prediction of actin fiber formation (left panel), mature focal adhesion formation (middle panel), and extracellular fiber deformation by the computational cell model.
2. **<u>Supplementary Movie M2</u>**: **Significant fiber deformations by contractile migrating hMSCs** Time lapse of phase contrast images, the fluorescent images of rhodamine tagged fibers and the corresponding pix2pix generated images, showing migrating human mesenchymal stem cells (hMSCs) significantly deforming fibers under contractile forces.
3. **<u>Supplementary Movie M3</u>**: **Cellular forces resolved by crosshatch nanonet force microscopy** Time lapse of phase images (Supplementary Movie M1), pix2pix model output, and line tension map obtained by solving the inverse problem. Scale bar: white bar: 50µm green arrow: 35mN/m.
4. **<u>Supplementary Movie M4</u>**: **Trained pix2pix model output** Time lapse of migrating hMSCs on deforming fibers showing the phase-contrast image (left), experimentally obtained fluorescent images or ground truth (center), and the output from trained pix2pix model (right). Pix2pix-generated images are devoid of the aberrations observed in the ground truth fluorescent images.
5. **<u>Supplementary Movie M5</u>**: **Trained pix2pix model output** Time lapse of migrating hMSCs on deforming fibers showing the phase-contrast image (left), experimentally obtained fluorescent images or ground truth (center), and the output from trained pix2pix model (right). Pix2pix-generated images are devoid of the aberrations observed in the ground truth fluorescent images.
6. **<u>Supplementary Movie M6</u>**: **Actin-dependent force dynamics** Line tension map and corresponding force polarity maps depicting higher contractility of cells in control condition and significantly reduced contractility when treated with actin inhibitor latrunculin B. Scale bar: white bar: 50µm green arrow: 35mN/m
7. **<u>Supplementary Movie M7</u>**: **ROCK pathway dictates cellular contractile dynamics** Line tension map and corresponding force polarity maps depicting higher contractility of cells in control condition and significantly reduced contractility when treated with ROCK inhibitor Y27632. Contractility is regained when the drug is washed off. Scale bar: white bar: 50µm green arrow: 35mN/m
8. **<u>Supplementary Movie M8</u>**: **Rectangular fiber arrangement causes cellular anisotropy** Line tension map and corresponding force polarity maps shows anisotropic cell shapes and polarized migration along rectangular crosshatch fiber arrangements Scale bar: white bar: 50µm green arrow: 35mN/m
9. **<u>Supplementary Movie M9</u>**: **Hybrid fiber architecture dictates varied cellular shape and contractility** Line tension map and corresponding force polarity maps shows varied cell shapes and cycles of low and high contractility as the cell migrates from a sparse region to a denser arrangement of fibers, accompanied by a drastic drop in the contractility by an order of magnitude. Scale bar: white bar: 50µm green arrow: 35mN/m
10. **<u>Supplementary Movie M10</u>**: **Force dynamics of multiple interacting cells** Line tension map and corresponding force polarity maps shows two initially interacting cells and the emergence of a third interacting cell through mitosis as they all move apart. Scale bar: white bar: 50µm green arrow: 35mN/m

