## Supplementary information for "Deep Learning Reveals How Cells Pull, Buckle, and Navigate Tissue-Like Environments"

### Supplementary Information for ‘Deep learning enables real-time cell force measurement in fibrous matrices’

### equal first authors

[1]Department of Mechanical Engineering, Virginia Tech, Blacksburg, VA 24061, USA
[2]Department of Computer Science, Virginia Tech, Blacksburg, VA 24061, USA
[3]Department of Mechanical and Industrial Engineering, New Jersey Institute of Technology, Newark, NJ 07102, USA
[4] NSF Science and Technology Center for Engineering Mechanobiology, and Department of Mechanical Engineering & Materials Science, Washington University in St. Louis, St. Louis, MO 63130
[5]Center of Soft Matter and Biological Physics, Virginia Tech, Blacksburg, VA 24061, USA

### 1 Three-dimensional cytoskeleton model

The cytoskeleton model includes three elements; the myosin molecular motors, the microtubules, and the actin filaments (Supplementary Figure 1).[1]

#### 1.1 Cytoskeletal components

The myosin element in our model generates cell internal contractility. This contractility is denoted by a tensor, $\rho_{ij}$, whose components are cell-generated stress in different directions.[2] The cell contractility $\rho_{ij}$ in our model changes in response to tension generated in the cytoskeleton. Below, we describe the contractility tensor $\rho_{ij}$ and we show how cell contractility in our model changes with tension.

The contractile force exerted by an individual myosin motor can be modeled as a force dipole which is a pair of equal forces $F_i(x_j)$ and $-F_i(x_j + \Delta x_j)$ directed oppositely, where $x_j$ and $x_j + \Delta x_j$ are the coordinates of myosin head domains, $|\Delta x_j|$ is the length of myosin filament, and $|F_i|$ is the magnitude of the force dipole.[3] The total mechanical work done by the force dipoles (all myosin motors) per volume is calculated as follows

$$W = \left(\frac{1}{V}\right)\sum_{k=1}^{N}\left[F_i^{(k)}u_i\left(x_j^{(k)} + \Delta x_j^{(k)}\right) - F_i^{(k)}u_i\left(x_j^{(k)}\right)\right] \tag{1.1}$$

where $u_i(x_j)$ and $u_i(x_j + \Delta x_j)$ are the cytoskeletal displacements at myosin head domains $x_j$ and $x_j + \Delta x_j$, respectively. Note that $V$ is the volume of the cytoplasm, and $N$ is the number of phosphorylated myosin motors. Using the following equation

$$\partial_j u_i = \frac{u_i\left(x_j^{(k)} + \Delta x_j^{(k)}\right) - u_i\left(x_j^{(k)}\right)}{\Delta x_j^{(k)}} \tag{1.2}$$

we can rewrite the total work per volume in Eq. (1.1) as follows

$$W = \left(\frac{1}{V}\right)\sum_{k=1}^{N} F_i^{(k)}\,\Delta x_j^{(k)}\partial_j u_i \tag{1.3}$$

The total work in Eq. (1.3) can be represented as a function of the contractility tensor $\rho_{ij}$ and the strain tensor $\varepsilon_{ij}$:

$$W = \rho_{ij}\varepsilon_{ij} \tag{1.4}$$

where $\rho_{ij}$ is the following symmetric tensor

$$\rho_{ij} = \left(\frac{1}{V}\right)\sum_{k=1}^{N} F_i^{(k)}\,\Delta x_j^{(k)} \tag{1.5}$$

and $\varepsilon_{ij}$ is the linearized strain tensor defined as follows

$$\varepsilon_{ij} = \frac{1}{2}\left(\partial_j u_i + \partial_i u_j\right) \tag{1.6}$$

Note that since the contractility tensor $\rho_{ij}$ is symmetric, it holds (1.7).

$$F_i^{(k)} \Delta x_j^{(k)} = F_j^{(k)} \Delta x_i^{(k)} \tag{1.7}$$

Equation (1.5) shows that the cell contractility tensor $\rho_{ij}$ is related to the density of myosin molecular motors per unit volume. As mentioned earlier, experimental observations have shown that the cells adjust the density of phosphorylated myosin in response to mechanical signals, hence increasing the cell contractility with tension.[4,5] To capture this fact in our coarse-grained model, we assume that the average cell contractility in all directions $\frac{1}{3}\rho_{kk} = \frac{\rho_{11}+\rho_{22}+\rho_{33}}{3}$ is related to the average tension in the cytoskeleton $\frac{1}{3}\sigma_{kk} = \frac{\sigma_{11}+\sigma_{22}+\sigma_{33}}{3}$ through a positive feedback loop

$$\frac{\rho_{kk}}{3} = f_{\mathrm{m}} \frac{\sigma_{kk}}{3} + f_0\, \rho_0 \tag{1.8}$$

where $f_{\mathrm{m}}$ is the feedback constant parameter that regulates the feedback mechanism between the cell contractility and the external tension. The constant parameter $\rho_0$ is the initial cell contractility, and the constant parameter $f_0$ regulates the average contractility when cell contraction is not resisted externally ($\sigma_{kk} = 0$).

After describing the main hypothesis of the model Eq. (1.8), we proceed to outline the details of the model and determination of the total stress $\sigma_{ij}$ and total stiffness $C_{ij}$ tensors required for the implementation of the model into a 3D finite element framework. We begin by defining the cell contractility tensor $\rho_{ij}$ as a function of the compressive strain tensor $\varepsilon^{(\mathrm{X})}$ (see Supplementary Figure 1). We will later show that this definition of contractility tensor in Eq. (1.9) allows us to capture the increase in cell contractility in response to tension, as shown in Eq. (1.8).

$$\rho_{ij} = K^{(\rho)} \varepsilon_{kk}^{(\mathrm{X})} \delta_{ij} + 2\mu^{(\rho)} \left( \varepsilon_{ij}^{(\mathrm{X})} - \frac{1}{3} \varepsilon_{kk}^{(\mathrm{X})} \delta_{ij} \right) + \bar{\rho}_0 \delta_{ij} \tag{1.9}$$

where $K^{(\rho)}$ is the effective modulus for the motor density

$$K^{(\rho)} = \frac{3K^{(\mathrm{MT})}\alpha_{\mathrm{v}} - 1}{3(\beta_{\mathrm{v}} - \alpha_{\mathrm{v}})} \tag{1.10}$$

$\mu^{(\rho)}$ is the effective modulus of the polarization

$$\mu^{(\rho)} = \frac{2\mu^{(\mathrm{MT})}\alpha_{\mathrm{d}} - 1}{2(\beta_{\mathrm{d}} - \alpha_{\mathrm{d}})} \tag{1.11}$$

$\bar{\rho}_0$ is the effective contractility

$$\bar{\rho}_0 = \frac{\beta_{\mathrm{v}}\, \rho_0}{\beta_{\mathrm{v}} - \alpha_{\mathrm{v}}} \tag{1.12}$$

$K^{(\mathrm{MT})}$ is the bulk modulus of the cytoskeletal components that are in compression (e.g., the microtubule network), and

$$K^{(\mathrm{MT})} = \frac{E^{(\mathrm{MT})}}{3(1 - 2v^{(\mathrm{MT})})} \tag{1.13}$$

$\mu^{(\mathrm{MT})}$ is the shear modulus of the cytoskeletal components that are in compression,

$$\mu^{(\mathrm{MT})} = \frac{E^{(\mathrm{MT})}}{2(1 + v^{(\mathrm{MT})})} \tag{1.14}$$

with $E^{(\mathrm{MT})}$ and $v^{(\mathrm{MT})}$ denoting the elastic modulus and the Poisson's ratio of the cytoskeletal components that are in compression. Eqs. (1.9-14) introduce four chemo-mechanical parameters $\alpha_{\mathrm{v}}$, $\alpha_{\mathrm{d}}$, $\beta_{\mathrm{v}}$ and $\beta_{\mathrm{d}}$. The volumetric chemo-mechanical feedback parameter $\alpha_{\mathrm{v}}$ regulates the feedback mechanism between contractility and mechanical tension, e.g., a large value of $\alpha_{\mathrm{v}}$ leads to higher overall phosphorylated myosin molecular motors and subsequently higher contractility. $\alpha_{\mathrm{d}}$ is the deviatoric chemo-mechanical feedback parameter, representing the cell's tendency to polarized contraction, e.g., a small value of $\alpha_{\mathrm{d}}$ leads to non-polarized contractility. $\beta_{\mathrm{v}}$ is the volumetric chemical stiffness parameter and regulates the mean contractility $\frac{1}{3}\rho_{kk} = \frac{\rho_{11}+\rho_{22}+\rho_{33}}{3}$ in the absence of tension ($\sigma_{kk} = 0$) by keeping the contractility at a basal level, e.g., a large value of $\beta_{\mathrm{v}}$ keeps the level of myosin phosphorylation low. $\beta_{\mathrm{d}}$ is the deviatoric chemical stiffness parameter which represents the disinclination of myosin to orient along the actin and cell's polarization direction, e.g., high values of $\beta_{\mathrm{d}}$ results in a random orientation of myosin motors.

As cells contract, it is known that a large portion of microtubules are compressed.[6,7] This compressive load in microtubules is represented by $\mathrm{C}_{ijkl}^{(\mathrm{MT})}\varepsilon_{kl}^{(\mathrm{X})}$ in our model where $\mathbf{C}^{(\mathrm{MT})}$ is a fourth-order tensor denoting the stiffness tensor of the microtubule network

$$\mathrm{C}_{ijkl}^{(\mathrm{MT})} = K^{(\mathrm{MT})}\delta_{ij}\delta_{kl} + \mu^{(\mathrm{MT})}\left(\delta_{ik}\delta_{jk} + \delta_{il}\delta_{jk} - \frac{2}{3}\delta_{ij}\delta_{kl}\right) \tag{1.15}$$

Unlike microtubules, actin filaments are known to experience tension as cells contract. This is shown in our model by Eq. (1.16) where, in addition to compressing microtubules, the cell contractility $\rho_{ij}$ generates tension in the actin network, $\sigma_{ij}$

$$\rho_{ij} = -\mathrm{C}_{ijkl}^{(\mathrm{MT})}\varepsilon_{kl}^{(\mathrm{X})} + \sigma_{ij} \tag{1.16}$$

As described in the main text, our simulations start with a state of uniform and isotropic cell contraction, reflecting an even distribution of myosin densities throughout the cell in all spatial directions. To show that the cell contractility is initially isotropic (the same in all directions), we can rewrite Eq. (1.9) in the following form

$$\rho_{ij} = \mathrm{C}_{ijkl}^{(\rho)}\varepsilon_{kl}^{(\mathrm{X})} + \bar{\rho}_0\delta_{ij} \tag{1.17}$$

where

$$\mathrm{C}_{ijkl}^{(\rho)} = K^{(\rho)}\delta_{ij}\delta_{kl} + \mu^{(\rho)}\left(\delta_{ik}\delta_{jk} + \delta_{il}\delta_{jk} - \frac{2}{3}\delta_{ij}\delta_{kl}\right) \tag{1.18}$$

These equations demonstrate that, in the initial stress-free condition $\sigma_{ij} = 0$ (initially there is no tension), the diagonal components of $\rho_{ij}$ are all equal and non-zero $\rho_{11} = \rho_{22} = \rho_{33} \neq 0$, while the off-diagonal components are all zero $\rho_{12} = \rho_{21} = \rho_{13} = \rho_{31} = \rho_{23} = \rho_{32} = 0$. This indicates that the contractility tensor $\rho_{ij}$ is initially isotropic. This is better shown by writing equation (1.17) in the following format using Voigt notation

$$\begin{Bmatrix}\rho_{11}\\ \rho_{22}\\ \rho_{33}\\ \rho_{12}\\ \rho_{13}\\ \rho_{23}\end{Bmatrix} = \begin{bmatrix} C^{(\rho)}_{1111} & C^{(\rho)}_{1122} & C^{(\rho)}_{1133} & C^{(\rho)}_{1112} & C^{(\rho)}_{1113} & C^{(\rho)}_{1123}\\ C^{(\rho)}_{2211} & C^{(\rho)}_{2222} & C^{(\rho)}_{2233} & C^{(\rho)}_{2212} & C^{(\rho)}_{2213} & C^{(\rho)}_{2223}\\ C^{(\rho)}_{3311} & C^{(\rho)}_{3322} & C^{(\rho)}_{3333} & C^{(\rho)}_{3312} & C^{(\rho)}_{3313} & C^{(\rho)}_{3323}\\ C^{(\rho)}_{1211} & C^{(\rho)}_{1222} & C^{(\rho)}_{1233} & C^{(\rho)}_{1212} & C^{(\rho)}_{1213} & C^{(\rho)}_{1223}\\ C^{(\rho)}_{1311} & C^{(\rho)}_{1322} & C^{(\rho)}_{1333} & C^{(\rho)}_{1312} & C^{(\rho)}_{1313} & C^{(\rho)}_{1323}\\ C^{(\rho)}_{2311} & C^{(\rho)}_{2322} & C^{(\rho)}_{2333} & C^{(\rho)}_{2312} & C^{(\rho)}_{2313} & C^{(\rho)}_{2323}\end{bmatrix} \begin{Bmatrix}\varepsilon_{11}\\ \varepsilon_{22}\\ \varepsilon_{33}\\ \varepsilon_{12}\\ \varepsilon_{13}\\ \varepsilon_{23}\end{Bmatrix} + \begin{Bmatrix}\bar{\rho}_0\\ \bar{\rho}_0\\ \bar{\rho}_0\\ 0\\ 0\\ 0\end{Bmatrix} \tag{1.19}$$

where the first, second, and third components of the last term in (1.19) have the same value $\bar{\rho}_0$. Note that combining Eqs. (1.16) with (1.17), one can derive the following expression,

$$\frac{\rho_{kk}}{3} = \left(\frac{3K^{(\mathrm{MT})}\alpha_{\mathrm{v}} - 1}{3K^{(\mathrm{MT})}\beta_{\mathrm{v}} - 1}\right)\frac{\sigma_{kk}}{3} + \left(\frac{3K^{(\mathrm{MT})}\beta_{\mathrm{v}}}{3K^{(\mathrm{MT})}\beta_{\mathrm{v}} - 1}\right)\rho_0$$

which is the expanded version of Eq. (1.8) and demonstrates the relationship between cell contractility and tension as the main hypothesis of our model.

Concomitant with the phosphorylation of more myosin, cells also respond to tension by polymerizing actin filaments and bundling and aligning them in the direction of tension.[8,9] Thus, in addition to the feedback mechanism between the contractility $\rho_{ij}$ and the stress $\sigma_{ij}$ in (1.8), we also hypothesize that the stiffness of the actin network $C^{(A)}_{ijkl}$ increases in proportion to, and in the direction of, the tensile principal components of the stress tensor $\sigma_{ij}$,[1]

$$C^{(A)}_{ijkl} = C^{(I)}_{ijkl} + C^{(F)}_{ijkl} \tag{1.20}$$

In the above equation, $\mathbf{C}^{(\mathrm{F})}$ represents the stiffening of the actin network with tension (but not in compression), while $\mathbf{C}^{(\mathrm{I})}$ is the initial stiffness of the actin filaments network

$$C^{(I)}_{ijkl} = K^{(\mathrm{I})}\delta_{ij}\delta_{kl} + \mu^{(\mathrm{I})}\left(\delta_{ik}\delta_{jk} + \delta_{il}\delta_{jk} - \frac{2}{3}\delta_{ij}\delta_{kl}\right) \tag{1.21}$$

where

$$K^{(\mathrm{I})} = \frac{E^{(\mathrm{I})}}{3(1 - 2v^{(\mathrm{I})})} \tag{1.22}$$

is the initial bulk modulus of the actin network, and

$$\mu^{(\mathrm{I})} = \frac{E^{(\mathrm{I})}}{2(1 + v^{(\mathrm{I})})} \tag{1.23}$$

is the initial shear modulus of the actin network. $E^{(\mathrm{I})}$ and $v^{(\mathrm{I})}$ are the initial elastic modulus and Poisson's ratio of the actin network.

To define the stiffening part of the stiffness tensor, $\mathbf{C}^{(\mathrm{F})}$, we first decompose $\boldsymbol{\sigma}$

$$\sigma_{ij} = \sigma^{(A)}_{ij} = \sigma^{(I)}_{ij} + \sigma^{(F)}_{ij} \tag{1.24}$$

where $\boldsymbol{\sigma}^{(\mathrm{I})}$

$$\sigma_{ij}^{(\mathrm{I})} = \mathrm{C}_{ijkl}^{(\mathrm{I})}\, \varepsilon_{kl}^{(\mathrm{Y})} \tag{1.25}$$

is linearly related to the strain tensor $\boldsymbol{\varepsilon}^{(\mathrm{Y})}$ (the three-dimensional representation of $\varepsilon^{(\mathrm{Y})}$ shown in Supplementary Figure 1) which can be written as a function of its eigenvalues (principal strains) $\varepsilon_1^{(\mathrm{Y})}$, $\varepsilon_2^{(\mathrm{Y})}$, $\varepsilon_3^{(\mathrm{Y})}$ and eigenvectors $\boldsymbol{n}_1$, $\boldsymbol{n}_2$, $\boldsymbol{n}_3$

$$\boldsymbol{\varepsilon}^{(\mathrm{Y})} = \sum_{i=1}^{3} \varepsilon_i^{(\mathrm{Y})}\, \boldsymbol{n}_i \otimes \boldsymbol{n}_i = \sum_{i=1}^{3} \varepsilon_i^{(\mathrm{Y})}\, \mathbf{E}_i \tag{1.26}$$

where the symmetric tensors $\mathbf{E}_1 = \boldsymbol{n}_1 \otimes \boldsymbol{n}_1$, $\mathbf{E}_2 = \boldsymbol{n}_2 \otimes \boldsymbol{n}_2$, and $\mathbf{E}_3 = \boldsymbol{n}_3 \otimes \boldsymbol{n}_3$ are the eigenprojections of $\boldsymbol{\varepsilon}^{(\mathrm{Y})}$ and $\otimes$ denotes the dyadic product of two arbitrary vectors $\boldsymbol{u}$ and $\boldsymbol{v}$ as $(\boldsymbol{u} \otimes \boldsymbol{v})_{ij} = u_i v_j$. With the eigenvalues ($\varepsilon_1^{(\mathrm{Y})}$, $\varepsilon_2^{(\mathrm{Y})}$, $\varepsilon_3^{(\mathrm{Y})}$) and eigenvectors ($\boldsymbol{n}_1$, $\boldsymbol{n}_2$, $\boldsymbol{n}_3$) at hand, we next define $\sigma_{ij}^{(\mathrm{F})}$ in (1.24)

$$\boldsymbol{\sigma}^{(\mathrm{F})} = \sum_{i=1}^{3} \frac{\partial f\left(\varepsilon_i^{(\mathrm{Y})}\right)}{\partial \varepsilon_i^{(\mathrm{Y})}} \boldsymbol{n}_i \otimes \boldsymbol{n}_i = \sum_{i=1}^{3} \sigma^{(\mathrm{F})}\left(\varepsilon_i^{(\mathrm{Y})}\right) \mathbf{E}_i = \sum_{i=1}^{3} \sigma_i^{(\mathrm{F})} \mathbf{E}_i \tag{1.27}$$

where $\sigma_i^{(\mathrm{F})}$ are the eigenvalues (principal stresses) of the stress tensor $\sigma_{ij}^{(\mathrm{F})}$. We define these principal stresses as follows

$$\sigma_i^{(\mathrm{F})} = \frac{\partial f\left(\varepsilon_i^{(\mathrm{Y})}\right)}{\partial \varepsilon_i^{(\mathrm{Y})}} = \tag{1.28}$$

$$\begin{cases} 0 & \varepsilon_i^{(\mathrm{Y})} < \epsilon_1 \\ \ell \dfrac{\left(\dfrac{\varepsilon_i^{(\mathrm{Y})} - \epsilon_1}{\epsilon_2 - \epsilon_1}\right)^t \left(\varepsilon_i^{(\mathrm{Y})} - \epsilon_1\right)^2}{(t+1)(t+2)} & \epsilon_1 \le \varepsilon_i^{(\mathrm{Y})} < \epsilon_2 \\ \ell \left[\dfrac{\left(1 + \varepsilon_i^{(\mathrm{Y})} - \epsilon_2\right)^{s+2} - 1}{(s+1)(s+2)} + \dfrac{\epsilon_2 - \varepsilon_i^{(\mathrm{Y})}}{s+1} + \dfrac{\left(\varepsilon_i^{(\mathrm{Y})} - \epsilon_2\right)(\epsilon_2 - \epsilon_1)}{t+1} + \dfrac{(\epsilon_2 - \epsilon_1)^2}{(t+1)(t+2)}\right] & \varepsilon_i^{(\mathrm{Y})} \ge \epsilon_2 \end{cases}$$

The above stress-strain relationship ensures the continuity and smoothness of the first and second derivatives of $\sigma_i^{(\mathrm{F})}$ with respect to $\varepsilon_i^{(\mathrm{Y})}$ at the transition points $\epsilon_1 = \epsilon_\mathrm{c} - 0.5\epsilon_\mathrm{t}$ and $\epsilon_2 = \epsilon_\mathrm{c} + 0.5\epsilon_\mathrm{t}$, where $\epsilon_\mathrm{t} = 0.25\epsilon_\mathrm{c}$ is the transition width, $\epsilon_\mathrm{c}$ is the critical (tensile) principal strain, and $t$ is the transition constant. Eq. (1.28) shows that for large tensile strains $\varepsilon_i^{(\mathrm{Y})} \ge \epsilon_2$, the principal stress $\sigma_i^{(\mathrm{F})}$ nonlinearly increases with the principal strain $\varepsilon_i^{(\mathrm{Y})}$ where this increase is regulated by the stiffening parameters $\ell$ and $s$. With $\boldsymbol{\sigma}^{(\mathrm{F})}$ at hand from Eq. (1.29), we can now determine $\mathbf{C}^{(\mathrm{F})}$ as follows

$$\mathrm{C}^{(\mathrm{F})}_{ijkl} = \frac{\mathrm{d}\sigma^{(\mathrm{F})}_{ij}}{\mathrm{d}\varepsilon^{(\mathrm{Y})}_{kl}} \qquad \text{or} \qquad \mathbf{C}^{(\mathrm{F})} = \frac{\mathrm{d}\boldsymbol{\sigma}^{(\mathrm{F})}}{\mathrm{d}\boldsymbol{\varepsilon}^{(\mathrm{Y})}} \tag{1.29}$$

The piecewise linear approximation in (1.29) requires $\sigma^{(\mathrm{F})}_{ij}$ as a function of $\varepsilon^{(\mathrm{Y})}_{ij}$ while Eq. (1.28) gives $\sigma^{(\mathrm{F})}_{i}$ as a function of $\varepsilon^{(\mathrm{Y})}_{i}$. Thus, we first write $\mathrm{C}^{(\mathrm{F})}_{ijkl}$ in the following form using the definition of $\sigma^{(\mathrm{F})}_{ij}$ in (1.27)

$$\mathbf{C}^{(\mathrm{F})} = \sum_{i=1}^{3}\left\{\mathbf{E}_i \otimes \frac{\mathrm{d}\sigma^{(\mathrm{F})}_{i}}{\mathrm{d}\boldsymbol{\varepsilon}^{(\mathrm{Y})}} + \sigma^{(\mathrm{F})}_{i}\frac{\mathrm{d}\mathbf{E}_i}{\mathrm{d}\boldsymbol{\varepsilon}^{(\mathrm{Y})}}\right\} \tag{1.30}$$

and we then expand the first term on the right-hand side of (1.30) by applying the chain rule

$$\mathbf{C}^{(\mathrm{F})} = \sum_{i=1}^{3}\left\{\sum_{j=1}^{3}\frac{\partial\sigma^{(\mathrm{F})}_{i}}{\partial\varepsilon^{(\mathrm{Y})}_{j}}\mathbf{E}_i \otimes \frac{\mathrm{d}\varepsilon^{(\mathrm{Y})}_{j}}{\mathrm{d}\boldsymbol{\varepsilon}^{(\mathrm{Y})}} + \sigma^{(\mathrm{F})}_{i}\frac{\mathrm{d}\mathbf{E}_i}{\mathrm{d}\boldsymbol{\varepsilon}^{(\mathrm{Y})}}\right\} \tag{1.31}$$

To further expand equation (1.31) and derive an exact form for $\mathbf{C}^{(\mathrm{F})}$, we consider the three following cases. In the first case, the three eigenvalues of the strain tensor $\varepsilon^{(\mathrm{Y})}_{ij}$ are all nonidentical ($\varepsilon^{(\mathrm{Y})}_{1} \neq \varepsilon^{(\mathrm{Y})}_{2} \neq \varepsilon^{(\mathrm{Y})}_{3}$). In this case, we can derive the following expression for $\mathbf{C}^{(\mathrm{F})}$ from Eq. (1.31) by taking the derivatives of $\varepsilon^{(\mathrm{Y})}_{j}$ and $\mathbf{E}_i$ with respect to $\boldsymbol{\varepsilon}^{(\mathrm{Y})}$

$$\begin{aligned}\mathbf{C}^{(\mathrm{F})} = \sum_{a=1}^{3}\frac{\sigma^{(\mathrm{F})}_{a}}{\left(\varepsilon^{(\mathrm{Y})}_{a}-\varepsilon^{(\mathrm{Y})}_{b}\right)\left(\varepsilon^{(\mathrm{Y})}_{a}-\varepsilon^{(\mathrm{Y})}_{c}\right)}&\left\{\frac{\mathrm{d}\left(\boldsymbol{\varepsilon}^{(\mathrm{Y})}\right)^2}{\mathrm{d}\boldsymbol{\varepsilon}^{(\mathrm{Y})}} - \left(\varepsilon^{(\mathrm{Y})}_{b}+\varepsilon^{(\mathrm{Y})}_{c}\right)\mathbf{I}_{\mathrm{S}}\right.\\ &- \left[\left(\varepsilon^{(\mathrm{Y})}_{a}-\varepsilon^{(\mathrm{Y})}_{b}\right)+\left(\varepsilon^{(\mathrm{Y})}_{a}-\varepsilon^{(\mathrm{Y})}_{c}\right)\right]\mathbf{E}_a\otimes\mathbf{E}_a\\ &\left.- \left(\varepsilon^{(\mathrm{Y})}_{b}-\varepsilon^{(\mathrm{Y})}_{c}\right)\left(\mathbf{E}_b\otimes\mathbf{E}_b-\mathbf{E}_c\otimes\mathbf{E}_c\right)\right\} + \sum_{i=1}^{3}\sum_{j=1}^{3}\frac{\partial\sigma^{(\mathrm{F})}_{i}}{\partial\varepsilon^{(\mathrm{Y})}_{j}}\mathbf{E}_i\otimes\mathbf{E}_j\end{aligned} \tag{1.32}$$

where

$$\left(\frac{\mathrm{D}\left(\boldsymbol{\varepsilon}^{(\mathrm{Y})}\right)^2}{\mathrm{d}\boldsymbol{\varepsilon}^{(\mathrm{Y})}}\right)_{ijkl} = \frac{1}{2}\left(\delta_{ik}\varepsilon^{(\mathrm{Y})}_{lj} + \delta_{il}\varepsilon^{(\mathrm{Y})}_{kj} + \delta_{jl}\varepsilon^{(\mathrm{Y})}_{ik} + \delta_{kj}\varepsilon^{(\mathrm{Y})}_{il}\right) \tag{1.33}$$

is the derivative of the square of $\boldsymbol{\varepsilon}^{(\mathrm{Y})}$, and

$$(\mathbf{I}_{\mathrm{S}})_{ijkl} = \frac{1}{2}\left(\delta_{ik}\delta_{jl} + \delta_{il}\delta_{jk}\right) \tag{1.34}$$

is the symmetric identity tensor.

In the second case, $\varepsilon^{(\mathrm{Y})}_{ij}$ has two identical eigenvalues ($\varepsilon^{(\mathrm{Y})}_{1} \neq \varepsilon^{(\mathrm{Y})}_{2} = \varepsilon^{(\mathrm{Y})}_{3}$) which gives the following analytical expression for $\mathbf{C}^{(\mathrm{F})}$

$$\mathbf{C}^{(\mathrm{F})} = s_1 \frac{\mathrm{d}\left(\boldsymbol{\varepsilon}^{(\mathrm{Y})}\right)^2}{\mathrm{d}\boldsymbol{\varepsilon}^{(\mathrm{Y})}} - s_2 \mathbf{I}_\mathrm{S} - s_3 \boldsymbol{\varepsilon}^{(\mathrm{Y})} \otimes \boldsymbol{\varepsilon}^{(\mathrm{Y})} + s_4 \boldsymbol{\varepsilon}^{(\mathrm{Y})} \otimes \mathbf{I} + s_5 \mathbf{I} \otimes \boldsymbol{\varepsilon}^{(\mathrm{Y})} - s_6 \mathbf{I} \otimes \mathbf{I} \tag{1.35}$$

where $\mathrm{I}_{ij} = \delta_{ij}$ is the second-order identity tensor, and

$$\begin{aligned}
s_1 &= \frac{\sigma_a^{(\mathrm{F})} - \sigma_c^{(\mathrm{F})}}{\left(\varepsilon_a^{(\mathrm{Y})} - \varepsilon_c^{(\mathrm{Y})}\right)^2} + \frac{1}{\varepsilon_a^{(\mathrm{Y})} - \varepsilon_c^{(\mathrm{Y})}} \left( \frac{\partial \sigma_c^{(\mathrm{F})}}{\partial \varepsilon_b^{(\mathrm{Y})}} - \frac{\partial \sigma_c^{(\mathrm{F})}}{\partial \varepsilon_c^{(\mathrm{Y})}} \right) \\
s_2 &= 2\varepsilon_c^{(\mathrm{Y})} \frac{\sigma_a^{(\mathrm{F})} - \sigma_c^{(\mathrm{F})}}{\left(\varepsilon_a^{(\mathrm{Y})} - \varepsilon_c^{(\mathrm{Y})}\right)^2} + \frac{\varepsilon_a^{(\mathrm{Y})} + \varepsilon_c^{(\mathrm{Y})}}{\varepsilon_a^{(\mathrm{Y})} - \varepsilon_c^{(\mathrm{Y})}} \left( \frac{\partial \sigma_c^{(\mathrm{F})}}{\partial \varepsilon_b^{(\mathrm{Y})}} - \frac{\partial \sigma_c^{(\mathrm{F})}}{\partial \varepsilon_c^{(\mathrm{Y})}} \right) \\
s_3 &= 2 \frac{\sigma_a^{(\mathrm{F})} - \sigma_c^{(\mathrm{F})}}{\left(\varepsilon_a^{(\mathrm{Y})} - \varepsilon_c^{(\mathrm{Y})}\right)^3} + \frac{1}{\left(\varepsilon_a^{(\mathrm{Y})} - \varepsilon_c^{(\mathrm{Y})}\right)^2} \left( \frac{\partial \sigma_a^{(\mathrm{F})}}{\partial \varepsilon_c^{(\mathrm{Y})}} + \frac{\partial \sigma_c^{(\mathrm{F})}}{\partial \varepsilon_a^{(\mathrm{Y})}} - \frac{\partial \sigma_a^{(\mathrm{F})}}{\partial \varepsilon_a^{(\mathrm{Y})}} - \frac{\partial \sigma_c^{(\mathrm{F})}}{\partial \varepsilon_c^{(\mathrm{Y})}} \right) \\
s_4 &= 2\varepsilon_c^{(\mathrm{Y})} \frac{\sigma_a^{(\mathrm{F})} - \sigma_c^{(\mathrm{F})}}{\left(\varepsilon_a^{(\mathrm{Y})} - \varepsilon_c^{(\mathrm{Y})}\right)^3} + \frac{1}{\varepsilon_a^{(\mathrm{Y})} - \varepsilon_c^{(\mathrm{Y})}} \left( \frac{\partial \sigma_a^{(\mathrm{F})}}{\partial \varepsilon_c^{(\mathrm{Y})}} - \frac{\partial \sigma_c^{(\mathrm{F})}}{\partial \varepsilon_b^{(\mathrm{Y})}} \right) \\
&\quad + \frac{\varepsilon_c^{(\mathrm{Y})}}{\left(\varepsilon_a^{(\mathrm{Y})} - \varepsilon_c^{(\mathrm{Y})}\right)^2} \left( \frac{\partial \sigma_a^{(\mathrm{F})}}{\partial \varepsilon_c^{(\mathrm{Y})}} + \frac{\partial \sigma_c^{(\mathrm{F})}}{\partial \varepsilon_a^{(\mathrm{Y})}} - \frac{\partial \sigma_a^{(\mathrm{F})}}{\partial \varepsilon_a^{(\mathrm{Y})}} - \frac{\partial \sigma_c^{(\mathrm{F})}}{\partial \varepsilon_c^{(\mathrm{Y})}} \right) \\
s_5 &= 2\varepsilon_c^{(\mathrm{Y})} \frac{\sigma_a^{(\mathrm{F})} - \sigma_c^{(\mathrm{F})}}{\left(\varepsilon_a^{(\mathrm{Y})} - \varepsilon_c^{(\mathrm{Y})}\right)^3} + \frac{1}{\varepsilon_a^{(\mathrm{Y})} - \varepsilon_c^{(\mathrm{Y})}} \left( \frac{\partial \sigma_c^{(\mathrm{F})}}{\partial \varepsilon_a^{(\mathrm{Y})}} - \frac{\partial \sigma_c^{(\mathrm{F})}}{\partial \varepsilon_b^{(\mathrm{Y})}} \right) \\
&\quad + \frac{\varepsilon_c^{(\mathrm{Y})}}{\left(\varepsilon_a^{(\mathrm{Y})} - \varepsilon_c^{(\mathrm{Y})}\right)^2} \left( \frac{\partial \sigma_a^{(\mathrm{F})}}{\partial \varepsilon_c^{(\mathrm{Y})}} + \frac{\partial \sigma_c^{(\mathrm{F})}}{\partial \varepsilon_a^{(\mathrm{Y})}} - \frac{\partial \sigma_a^{(\mathrm{F})}}{\partial \varepsilon_a^{(\mathrm{Y})}} - \frac{\partial \sigma_c^{(\mathrm{F})}}{\partial \varepsilon_c^{(\mathrm{Y})}} \right) \\
s_6 &= 2\varepsilon_c^{(\mathrm{Y})} \frac{\sigma_a^{(\mathrm{F})} - \sigma_c^{(\mathrm{F})}}{\left(\varepsilon_a^{(\mathrm{Y})} - \varepsilon_c^{(\mathrm{Y})}\right)^3} + \frac{\varepsilon_a^{(\mathrm{Y})}\varepsilon_c^{(\mathrm{Y})}}{\left(\varepsilon_a^{(\mathrm{Y})} - \varepsilon_c^{(\mathrm{Y})}\right)^2} \left( \frac{\partial \sigma_a^{(\mathrm{F})}}{\partial \varepsilon_c^{(\mathrm{Y})}} + \frac{\partial \sigma_c^{(\mathrm{F})}}{\partial \varepsilon_a^{(\mathrm{Y})}} \right) \\
&\quad - \frac{\left(\varepsilon_c^{(\mathrm{Y})}\right)^2}{\left(\varepsilon_a^{(\mathrm{Y})} - \varepsilon_c^{(\mathrm{Y})}\right)^2} \left( \frac{\partial \sigma_a^{(\mathrm{F})}}{\partial \varepsilon_a^{(\mathrm{Y})}} + \frac{\partial \sigma_c^{(\mathrm{F})}}{\partial \varepsilon_c^{(\mathrm{Y})}} \right) - \frac{\varepsilon_a^{(\mathrm{Y})} + \varepsilon_c^{(\mathrm{Y})}}{\varepsilon_a^{(\mathrm{Y})} - \varepsilon_c^{(\mathrm{Y})}} \frac{\partial \sigma_c^{(\mathrm{F})}}{\partial \varepsilon_b^{(\mathrm{Y})}}
\end{aligned} \tag{1.36}$$

are constants with $(a, b, c)$ being cyclic permutations of (1,2,3).

In the third case, the three eigenvalues of $\varepsilon_{ij}^{(\mathrm{Y})}$ are all identical ($\varepsilon_1^{(\mathrm{Y})} = \varepsilon_2^{(\mathrm{Y})} = \varepsilon_3^{(\mathrm{Y})}$) which gives the following expression for $\mathbf{C}^{(\mathrm{F})}$

$$\mathbf{C}^{(\mathrm{F})} = \left( \frac{\partial \sigma_1^{(\mathrm{F})}}{\partial \varepsilon_1^{(\mathrm{Y})}} - \frac{\partial \sigma_1^{(\mathrm{F})}}{\partial \varepsilon_2^{(\mathrm{Y})}} \right) \mathbf{I}_\mathrm{S} + \frac{\partial \sigma_1^{(\mathrm{F})}}{\partial \varepsilon_2^{(\mathrm{Y})}} \mathbf{I} \otimes \mathbf{I} \tag{1.37}$$

Note that we need $\frac{\partial \sigma_i^{(\mathrm{F})}}{\partial \varepsilon_j^{(\mathrm{Y})}}$ for all three cases which can be determined by taking the first derivative of $\sigma_i^{(\mathrm{F})}$ in Eq. (1.29)

$$\frac{\partial \sigma_i^{(\mathrm{F})}}{\partial \varepsilon_i^{(\mathrm{Y})}} = \frac{\partial}{\partial \varepsilon_i^{(\mathrm{Y})}}\left(\frac{\partial f}{\partial \varepsilon_i^{(\mathrm{Y})}}\right) = \begin{cases} 0, & \varepsilon_i^{(\mathrm{Y})} < \epsilon_1 \\ \ell \dfrac{\left(\dfrac{\varepsilon_i^{(\mathrm{Y})} - \epsilon_1}{\epsilon_2 - \epsilon_1}\right)^t \left(\varepsilon_i^{(\mathrm{Y})} - \epsilon_1\right)}{t+1}, & \epsilon_1 \le \varepsilon_i^{(\mathrm{Y})} < \epsilon_2 \\ \ell\left[\dfrac{\left(1 + \varepsilon_i^{(\mathrm{Y})} - \epsilon_2\right)^{s+1} - 1}{s+1} + \dfrac{\epsilon_2 - \epsilon_1}{t+1}\right], & \varepsilon_i^{(\mathrm{Y})} \ge \epsilon_2 \end{cases} \tag{1.38}$$

With $\mathbf{C}^{(\mathrm{F})}$ at hand, the stiffness of the actin filament network $\mathbf{C}^{(\mathrm{A})}$ can be obtained from (1.20).

#### 1.2 Total stiffness of the cytoskeleton

To determine the total stiffness of the cytoskeleton, we first degrade the fourth-order tensors $\mathbf{C}^{(\mathrm{MT})}$ (1.15), $\mathbf{C}^{(\rho)}$ (1.18), and $\mathbf{C}^{(\mathrm{A})}$ (1.20) to the second-order tensors $\mathbf{C}^{(\mathrm{MT})}$, $\mathbf{C}^{(\rho)}$, and $\mathbf{C}^{(\mathrm{A})}$. Eq. (1.19) shows how a fourth-order tensor (*e.g.*, $\mathrm{C}_{ijkl}^{(\rho)}$) is degraded to a second-order 6×6 matrix (*e.g.*, $\mathrm{C}_{ij}^{(\rho)}$).

To determine the total stiffness tensor, one should note that the microtubule network is connected to the myosin in parallel, and they are both connected to the actin network in series (Supplementary Figure 1). Therefore, the total stiffness of the cell, $\mathbf{C}$, is obtained as follows

$$\mathbf{C} = \left(\left(\mathbf{C}^{(\mathrm{X})}\right)^{-1} + \left(\mathbf{C}^{(\mathrm{Y})}\right)^{-1}\right)^{-1} \tag{1.39}$$

where

$$\mathbf{C}^{(\mathrm{X})} = \mathbf{C}^{(\rho)} + \mathbf{C}^{(\mathrm{MT})} \tag{1.40}$$

and

$$\mathbf{C}^{(\mathrm{Y})} = \mathbf{C}^{(\mathrm{A})} = \mathbf{C}^{(\mathrm{I})} + \mathbf{C}^{(\mathrm{F})} \tag{1.41}$$

#### 1.3 Solving the set of nonlinear equations

In the previous sections, we presented the constitutive equations of the stress $\sigma_{ij}$ and the stiffness $\mathrm{C}_{ij}$ fields. However, it should be noted that the stress and stiffness tensors $\sigma_{ij}$ and $\mathrm{C}_{ij}$ are functions of the unknown strain tensors $\varepsilon_{ij}^{(\mathrm{X})}$ and $\varepsilon_{ij}^{(\mathrm{Y})}$ (and not $\varepsilon_{ij}$). To determine $\varepsilon_{ij}^{(\mathrm{X})}$ and $\varepsilon_{ij}^{(\mathrm{Y})}$, we first define the following 12×1 vector

$$\begin{aligned} \boldsymbol{u} &= \left\{\varepsilon_{11}^{(\mathrm{X})} \quad \varepsilon_{22}^{(\mathrm{X})} \quad \varepsilon_{33}^{(\mathrm{X})} \quad \varepsilon_{12}^{(\mathrm{X})} \quad \varepsilon_{13}^{(\mathrm{X})} \quad \varepsilon_{23}^{(\mathrm{X})} \quad \varepsilon_{11}^{(\mathrm{Y})} \quad \varepsilon_{22}^{(\mathrm{Y})} \quad \varepsilon_{33}^{(\mathrm{Y})} \quad \varepsilon_{12}^{(\mathrm{Y})} \quad \varepsilon_{13}^{(\mathrm{Y})} \quad \varepsilon_{23}^{(\mathrm{Y})}\right\}^{\mathrm{T}} \\ &= \{u_1 \quad u_2 \quad \cdots \quad u_{12}\}^{\mathrm{T}} \end{aligned} \tag{1.42}$$

which contains all 12 unknown variables in the strain tensors $\varepsilon_{ij}^{(\mathrm{X})}$ and $\varepsilon_{ij}^{(\mathrm{Y})}$. To determine the 12 unknowns, we need 12 equations. As the actin filament is connected to other elements in series, we can use the following condition which gives us 6 equations

$$\boldsymbol{\sigma} = \boldsymbol{\sigma}^{(\mathrm{X})} = \boldsymbol{\sigma}^{(\mathrm{Y})} \tag{1.43}$$

where the stress $\boldsymbol{\sigma}^{(\mathrm{X})}$ (Eqs. (1.16) and (1.17)) is the total stress in the parallel elements

$$\sigma_{ij}^{(\mathrm{X})} = \sigma_{ij} = \left(\mathrm{C}_{ijkl}^{(\rho)} + \mathrm{C}_{ijkl}^{(\mathrm{MT})}\right)\varepsilon_{kl}^{(\mathrm{X})} + \bar{\rho}_0\delta_{ij} \tag{1.44}$$

and $\boldsymbol{\sigma}^{(\mathrm{Y})}$ is the stress in the series element (actin element)

$$\sigma_{ij}^{(\mathrm{Y})} = \sigma_{ij} = \sigma_{ij}^{(\mathrm{A})} = \sigma_{ij}^{(\mathrm{I})} + \sigma_{ij}^{(\mathrm{F})} \tag{1.45}$$

The other 6 equations can be obtained from the following condition

$$\boldsymbol{\varepsilon} = \boldsymbol{\varepsilon}^{(\mathrm{X})} + \boldsymbol{\varepsilon}^{(\mathrm{Y})} \tag{1.46}$$

where $\boldsymbol{\varepsilon}^{(\mathrm{X})}$ is the strain of the cytoskeletal components that are in compression (e.g., microtubule network), $\boldsymbol{\varepsilon}^{(\mathrm{Y})}$ is the strain of the cytoskeletal components that are in tension (e.g., actin filaments), and $\boldsymbol{\varepsilon}$ is the total strain of the cell (Supplementary Figure 1). Note that all stress and strain tensors $\sigma_{ij}^{(\mathrm{X})}$, $\sigma_{ij}^{(\mathrm{Y})}$, $\varepsilon_{ij}^{(\mathrm{X})}$, and $\varepsilon_{ij}^{(\mathrm{Y})}$ are symmetric. Therefore, the conditions in (1.43) and (1.46) can be defined by the following 12 equations in the 12×1 vector $\boldsymbol{f}$

$$\boldsymbol{f} = \{f_1 \quad f_2 \quad \cdots \quad f_{12}\}^{\mathrm{T}} \tag{1.47}$$

where

$$f_1 = \sigma_{11}^{(\mathrm{X})} - \sigma_{11}^{(\mathrm{Y})}$$

$$f_2 = \sigma_{22}^{(\mathrm{X})} - \sigma_{22}^{(\mathrm{Y})}$$

$$f_3 = \sigma_{33}^{(\mathrm{X})} - \sigma_{33}^{(\mathrm{Y})}$$

$$f_4 = \sigma_{12}^{(\mathrm{X})} - \sigma_{12}^{(\mathrm{Y})}$$

$$f_5 = \sigma_{13}^{(\mathrm{X})} - \sigma_{13}^{(\mathrm{Y})}$$

$$f_6 = \sigma_{23}^{(\mathrm{X})} - \sigma_{23}^{(\mathrm{Y})}$$

$$f_7 = \varepsilon_{11} - \varepsilon_{11}^{(\mathrm{X})} - \varepsilon_{11}^{(\mathrm{Y})}$$

$$f_8 = \varepsilon_{22} - \varepsilon_{22}^{(\mathrm{X})} - \varepsilon_{22}^{(\mathrm{Y})}$$

$$f_9 = \varepsilon_{33} - \varepsilon_{33}^{(\mathrm{X})} - \varepsilon_{33}^{(\mathrm{Y})}$$

$$f_{10} = \varepsilon_{12} - \varepsilon_{12}^{(\mathrm{X})} - \varepsilon_{12}^{(\mathrm{Y})}$$

$f_{11} = \varepsilon_{13} - \varepsilon_{13}^{(\mathrm{X})} - \varepsilon_{13}^{(\mathrm{Y})}$

$f_{12} = \varepsilon_{23} - \varepsilon_{23}^{(\mathrm{X})} - \varepsilon_{23}^{(\mathrm{Y})}$

We next determine the 12×12 Jacobian matrix **J**

$$\mathbf{J} = \begin{bmatrix} \frac{\partial f_1}{\partial u_1} & \frac{\partial f_1}{\partial u_2} & \cdots & \frac{\partial f_1}{\partial u_{12}} \\ \frac{\partial f_2}{\partial u_1} & \frac{\partial f_2}{\partial u_2} & \cdots & \frac{\partial f_2}{\partial u_{12}} \\ \vdots & \vdots & & \vdots \\ \frac{\partial f_{12}}{\partial u_1} & \frac{\partial f_{12}}{\partial u_2} & \cdots & \frac{\partial f_{12}}{\partial u_{12}} \end{bmatrix} = \begin{bmatrix} \mathbf{C}^{(\mathrm{X})} & -\mathbf{C}^{(\mathrm{Y})} \\ -\mathbf{I} & -\mathbf{I} \end{bmatrix} \tag{1.48}$$

using equations (1.42) and (1.47) for $u_i$ and $f_i$, respectively. Finally, we use the Newton-Raphson method

$$\boldsymbol{u}_{i+1} = \boldsymbol{u}_i - \mathbf{J}^{-1}\boldsymbol{f}(\boldsymbol{u}_i) \tag{1.49}$$

to determine the unknown vector $\boldsymbol{u}$**,** where the vectors $\boldsymbol{u}_i$ and $\boldsymbol{u}_{i+1}$ are respectively the solutions for $i$ and $i+1$ iterations. To find the solutions of the nonlinear equations, we use the following initial guess $\boldsymbol{u}_0$

$$\boldsymbol{u}_0 = \{0 \quad 0 \quad \ldots \quad 0\}^{\mathrm{T}}_{1\times 12} \tag{1.50}$$

and the convergence criterion

$$|\boldsymbol{f}| = \sqrt{(f_1)^2 + (f_2)^2 + \cdots + (f_{12})^2} < \epsilon_{\mathrm{Tol}} \tag{1.51}$$

where $|\boldsymbol{f}|$ is the magnitude of the vector $\boldsymbol{f}$, and $\epsilon_{\mathrm{Tol}}$ is the convergence threshold. Using $\epsilon_{\mathrm{Tol}} = 10^{-8}$ in our simulations, we stop the iterations in (1.49) when $|\boldsymbol{f}|$ is less than $\epsilon_{\mathrm{Tol}}$. With the strain tensors $\varepsilon_{ij}^{(\mathrm{X})}$ and $\varepsilon_{ij}^{(\mathrm{Y})}$ determined from (1.49), we can calculate the stress tensor $\sigma_{ij}$ (from (1.16) or (1.24) and the stiffness tensor $\mathrm{C}_{ij}$ (from (1.39)).

#### 2 Three-dimensional focal adhesion model

In our finite element framework, the focal adhesion is treated as a thin, initially soft layer, which undergoes stiffening when tensile forces exceed a specific threshold. This is captured by decomposing the total Cauchy stress tensor $\hat{\boldsymbol{\sigma}} = \hat{\boldsymbol{\sigma}}^{(\mathrm{I})} + \hat{\boldsymbol{\sigma}}^{(\mathrm{F})}$ into the initial, $\hat{\boldsymbol{\sigma}}^{(\mathrm{I})}$, and stiffening, $\hat{\boldsymbol{\sigma}}^{(\mathrm{F})}$, parts

$$\hat{\sigma}_{ij} = \hat{\sigma}_{ij}^{(\mathrm{I})} + \hat{\sigma}_{ij}^{(\mathrm{F})} \tag{2.1}$$

$\hat{\boldsymbol{\sigma}}^{(\mathrm{I})}$ represents the initially pliable behavior of focal adhesions and is linearly related to the strain tensor $\hat{\boldsymbol{\varepsilon}}$ using the stiffness tensor $\hat{\mathbf{C}}^{(\mathrm{I})}$

$$\hat{\sigma}_{ij}^{(\mathrm{I})} = \hat{\mathrm{C}}_{ijkl}^{(\mathrm{I})}\, \hat{\varepsilon}_{kl} \tag{2.2}$$

The initial stiffness tensor of the focal adhesion layer $\hat{\mathrm{C}}_{ijkl}^{(\mathrm{I})}$ is determined as follows

$$\hat{\mathrm{C}}_{ijkl}^{(\mathrm{I})} = \hat{\kappa}\delta_{ij}\delta_{kl} + \hat{\mu}\left(\delta_{ik}\delta_{jk} + \delta_{il}\delta_{jk} - \frac{2}{3}\delta_{ij}\delta_{kl}\right) \tag{2.3}$$

where $\delta_{ij}$ is Kronecker's delta, $\hat{\kappa}$ is the bulk modulus

$$\hat{\kappa} = \frac{\hat{E}}{3(1-2\hat{v})} \tag{2.4}$$

and $\hat{\mu}$ is the shear modulus of the focal adhesion layer

$$\hat{\mu} = \frac{\hat{E}}{2(1+\hat{v})} \tag{2.5}$$

The two parameters $\hat{E}$ and $\hat{v}$, required to determine the stiffness tensor $\hat{\mathbf{C}}^{(\mathrm{I})}$ in Eq, (2.3), are the initial elastic modulus and Poison's ratio of the focal adhesion layer, respectively. Note that a low value for $\hat{E}$ is chosen in our simulations to represent the initially soft behavior of focal adhesions.

The second part of the stress tensor, $\hat{\boldsymbol{\sigma}}^{(\mathrm{F})}$, represents the stiffening of focal adhesions in tension and is defined as a nonlinear function of the strain tensor $\hat{\boldsymbol{\varepsilon}}$

$$\hat{\boldsymbol{\sigma}}^{(\mathrm{F})} = \sum_{i=1}^{3} \frac{\partial \hat{f}(\hat{\varepsilon}_i)}{\partial \hat{\varepsilon}_i} \hat{\boldsymbol{m}}_i \otimes \hat{\boldsymbol{m}}_i = \sum_{i=1}^{3} \hat{\sigma}^{(\mathrm{F})}(\hat{\varepsilon}_i)\, \hat{\mathbf{E}}_i = \sum_{i=1}^{3} \hat{\sigma}_i^{(\mathrm{F})}\, \hat{\mathbf{E}}_i \tag{2.6}$$

where the orthogonal eigenvectors of stress $\hat{\boldsymbol{m}}_1$, $\hat{\boldsymbol{m}}_2$, and $\hat{\boldsymbol{m}}_3$ are respectively the unit vectors in the directions of the principal strains $\hat{\varepsilon}_1$, $\hat{\varepsilon}_2$, and $\hat{\varepsilon}_3$ and the symmetric tensors $\hat{\mathbf{E}}_1 = \hat{\boldsymbol{m}}_1 \otimes \hat{\boldsymbol{m}}_1$, $\hat{\mathbf{E}}_2 = \hat{\boldsymbol{m}}_2 \otimes \hat{\boldsymbol{m}}_2$, and $\hat{\mathbf{E}}_3 = \hat{\boldsymbol{m}}_3 \otimes \hat{\boldsymbol{m}}_3$ are the eigenprojections of $\hat{\boldsymbol{\varepsilon}}$. Note that $\otimes$ denotes the dyadic product of two arbitrary vectors $\boldsymbol{u}$ and $\boldsymbol{v}$ as $(\boldsymbol{u} \otimes \boldsymbol{v})_{ij} = u_i v_j$.

Since $\hat{\boldsymbol{\sigma}}^{(\mathrm{F})}$ in Eq. (2.6) is defined as a function of its principal stresses ($\hat{\sigma}_1^{(\mathrm{F})}$, $\hat{\sigma}_2^{(\mathrm{F})}$, $\hat{\sigma}_3^{(\mathrm{F})}$) and eigenprojections ($\hat{\mathbf{E}}_1$, $\hat{\mathbf{E}}_2$, $\hat{\mathbf{E}}_3$), our next step involves the determination of these principal stresses and eigenprojections. The eigenprojections can be determined from the following spectral decomposition of the strain tensor $\hat{\boldsymbol{\varepsilon}}$

$$\hat{\boldsymbol{\varepsilon}} = \sum_{i=1}^{3} \hat{\varepsilon}_i \, \hat{\boldsymbol{m}}_i \otimes \hat{\boldsymbol{m}}_i = \sum_{i=1}^{3} \hat{\varepsilon}_i \, \hat{\mathbf{E}}_i \tag{2.7}$$

The principal stresses ($\hat{\sigma}_1^{(\mathrm{F})}$, $\hat{\sigma}_2^{(\mathrm{F})}$, $\hat{\sigma}_3^{(\mathrm{F})}$) are defined as functions of the principal strains ($\hat{\varepsilon}_1$, $\hat{\varepsilon}_2$, $\hat{\varepsilon}_3$) using the following constitutive equations

$$\hat{\sigma}_i^{(\mathrm{F})} = \frac{\partial \hat{f}(\hat{\varepsilon}_i)}{\partial \hat{\varepsilon}_i} = \tag{2.8}$$

$$\begin{cases} 0 & \hat{\varepsilon}_i < \hat{\epsilon}_1 \\ \hat{\ell}\left[\dfrac{\left(\dfrac{\hat{\varepsilon}_i - \hat{\epsilon}_1}{\hat{\epsilon}_2 - \hat{\epsilon}_1}\right)^{\hat{t}} (\hat{\varepsilon}_i - \hat{\epsilon}_1)^2}{(\hat{t}+1)(\hat{t}+2)}\right] & \hat{\epsilon}_1 \leq \hat{\varepsilon}_i < \hat{\epsilon}_2 \\ \hat{\ell}\left[\dfrac{(1+\hat{\varepsilon}_i - \hat{\epsilon}_2)^{\hat{s}+2} - 1}{(\hat{s}+1)(\hat{s}+2)} + \dfrac{\hat{\epsilon}_2 - \hat{\varepsilon}_i}{\hat{s}+1} + \dfrac{(\hat{\varepsilon}_i - \hat{\epsilon}_2)(\hat{\epsilon}_2 - \hat{\epsilon}_1)}{\hat{t}+1} + \dfrac{(\hat{\epsilon}_2 - \hat{\epsilon}_1)^2}{(\hat{t}+1)(\hat{t}+2)}\right] & \hat{\varepsilon}_i \geq \hat{\epsilon}_2 \end{cases}$$

where the principal strains ($\hat{\varepsilon}_1$, $\hat{\varepsilon}_2$, $\hat{\varepsilon}_3$) are obtained from (2.7). In Eq. (2.8), $\hat{\epsilon}_1 = \hat{\epsilon}_\mathrm{c} - 0.5\hat{\epsilon}_\mathrm{t}$ and $\hat{\epsilon}_2 = \hat{\epsilon}_\mathrm{c} + 0.5\hat{\epsilon}_\mathrm{t}$ are the transition points where focal adhesions transition into a stiffer state. $\hat{\epsilon}_\mathrm{t} = 0.25\hat{\epsilon}_\mathrm{c}$ is the transition width, $\hat{\epsilon}_\mathrm{c}$ is the critical (tensile) principal strain, and $\hat{t}$ is the transition constant. $\hat{\ell}$ and $\hat{s}$ are also the stiffening parameters and regulate the strain-stiffening behavior of focal adhesions.

Similar to Eq. (2.1), the total stiffness tensor $\hat{\mathbf{C}}$ is decomposed into the initial, $\hat{\mathbf{C}}^{(\mathrm{I})}$, and stiffening, $\hat{\mathbf{C}}^{(\mathrm{F})}$, parts

$$\hat{\mathrm{C}}_{ijkl} = \hat{\mathrm{C}}_{ijkl}^{(\mathrm{I})} + \hat{\mathrm{C}}_{ijkl}^{(\mathrm{F})} \tag{2.9}$$

where $\hat{\mathbf{C}}^{(\mathrm{I})}$ is determined from Eq. (2.3) and the strain-stiffening term $\hat{\mathbf{C}}^{(\mathrm{F})}$ is determined using the following piecewise linear approximation

$$\hat{\mathrm{C}}_{ijkl}^{(\mathrm{F})} = \frac{\mathrm{d}\hat{\sigma}_{ij}^{(\mathrm{F})}}{\mathrm{d}\hat{\varepsilon}_{kl}} \quad \text{or} \quad \hat{\mathbf{C}}^{(\mathrm{F})} = \frac{\mathrm{d}\hat{\boldsymbol{\sigma}}^{(\mathrm{F})}}{\mathrm{d}\hat{\boldsymbol{\varepsilon}}} \tag{2.10}$$

Note that Eq. (2.10) gives the principal stress $\hat{\sigma}_i^{(\mathrm{F})}$ as a function of the principal strain $\hat{\varepsilon}_i$ and not $\hat{\sigma}_{ij}^{(\mathrm{F})}$ as a function of $\hat{\varepsilon}_{ij}$. Thus, using the spectral decomposition of $\hat{\boldsymbol{\sigma}}^{(\mathrm{F})}$ in (2.6), $\hat{\mathbf{C}}^{(\mathrm{F})}$ is written in the following form:

$$\hat{\mathbf{C}}^{(\mathrm{F})} = \sum_{i=1}^{3} \left\{ \hat{\mathbf{E}}_i \otimes \frac{\mathrm{d}\hat{\sigma}_i^{(\mathrm{F})}}{\mathrm{d}\hat{\boldsymbol{\varepsilon}}} + \hat{\sigma}_i^{(\mathrm{F})} \frac{\mathrm{d}\hat{\mathbf{E}}_i}{\mathrm{d}\hat{\boldsymbol{\varepsilon}}} \right\} \tag{2.11}$$

which can be further expanded by applying the chain rule to the first term of the above equation

$$\hat{\mathbf{C}}^{(\mathrm{F})} = \sum_{i=1}^{3}\left\{\sum_{j=1}^{3}\frac{\partial\hat{\sigma}_i^{(\mathrm{F})}}{\partial\hat{\varepsilon}_j}\hat{\mathbf{E}}_i\otimes\frac{\mathrm{d}\hat{\varepsilon}_j}{\mathrm{d}\hat{\boldsymbol{\varepsilon}}}+\hat{\sigma}_i^{(\mathrm{F})}\frac{\mathrm{d}\hat{\mathbf{E}}_i}{\mathrm{d}\hat{\boldsymbol{\varepsilon}}}\right\} \tag{2.12}$$

If $\hat{\varepsilon}_{ij}$ has three distinct eigenvalues ($\hat{\varepsilon}_1 \neq \hat{\varepsilon}_2 \neq \hat{\varepsilon}_3$), by taking the derivatives of $\hat{\varepsilon}_j$ and $\hat{\mathbf{E}}_i$ with respect to $\hat{\boldsymbol{\varepsilon}}$ in Eq. (2.12), we can derive $\hat{\mathbf{C}}^{(\mathrm{F})}$ as follows

$$\begin{aligned}\hat{\mathbf{C}}^{(\mathrm{F})} &= \sum_{a=1}^{3}\frac{\hat{\sigma}_a^{(\mathrm{F})}}{(\hat{\varepsilon}_a-\hat{\varepsilon}_b)(\hat{\varepsilon}_a-\hat{\varepsilon}_c)}\left\{\frac{\mathrm{d}(\hat{\boldsymbol{\varepsilon}})^2}{\mathrm{d}\hat{\boldsymbol{\varepsilon}}}-(\hat{\varepsilon}_b+\hat{\varepsilon}_c)\mathbf{I}_{\mathrm{S}}-[(\hat{\varepsilon}_a-\hat{\varepsilon}_b)+(\hat{\varepsilon}_a-\hat{\varepsilon}_c)]\hat{\mathbf{E}}_a\otimes\hat{\mathbf{E}}_a\right.\\ &\qquad\left.-(\hat{\varepsilon}_b-\hat{\varepsilon}_c)\left(\hat{\mathbf{E}}_b\otimes\hat{\mathbf{E}}_b-\hat{\mathbf{E}}_c\otimes\hat{\mathbf{E}}_c\right)\right\}+\sum_{i=1}^{3}\sum_{j=1}^{3}\frac{\partial\hat{\sigma}_i^{(\mathrm{F})}}{\partial\hat{\varepsilon}_j}\hat{\mathbf{E}}_i\otimes\hat{\mathbf{E}}_j\end{aligned} \tag{2.13}$$

where the fourth-order tensor $\frac{\mathrm{d}(\hat{\boldsymbol{\varepsilon}})^2}{\mathrm{d}\hat{\boldsymbol{\varepsilon}}}$ is the derivative of the square of the second-order tensor $\hat{\boldsymbol{\varepsilon}}$

$$\left(\frac{\mathrm{d}(\hat{\boldsymbol{\varepsilon}})^2}{\mathrm{d}\hat{\boldsymbol{\varepsilon}}}\right)_{ijkl} = \frac{1}{2}\left(\delta_{ik}\hat{\varepsilon}_{lj}+\delta_{il}\hat{\varepsilon}_{kj}+\delta_{jl}\hat{\varepsilon}_{ik}+\delta_{kj}\hat{\varepsilon}_{il}\right) \tag{2.14}$$

and $\mathbf{I}_{\mathrm{S}}$ is the following fourth-order symmetric identity tensor

$$(\mathbf{I}_{\mathrm{S}})_{ijkl} = \frac{1}{2}\left(\delta_{ik}\delta_{jl}+\delta_{il}\delta_{jk}\right) \tag{2.15}$$

If there are two identical eigenvalues ($\hat{\varepsilon}_1 \neq \hat{\varepsilon}_2 = \hat{\varepsilon}_3$), $\hat{\mathbf{C}}^{(\mathrm{F})}$ is determined as follows

$$\hat{\mathbf{C}}^{(\mathrm{F})} = s_1\frac{\mathrm{d}(\hat{\boldsymbol{\varepsilon}})^2}{\mathrm{d}\hat{\boldsymbol{\varepsilon}}}-s_2\mathbf{I}_{\mathrm{S}}-s_3\hat{\boldsymbol{\varepsilon}}\otimes\hat{\boldsymbol{\varepsilon}}+s_4\hat{\boldsymbol{\varepsilon}}\otimes\mathbf{I}+s_5\mathbf{I}\otimes\hat{\boldsymbol{\varepsilon}}-s_6\mathbf{I}\otimes\mathbf{I} \tag{2.16}$$

where $\mathbf{I}$ is the second-order identity tensor

$$\mathrm{I}_{ij} = \delta_{ij} \tag{2.17}$$

and the constants $s_1$, $s_2$, $s_3$, $s_4$, $s_5$, and $s_6$ are given as follows

$$s_1 = \frac{\hat{\sigma}_a^{(\mathrm{F})}-\hat{\sigma}_c^{(\mathrm{F})}}{(\hat{\varepsilon}_a-\hat{\varepsilon}_c)^2}+\frac{1}{\hat{\varepsilon}_a-\hat{\varepsilon}_c}\left(\frac{\partial\hat{\sigma}_c^{(\mathrm{F})}}{\partial\hat{\varepsilon}_b}-\frac{\partial\hat{\sigma}_c^{(\mathrm{F})}}{\partial\hat{\varepsilon}_c}\right) \tag{2.18a}$$

$$s_2 = 2\hat{\varepsilon}_c\frac{\hat{\sigma}_a^{(\mathrm{F})}-\hat{\sigma}_c^{(\mathrm{F})}}{(\hat{\varepsilon}_a-\hat{\varepsilon}_c)^2}+\frac{\hat{\varepsilon}_a+\hat{\varepsilon}_c}{\hat{\varepsilon}_a-\hat{\varepsilon}_c}\left(\frac{\partial\hat{\sigma}_c^{(\mathrm{F})}}{\partial\hat{\varepsilon}_b}-\frac{\partial\hat{\sigma}_c^{(\mathrm{F})}}{\partial\hat{\varepsilon}_c}\right) \tag{2.18b}$$

$$s_3 = 2\frac{\hat{\sigma}_a^{(\mathrm{F})}-\hat{\sigma}_c^{(\mathrm{F})}}{(\hat{\varepsilon}_a-\hat{\varepsilon}_c)^3}+\frac{1}{(\hat{\varepsilon}_a-\hat{\varepsilon}_c)^2}\left(\frac{\partial\hat{\sigma}_a^{(\mathrm{F})}}{\partial\hat{\varepsilon}_c}+\frac{\partial\hat{\sigma}_c^{(\mathrm{F})}}{\partial\hat{\varepsilon}_a}-\frac{\partial\hat{\sigma}_a^{(\mathrm{F})}}{\partial\hat{\varepsilon}_a}-\frac{\partial\hat{\sigma}_c^{(\mathrm{F})}}{\partial\hat{\varepsilon}_c}\right) \tag{2.18c}$$

$$s_4 = 2\hat{\varepsilon}_c \frac{\hat{\sigma}_a^{(\mathrm{F})} - \hat{\sigma}_c^{(\mathrm{F})}}{(\hat{\varepsilon}_a - \hat{\varepsilon}_c)^3} + \frac{1}{\hat{\varepsilon}_a - \hat{\varepsilon}_c}\left(\frac{\partial \hat{\sigma}_a^{(\mathrm{F})}}{\partial \hat{\varepsilon}_c} - \frac{\partial \hat{\sigma}_c^{(\mathrm{F})}}{\partial \hat{\varepsilon}_b}\right) + \frac{\hat{\varepsilon}_c}{(\hat{\varepsilon}_a - \hat{\varepsilon}_c)^2}\left(\frac{\partial \hat{\sigma}_a^{(\mathrm{F})}}{\partial \hat{\varepsilon}_c} + \frac{\partial \hat{\sigma}_c^{(\mathrm{F})}}{\partial \hat{\varepsilon}_a} - \frac{\partial \hat{\sigma}_a^{(\mathrm{F})}}{\partial \hat{\varepsilon}_a} - \frac{\partial \hat{\sigma}_c^{(\mathrm{F})}}{\partial \hat{\varepsilon}_c}\right) \tag{2.18d}$$

$$s_5 = 2\hat{\varepsilon}_c \frac{\hat{\sigma}_a^{(\mathrm{F})} - \hat{\sigma}_c^{(\mathrm{F})}}{(\hat{\varepsilon}_a - \hat{\varepsilon}_c)^3} + \frac{1}{\hat{\varepsilon}_a - \hat{\varepsilon}_c}\left(\frac{\partial \hat{\sigma}_c^{(\mathrm{F})}}{\partial \hat{\varepsilon}_a} - \frac{\partial \hat{\sigma}_c^{(\mathrm{F})}}{\partial \hat{\varepsilon}_b}\right) + \frac{\hat{\varepsilon}_c}{(\hat{\varepsilon}_a - \hat{\varepsilon}_c)^2}\left(\frac{\partial \hat{\sigma}_a^{(\mathrm{F})}}{\partial \hat{\varepsilon}_c} + \frac{\partial \hat{\sigma}_c^{(\mathrm{F})}}{\partial \hat{\varepsilon}_a} - \frac{\partial \hat{\sigma}_a^{(\mathrm{F})}}{\partial \hat{\varepsilon}_a} - \frac{\partial \hat{\sigma}_c^{(\mathrm{F})}}{\partial \hat{\varepsilon}_c}\right) \tag{2.18e}$$

$$s_6 = 2\hat{\varepsilon}_c \frac{\hat{\sigma}_a^{(\mathrm{F})} - \hat{\sigma}_c^{(\mathrm{F})}}{(\hat{\varepsilon}_a - \hat{\varepsilon}_c)^3} + \frac{\hat{\varepsilon}_a \hat{\varepsilon}_c}{(\hat{\varepsilon}_a - \hat{\varepsilon}_c)^2}\left(\frac{\partial \hat{\sigma}_a^{(\mathrm{F})}}{\partial \hat{\varepsilon}_c} + \frac{\partial \hat{\sigma}_c^{(\mathrm{F})}}{\partial \hat{\varepsilon}_a}\right) - \frac{(\hat{\varepsilon}_c)^2}{(\hat{\varepsilon}_a - \hat{\varepsilon}_c)^2}\left(\frac{\partial \hat{\sigma}_a^{(\mathrm{F})}}{\partial \hat{\varepsilon}_a} + \frac{\partial \hat{\sigma}_c^{(\mathrm{F})}}{\partial \hat{\varepsilon}_c}\right) - \frac{\hat{\varepsilon}_a + \hat{\varepsilon}_c}{\hat{\varepsilon}_a - \hat{\varepsilon}_c}\frac{\partial \hat{\sigma}_c^{(\mathrm{F})}}{\partial \hat{\varepsilon}_b} \tag{2.18f}$$

where $(a, b, c)$ are here cyclic permutations of (1,2,3).

Finally, if there are three identical eigenvalues ($\hat{\varepsilon}_1 = \hat{\varepsilon}_2 = \hat{\varepsilon}_3$), $\hat{\mathbf{C}}^{(\mathrm{F})}$ is obtained as follows

$$\hat{\mathbf{C}}^{(\mathrm{F})} = \left(\frac{\partial \hat{\sigma}_1^{(\mathrm{F})}}{\partial \hat{\varepsilon}_1} - \frac{\partial \hat{\sigma}_1^{(\mathrm{F})}}{\partial \hat{\varepsilon}_2}\right)\mathbf{I}_{\mathrm{S}} + \frac{\partial \hat{\sigma}_1^{(\mathrm{F})}}{\partial \hat{\varepsilon}_2}\mathbf{I} \otimes \mathbf{I} \tag{2.19}$$

Note that $\frac{\partial \hat{\sigma}_i^{(\mathrm{F})}}{\partial \hat{\varepsilon}_j}$ in (2.13), (2.18), and (2.19) can be obtained by taking the derivative of $\hat{\sigma}_i^{(\mathrm{F})}$ in Eq. (2.8) where $\hat{\sigma}_i^{(\mathrm{F})}$ is a function of $\hat{\varepsilon}_i$ only and $\hat{\sigma}_i^{(\mathrm{F})}$ does not depend on $\hat{\varepsilon}_j$ for $i \neq j$.

$$\frac{\partial \hat{\sigma}_i^{(\mathrm{F})}}{\partial \hat{\varepsilon}_i} = \frac{\partial}{\partial \hat{\varepsilon}_i}\left(\frac{\partial \hat{f}}{\partial \hat{\varepsilon}_i}\right) = \begin{cases} 0 & \hat{\varepsilon}_i < \hat{\epsilon}_1 \\ \hat{\ell}\left[\dfrac{\left(\dfrac{\hat{\varepsilon}_i - \hat{\epsilon}_1}{\hat{\epsilon}_2 - \hat{\epsilon}_1}\right)^{\hat{t}} (\hat{\varepsilon}_i - \hat{\epsilon}_1)}{\hat{t} + 1}\right] & \hat{\epsilon}_1 \le \hat{\varepsilon}_i < \hat{\epsilon}_2 \\ \hat{\ell}\left[\dfrac{(1 + \hat{\varepsilon}_i - \hat{\epsilon}_2)^{\hat{s}+1} - 1}{\hat{s} + 1} + \dfrac{\hat{\epsilon}_2 - \hat{\epsilon}_1}{\hat{t} + 1}\right] & \hat{\varepsilon}_i \ge \hat{\epsilon}_2 \end{cases} \tag{2.20}$$

### 3 Additional Details of Deep Learning Components

#### 3.1 Phase to Fluorescent Image Translation using pix2pix

##### 3.1.1 Model Summary: pix2pix

The pix2pix model belongs to the class of deep learning models referred to as conditional Generative Adversarial Networks (cGANs)[10], which comprises of two neural network models, a generator $G$ and a discriminator $D$. The generator learns a mapping from the observed input image $x$ (from phase microscopy) and a random noise vector $z$ (simulated through dropout[11] as originally proposed in the pix2pix paper) to the output image $y$ (fluorescent), $G: \{x, z\} \rightarrow y$. The generator $G$ is trained to produce synthetic (or "fake") fluorescent images that cannot be distinguished from the "real" fluorescent images on the training set. On the other hand, the discriminator $D$ is adversarially trained to distinguish between the generated "fake" images and the "real" ones on the training set. Both the generator and discriminator neural networks are simultaneously trained by optimizing the following objective function

$$L_{cGAN}(G, D) = \mathbb{E}_{(x,y)}\,[\log D(x, y)] + \mathbb{E}_{(x,z)}\,\left[\log\left(1 - D\big(x, G(x, z)\big)\right)\right] + \lambda\, L_{L1}(G) \tag{3.1}$$

$$L_{L1}(G) = \mathbb{E}_{(x,y,z)}\,[\|y - G(x, z)\|_1], \tag{3.2}$$

where $\mathbb{E}[\cdot]$ denotes the expectation over the training set, $D(x, y)$ represents the real-valued score between 0 to 1 produced by the discriminator (ideally 0 for fake outputs and 1 for real outputs), $G(x, z)$ represents the fake fluorescent outputs of the generator, $L_{L1}(G)$ is the L1-norm between the generated "fake" images and the "real" images over the training set, and $\lambda$ is a trade-off hyper-parameter that weights the importance of the L1-norm in the optimization objective. The goal of the generator $G$ is to minimize this objective function (thus "fooling" the discriminator to produce similar scores for the fake outputs as the real ones), while the discriminator $D$ tries to maximize the objective function (so as to challenge the generator to generate better synthetic outputs). The L1-norm term additionally encourages the generator to produce outputs to be close to the "real" outputs ($\lambda = 100$ was used for training).

##### 3.1.2 Training and Testing Setups for pix2pix

To overcome the challenges of blurring, grainy, and photo-bleaching associated with the fluorescent images, we used Sharpness Index[12] (SI) as a quantitative metric describing the quality of an image $I$. Since these artifacts may sometimes appear in small local regions (or patches) of the image, we divided every image $I$ into $4 \times 4$ patches, and computed the SI of the entire image as the minimum SI of all patches as follows:

$$\mathrm{SI}(I) = \min_{1 \leq i,j \leq 4} \mathbf{var}\left(\mathrm{Laplacian}\big(I_{i,j}\big)\right) \tag{3.3}$$

where $I_{i,j}$ is the image patch extracted from the $i$-th row and $j$-th column, the Laplacian operator generates the 2-nd derivative of an image (quantifying steep intensity changes within an image), and the $\mathbf{var}(\cdot)$ operator computes the variance of a matrix.

SI analysis allowed us to create a "pristine" set of phase-fluorescent image pairs (example shown in Fig. 3Aii) for training our model by excluding images with SI < 400 (blurry) or those with artifacts (see Supplementary Figure 5 for some examples).

We divide the entire set of images from 273 time-lapse videos into a training set consisting of 208 videos (which roughly translates to a training fraction of 0.75) and a test set consisting of the remaining 65 videos. Both the training and test sets were partitioned into two subsets: $\mathrm{SI} \geq 400$ and $\mathrm{SI} < 400$. The set of images with $\mathrm{SI} \geq 400$ were further subdivided through manual supervision into 4 categories: pristine set, partially blurry set of images, set of images with hazy fiber networks, and set of images with bright spot/background noise, both over training and test partitions. Also, an additional subset of images was selected from the set of images with $\mathrm{SI} < 400$ and tagged as the completely blurry set. More specifically, the 208 videos that were selected for training comprised of 10500 images, out of which only 3119 images had $\mathrm{SI} \geq 400$. We subdivided these images into 4 categories as follows: (a) Pristine set: 850 images, (b) Partially-blurry set: 911 images, (c) Set of images with bright spot/background noise: 952 images, and (d) Set of images with hazy fiber networks: 406 images. From the 7381 images which had $\mathrm{SI} < 400$, a completely blurry set of 600 images were manually selected. A similar scheme was used to partition the 65 videos in the test set.

The $1388 \times 1040$ resolution images were obtained experimentally where 1 pixel corresponds to 0.32 micrometer. As a pre-processing step, we re-scaled all images to $1024 \times 1024$ before feeding them into the pix2pix model, which generated fluorescent images of the same resolution. The generated fluorescent images were re-scaled back to the original dimension before tracking the nodal intersections. Our pix2pix architecture was same as the one proposed in the original paper.[13]

##### 3.1.3 Ablation Study of Proposed pix2pix Model

We also performed an in-depth study to evaluate the performance of pix2pix on various training and testing setups (See Supplementary Table 3-1). It is observed that the pix2pix performs best when it is trained on the pristine set, with 2-d cross-correlation (2-d cross-correlation equal to 1 represents perfect reconstruction of the fluorescent images by the pix2pix) of 0.887 even evaluated on the Pristine Set. Although it might seem that training on pix2pix and partially blurry set achieved similar performance (with correlation values of 0.88), it should be noted that images tagged as partially blurry were done under very stringent conditions determined manually. Also, the size of the training set was significantly larger (almost twice as compared to the pristine set), which also affects the training time by a large margin. It can be seen that the addition of other sets such as hazy and bright spot slightly degraded the performance of the pix2pix on the pristine set, although there were significant increases in correlation values on the hazy and bright sets. This is due to the fact that when the pix2pix is trained using the hazy/bright spot set, it is able to learn these imperfections and replicate them on the training data. Ideally, it is not desirable to have these artefacts in our predictions. Also, it is not surprising that when trained on the completely blurry set, the pix2pix performance significantly dropped for all the testing setups (See last row of Supplementary Table 3-1). Thus, it can be concluded that training the pix2pix on the pristine set is ideal and addition of other imperfections in the training data only hampers the model performance on unseen test instances.

| | | **Testing Setup** | | | | |
|---|---|---|---|---|---|---|
| | | Pristine | Partially Blurry | Hazy | Bright | Completely Blurry |
| **Training Setup** | Pristine Set | 0.887 | 0.877 | 0.702 | 0.667 | 0.335 |
| | Pristine + Blurry (SI > 400) | 0.88 | 0.886 | 0.697 | 0.676 | 0.418 |
| | Pristine + Hazy | 0.859 | 0.889 | 0.762 | 0.671 | 0.371 |
| | Pristine + Bright | 0.867 | 0.871 | 0.745 | 0.78 | 0.413 |

| | | | | | | |
|---|---|---|---|---|---|---|
| | Completely Blurry (SI < 400) | 0.604 | 0.621 | 0.373 | 0.482 | 0.344 |

Supplementary Table 3-1: Ablation study of proposed image-translation model (pix2pix) where the 2-d cross-correlation values for each Training/Testing combination is used to compare the different setups.

#### 3.2 Node Detection and Tracking

##### 3.2.1 Model Summary: Object Detection using RetinaNet

Object Detection[14] is the task of identifying or detecting objects in an image. Objects in an image are usually depicted by drawing a box around its extent, referred to as a bounding box. The task of detecting objects is quite natural for humans as the human visual system can effortlessly detect objects at a glance. However, to replicate the effectiveness of human visual systems for object detection in machines, we commonly rely on computer vision algorithms based on deep learning that are capable of learning image features relating to specific classes of objects in an image. In the following, we describe our approach for detecting intersection points (nodes) of fiber networks in synthetic fluorescent images using deep learning-based object detection methods.

We used the RetinaNet[15] model for node detection, which is one of the commonly used object detection models in the field of computer vision. The architecture of the RetinaNet model comprises of a neural network that extracts image-based features from a pretrained residual network (ResNet)[16] model, a feature pyramid network (FPN)[17] model that converts the ResNet features into a hierarchy of down-scaled features at multiple spatial scales, and two sub-networks for performing classification and regression on bounding boxes, respectively (see a schematic representation of RetinaNet architecture in Figure 3Bi). The feature maps from the pretrained ResNet model and the FPN model form the backbone of the RetinaNet architecture. The bottom-up path (from the image to extracted features) uses a convolutional neural network[18] architecture with weights that are pretrained on the ImageNet dataset[19], which is a publicly available dataset containing more than 1 million images of real-world objects. The top-down network of the FPN downscales these feature maps and combines information at different resolutions. This backbone helps the RetinaNet in being scale- and resolution-independent. RetinaNet uses anchors at various spatial locations in all the feature map levels from the FPN to generate potential regions of different sizes and aspect ratios that are likely to contain the objects of interest. The classification sub-network, which is a convolutional neural network, predicts the probability of an object being present and its class, based on the input anchors. A second sub-network regresses the anchor boxes to predict accurate and well-fitting bounding boxes around each detected object. Together, the output of the two subnetworks gives the predicted bounding boxes and the class probabilities of objects present in each bounding box of the input image. RetinaNet utilizes a variation of the Cross Entropy Loss known as the Focal Loss as its learning objective. The goal of focal loss is to increase the contribution of hard (or often misclassified) samples over the easy samples in the loss computation. This is done to address the challenge of class imbalance between objects of the foreground and background classes. Specifically, the focal loss for an object detection model's predicted class probabilities is defined using the following equation:

$$FL(p_t) = -\alpha_t (1 - p_t)^{\gamma} \log(p_t) \tag{3.4}$$

$$p_t = \begin{cases} p & \text{if } y = 1 \\ 1 - p & \text{otherwise} \end{cases} \tag{3.5}$$

where, $y$ is the ground truth class of an object and $p$ is the model's predicted probability for the ground class of an object. The hyper-parameter $\alpha$ modulates the effect of class imbalance already presents in the dataset while $\gamma$ regulates the effect of hard samples on the loss. The higher the value of $\gamma$, the more is the importance of hard samples, and this penalizes the model more for hard samples than easy samples.

Most object detection datasets have few instances of the same object class in an image. On the contrary, we have a large number of nodes in every frame (typically of the order $100 \sim 500$), which requires appropriate adaptation of the RetinaNet hyper-parameters. In particular, we increase the number of test detections per image and the proposals per feature map as compared to the default values of the RetinaNet. Additionally, we perform post-processing of the predicted nodes by thresholding based on their confidence scores from the object detection module, i.e., bounding boxes with confidence scores less than 0.7 are eliminated. The remaining bounding boxes are then filtered by applying Non-Maximal Suppression[20] to remove duplicate or overlapping boxes based on their Intersection over Union (IoU) score[20] such that only the bounding box with the highest confidence score (termed the maximal box) is retained and overlapping boxes whose IoU score with the maximal box is greater than 0.95 are eliminated.

##### 3.2.2 Tracking Node Intersections Across Frames

Object detection is only able to detect the nodes present in one particular frame using the spatial features of the nodes at that frame and does not utilize the information present in the previous and future frames. To overcome this limitation, we developed a bi-directional node tracking algorithm that first propagates information about the locations of detected nodes from the first frame to the last frame of the video using a forward pass. The backward pass then applies the same algorithm in the opposite direction, propagating information about the locations of tracked nodes from the last frame to the first frame. In this way, temporal information is leveraged across all past and future frames to track nodes in a current frame. The node tracking algorithm serves two purposes. First, it generates a mapping between the detected nodes at consecutive frames, such that every node is tracked from the first frame to the last frame. Second, it also ensures that if a node is detected even in a single frame of a video, then it can be propagated to all other frames before and after that frame. This increases the total number of correctly detected nodes in every frame and even if the object detection module has missed a node in a frame, we can recover it by jointly using spatial and temporal information.

Our proposed bi-directional node tracking algorithm comprises of two phases—the forward pass and the backward pass. Both these passes utilize a 2D-cross correlation module that projects the location of every node in a source image (e.g., frame at timestamp $t-1$) to a predicted location in a destination image (e.g., frame at timestamp $t$), referred to as a cross-correlated node. Specifically, to cross-correlate a particular node $i$ from a frame at time $t$ to the next frame, we crop a small source patch $I_t^i$ of $N \times N$ pixels centered around that node ($N = 35$, Supplementary Figure 7). We crop a larger destination patch $I_{t+1}^i$ of size $M \times M$ from the next frame ($M = 55$) with the same center coordinates. We then compute the 2D-cross-correlation matrix between the source patch and destination patch and then take its Hadamard product with a symmetric Gaussian prior centered at the node location with 15 standard deviation in the source patch. The $(x, y)$ coordinates of the cross-node point for the next frame are then computed using the following formula:

$$\left(x_{t+1}^{i}, y_{t+1}^{i}\right) = \text{argmax}_{(x,y)}\mathbf{corr2D}\left(\mathrm{I}_{\mathrm{t}}^{\mathrm{i}}, \mathrm{I}_{\{\mathrm{t+1}\}}^{\mathrm{i}}\right) \tag{3.6}$$

where $i \in \{1,2,\dots,N\}$, $t \in \{1,2,\dots,T-1\}$, and $\mathbf{corr2D}$ is the result of the Hadamard product between the 2-D cross correlation operator and the Gaussian prior. This ensures that points closer to the location of the nodal point in the source frame are given a higher weight than those that are farther away, thus reducing the sensitivity to distant bright spots in the large $M \times M$ window around the source location. In our implementation, the values of $M$ and $N$ were chosen in such a way that the patches generally contain only one node intersection.

In the forward pass, after performing cross-correlation, we have two different sets of nodes for every frame starting from the second frame of a video: the cross-correlated nodes obtained by propagating information from the previous frame, and the detected nodes. We compute the Intersection over Union (IoU)[20] between the bounding boxes of these two sets of nodes at every frame. For the cross-correlated nodes that match with the detected nodes (i.e., show an IoU score greater than 0.75), the bounding boxes of the detected nodes are used to create the set of tracked nodes at every frame. This is done because the deep learning-based object detection module is expected to be more accurate at drawing bounding boxes around nodes than the cross-correlation module, as it can extract sophisticated spatial features corresponding to intersection points from the images. However, the detection module can still miss nodes that are difficult to detect solely based on spatial information. Hence, we add the bounding boxes of cross-correlated nodes that have not been matched with any of the detected nodes to the set of tracked nodes at every frame. In this way, we combine the results of the object detection module and the cross-correlation module for all frames in a video from beginning to end in the forward pass. In the backward pass, we reverse the direction of information propagation from the last frame to the first frame of the video, now matching the tracked nodes from the forward pass at every frame with the cross-correlated nodes coming from the next frame in the future. Specifically, the cross-correlated nodes in the backward pass are computed by projecting the tracked nodes at timestamp t (source image) to timestamp $t-1$ (destination image). This is applied to every frame starting from the second last frame to the first frame. Once we generate the final tracked nodes across all frames in the backward pass, changes in the locations of tracked nodes across consecutive frames are utilized to calculate the nodal displacement fields for force extraction.

##### 3.2.3 Training and Testing Setups for Node Detection and Tracking

The dataset used for the object detection network consists of 165 synthetic fluorescent images generated using pix2pix, spread over 17 cell videos. The ground truth labels of intersection points (nodes) for these frames were generated semi-automatically using Harris Corner Detection[21] to detect nodes on the first frame, which were then cross correlated in the forward direction to all subsequent frames. This was followed by a manual cleaning step to remove false positives generated by the Harris Corner Detection algorithm due to the presence of image artifacts. This process created a set of synthetic fluorescent images with manually corrected list of ground-truth node locations, which was then used for training the RetinaNet model for node detection. We placed a bounding box of size $30 \times 30$ pixels around every ground-truth node to be used as labels in the training of RetinaNet. Additionally, we implemented 5-fold cross-validation[22] on the 17 videos to analyze the performance on different videos and grids. In particular, the 17 videos were randomly split into 5 folds, $F_1$ to $F_5$. At the $i$-th run of cross-validation (where $i$ varies from 1 to 5), videos from fold $F_i$ were used for testing the node detection and tracking modules, while videos from all remaining folds were used for training. This way, every video was used for testing in exactly one of the cross-validation runs. To calculate the accuracy of node detection and tracking during testing, an IoU matching scheme is applied between the predicted boxes and the ground-truth boxes as follows. A predicted box that matches with any of the ground-truth boxes with an IoU score greater than 0.75 was labeled as a *true positive* (TP), indicating correct prediction. A predicted box that could not be matched with any ground-truth box was labeled a *false positive* (FP), indicating incorrect prediction. A ground-truth box that could not be matched with any of the predicted boxes was labeled a *false negative* (FN), indicating missed detection. By counting the total number of TPs, FPs, and FNs, we computed the precision and recall values as:

$$\text{Precision} = \frac{\text{TP}}{\text{TP} + \text{FP}} \tag{3.7}$$

$$\text{Recall} = \frac{\text{TP}}{\text{TP} + \text{FN}} \tag{3.8}$$

We expect both precision and recall being equal to 1 for ideal predictions. We also computed the root mean square error (RMSE) between the centers of the predicted box and matched ground-truth box, as another metric to evaluate the nodal errors in our predictions. A lower value of RMSE signifies that the model was able to precisely detect the actual nodes. The final model trained on all 17 videos was used to predict nodes on unseen out-of-distribution videos during testing.

### 4 Processing steps in generating the finite-elemental model

An experimentally obtained phase image (Supplementary Figure 9a) is processed using the pix2pix algorithm to generate a fluorescent image of the nanonet. The grid intersection points are then identified on the deformed crosshatch network using the object detection and tracking algorithm (Supplementary Figure 9b). A cell mask is generated by manually tracing the boundaries of the cell (Supplementary Figure 9c). The cell mask provides the region where the unknown forces are to be resolved. A bounding box of a size sufficiently larger than the cell (*i.e.* box size ~100-200 µm larger than the cell diameter) is used for the inverse analysis, see a representative example in Supplementary Figure 9d.

The reference configuration of the nanonet (*i.e.* fiber positions before the cells attach on nanonets) is not readily available due to several technical challenges in the current experimental setup. Therefore, we resort to a simple but effective approach to generate the reference configuration by leveraging the grid structure of the nanonets. The reference nanonet grid is generated by connecting the nodes on the opposite edges of the bounding box with straight. The bounding box is selected to be at least 2-3 grid rows or columns away from the cell boundary. The intersection of vertical and horizontal lines provides an estimate of the reference position at every grid intersection node within the bounding box. Although this approach is observed to work well, it is limited to rectilinear grid structures and further considerations are necessary to deal with non-rectangular nanonets, hexagonal nanonets for instance. Our ongoing work is directed towards improvements in the experimental protocol and computational implementation to reliably obtain the reference configurations.

The reference positions of grid intersections are used to generate the mesh with corotational beam elements (Supplementary Figure 9f). The surrounding region is meshed using triangular elements representing a homogenized Cosserat continuum (Supplementary Figure 9e), the outer box-size $L_b$ = 2000 µm is chosen to be sufficiently larger than the cell size to capture the experimental setup and limit boundary effects. The force pattern obtained after solving the inverse problem is shown in Supplementary Figure 10.

### 5 Formulation and solution procedure of the inverse problem

In this section, we first provide the detailed description of the finite-elemental model to capture the mechanical behavior of nanonets. Building on this forward problem formulation, we then present the traction force microscopy inverse problem on nanonets.

We consider a crosshatched nanonet roughly of size 400 µm × 400 µm for our analysis which corresponds to the typical size of the experimental phase images of the crosshatch network. Crosshatched nanonets can have square or rectangular grid patterns with a typical spacing of 8-25 µm between fibers. The spacing between the fibers and fiber diameters can be precisely controlled to generate a wide range of crosshatch network patterns[23]. For cells suspended between aligned fibers[24,25], cell exerted forces have been estimated from the deflection profile of a single nanofiber using the nanonet force microscopy (NFM) method[26]. Here we develop and validate a force microscopy approach based on crosshatched nanonets.

#### 5.1 Forward problem formulation

The nanofiber diameter *d* = 0.25 µm is sufficiently smaller than the typical fiber length determined by the grid spacing of about 25 µm. Nanofibers also have a significant pretension which contributes to the overall mechanical stiffness of the network. For fiber deflections an order of magnitude smaller than the fiber length and in the absence of nonlinear buckling under compression, mechanics of the nanofiber network can be captured using pre-tensed Euler-Bernoulli beams.[27] However, forces exerted by cells were observed to induce significantly larger deformations on the nanonets, especially for nanofibers close to and underneath the cells. To account for the geometrical nonlinearities associated with such large deformations, we model the mechanical response of nanofibers using the corotational beam formulation that accounts for large rigid body motions while still working under small-strain assumption in a corotating frame.[28] The corotational formulation for pre-tensed beams is presented next to set up the forward problem. Note that the significant pretension in nanofibers provides a sufficient buffer that limits the deformations to be in the elastic regime. For exceptionally high contractile forces (larger than 1000 nN per adhesion patch) plastic deformations may occur where this model is not applicable, but such cases are rarely observed in our experiments.

##### 5.1.1 Euler-Bernoulli beam with pretension

The mechanical response of a pre-tensed Euler-Bernoulli beam obtained from classical structural analysis[27] as a linearized relation between the end moments and small nodal rotations and transverse displacements is given as (Supplementary Figure 11)

$$\begin{bmatrix} M_a \\ M_b \end{bmatrix} = \begin{bmatrix} ks & ksc \\ ksc & ks \end{bmatrix} \begin{bmatrix} \theta_a - \psi \\ \theta_b - \psi \end{bmatrix} \tag{5.1}$$

where

$$\psi = \frac{v_b - v_a}{L}$$

is the small rigid rotation of the beam element, $v_a$ and $v_b$ are the transverse displacements at the ends, $\theta_a$ and $\theta_b$ are the nodal rotation at the ends, $M_a$ and $M_b$ are the end moments, $L$ is the length of the beam, and $F_0$ is the pretension force. For a beam with an uniform cross-section, $s$ and $c$ are defined as[27]

$$s \quad = \frac{\alpha(\alpha\cosh\,\alpha - \sinh\,\alpha)}{2 - 2\cosh\,\alpha + \alpha\sinh\,\alpha} \tag{5.2}$$

$$c \quad = \frac{\sinh\,\alpha - \alpha}{\alpha\cosh\,\alpha - \sinh\,\alpha} \tag{5.3}$$

where

$$\alpha = \sqrt{\frac{F_0}{EI}}L,$$

$E$ is the Young's modulus of the fiber material, $I$ is the second moment of area of the beam cross-section, and $k \ = \frac{EI}{L}$. The stretching response is obtained simply as

$$F = \frac{EA}{L}(u_b - u_a) + F_0.$$

For typical pretension and material stiffness values of nanofibers in this study, the bending stiffness generated by the fiber pre-tension is significantly larger than bending stiffness of the fiber itself. The overall mechanical behavior is therefore expected to be dominated by fiber pretension. The mechanical properties of nanofibers used in this study are summarized in Supplementary Table 5-1.[26]

| Parameter | Description | Value |
|---|---|---|
| $d$ | Fiber diameter | 250 nm |
| $F_0$ | Fiber pretension | 200 nN |
| $E$ | Young's modulus | 1 GPa |
| $L$ | Fiber spacing | 8-25 µm |

Supplementary Table 5-1: Summary of nanofiber properties.

##### 5.1.2 Corotational beam formulation

The corotational formulation captures the geometrical nonlinearities arising from large displacements and rotations in structural elements, while small strains are assumed in a local corotating frame. The 2D corotational beam framework proposed by Crisfield[28] is used here with the necessary modifications to account for the pretension in nanofibers.

A corotating reference frame is attached to each beam element such that rigid body translation and rotations can be separated from strain causing deformations in the rotated reference frame. Consider a beam element initially oriented at an angle $\phi_0$ with respect to the $X$ −axis of the global reference frame. This is the reference configuration where pretensed structure is assumed to be in static equilibrium. The reference and deformed configurations of a beam element are shown in Supplementary Figure 12. The reference configuration is denoted by the nodal coordinates of the end nodes $a$ and $b$ as $\mathbf{X}_a$ and
$\mathbf{X}_b$, while the deformed configuration is identified with nodal coordinates $\mathbf{x}_a = \mathbf{X}_a + \mathbf{u}_a$ and $\mathbf{x}_b = \mathbf{X}_b + \mathbf{u}_b$, where $\mathbf{u}_a$ and $\mathbf{u}_b$ are the nodal displacements. A local co-rotating reference frame $\{X_l, Y_l\}$ is introduced for a given deformed configuration (Supplementary Figure 12) to remove the rigid body motions in the local reference frame.

In this local corotated frame, the transverse nodal displacements are zero and the linearized beam response is given in the following form using Eq. (5.1) as

$$\begin{bmatrix}\widehat{M}_a\\ \widehat{M}_b\end{bmatrix} = \begin{bmatrix}ks & ksc\\ ksc & ks\end{bmatrix}\begin{bmatrix}\hat{\theta}_a - \psi\\ \hat{\theta}_b - \psi\end{bmatrix} \tag{5.4}$$

where $\widehat{M}_a$ and $\widehat{M}_b$ are local end moments, and $\hat{\theta}_a$ and $\hat{\theta}_b$ are the nodal rotations in the local frame.

The angle made by the line connecting the end nodes of an element in the deformed configuration is denoted by $\phi$. The local nodal relations are then obtained as

$$\hat{\theta}_a = \theta_a + \phi_0 - \phi \tag{5.5}$$

$$\hat{\theta}_b = \theta_b + \phi_0 - \phi \tag{5.6}$$

where $\theta_a$ an $\theta_b$ are the nodal rotations in the global frame. The axial response of in the corotated frame is given as

$$\hat{F} = F_0 + \frac{EA}{L_0}(L - L_0) = F_0 + \frac{EA}{L_0}\hat{u}_l, \tag{5.7}$$

where $L$ and $L_0$ are the lengths of beams in deformed and reference configurations. The change in length of the element is denoted by $\hat{u}_l = L - L_0$.

A relation can be established between the local and global degrees of freedom after expressing the variation of $\hat{u}_l$, $\hat{\theta}_a$, and $\hat{\theta}_b$ as follows:

$$\begin{bmatrix}\delta\hat{u}_l\\ \delta\hat{\theta}_a\\ \delta\hat{\theta}_b\end{bmatrix} = \begin{bmatrix}-\cos\phi & -\sin\phi & 0 & \cos\phi & \sin\phi & 0\\ -\sin\phi/L & \cos\phi/L & 1 & \sin\phi/L & -\cos\phi/L & 0\\ -\sin\phi/L & \cos\phi/L & 0 & \sin\phi/L & -\cos\phi/L & 1\end{bmatrix}\begin{bmatrix}\delta u_a\\ \delta v_a\\ \delta\theta_a\\ \delta u_b\\ \delta v_b\\ \delta\theta_b\end{bmatrix}. \tag{5.8}$$

The vector of local degrees of freedom is given by $d_l = \left[\hat{u}_l, \hat{\theta}_a, \hat{\theta}_b\right]^T$ and the vector of degree of freedom in the global reference frame is given by $d = [u_a, v_a, \theta_a, u_b, v_b, \theta_b]^T$, then the mapping shown above can be written as $\delta\mathbf{d}_l = \mathbf{B}\delta\mathbf{d}$. The vector of internal force in the local frame of reference is denoted by $\mathbf{f}_l = \left[\hat{F}, \widehat{M}_a, \widehat{M}_b\right]^T$.

The virtual work principle for the beam element is expressed in the following form

$$\mathbf{f}_l \cdot \delta\mathbf{d}_l - \mathbf{f}_e \cdot \delta\mathbf{d} = 0, \tag{5.9}$$

where $\mathbf{f}_e$ is the constant external nodal force vector in the global frame.

Note that the internal mechanical work is associated only with the deformations in the local frame. Rigid body motions do not contribute to the internal work. Using the relation between $\delta\mathbf{d}_l$ and $\delta\mathbf{d}$ we get the virtual work statement in the global reference frame as follows

$$\left(\mathbf{B}^T\mathbf{f}_l - \mathbf{f}_e\right) \cdot \delta\mathbf{d} = \mathbf{0} \tag{5.10}$$

Using the arbitrariness of $\delta\mathbf{d}$ we arrive at the force balance equations as

$$\mathbf{B}^T\mathbf{f}_l - \mathbf{f}_e = \mathbf{0} \tag{5.11}$$

where the geometric nonlinearity is embedded in the mapping $\mathbf{B}$ which depends on the current configuration. The governing equations for the entire network of nanofibers are obtained by assembling together the discrete beam elements forming the network. The assembled system of nonlinear equations is given as $\mathbf{R}(\mathbf{u},\mathbf{f}) = \mathbf{0}$ where $\mathbf{u}$ is the assembled vector of all nodal degrees of freedom and $\mathbf{f}$ is the assembled vector of all nodal forces, both expressed in the global frame.

The resulting nonlinear system of equations can be solved using the Newton-Raphson method. The consistent tangent stiffness matrix is obtained element by taking variation of the force balance equation Eq. (5.11),

$$\delta(\mathbf{B}^T\mathbf{f}_l) = \delta\mathbf{B}^T\mathbf{f}_l + \mathbf{B}^T\delta\mathbf{f}_l. \tag{5.12}$$

where $\delta f_l$ is obtained from the local relations for end moments and forces as follows

$$\delta\mathbf{f}_l = \begin{bmatrix}\delta\hat{F}\\ \delta\widehat{M}_a\\ \delta\widehat{M}_b\end{bmatrix} = \begin{bmatrix}EA/L_0 & 0 & 0\\ 0 & ks & ksc\\ 0 & ksc & ks\end{bmatrix}\begin{bmatrix}\delta\hat{u}_l\\ \delta\hat{\theta}_a\\ \delta\hat{\theta}_b\end{bmatrix} = \mathbf{C}_b\delta\mathbf{d}_l = \mathbf{C}_b\mathbf{B}\delta\mathbf{d} \tag{5.13}$$

where $\mathbf{C}_b$ is the local stiffness matrix of the pre-tensed Euler-Bernoulli beam element. The second term in Eq. (5.12) is expressed as

$$\mathbf{B}^T\delta\boldsymbol{f}_l = (\mathbf{B}^T\mathbf{C}_b\mathbf{B})\delta\boldsymbol{d} = \mathbf{K}_l\delta\boldsymbol{d}$$

where

$$\mathbf{K}_l = \mathbf{B}^T\mathbf{C}_b\mathbf{B} \tag{5.14}$$

is the beam tangent stiffness matrix expressed in the global reference frame.

The first term in Eq. (5.12) gives the geometric contribution to the tangent stiffness matrix expressed as

$$\delta\mathbf{B}^T\mathbf{f}_l = \delta\mathbf{B}_1\hat{F} + \delta\mathbf{B}_2\widehat{M}_a + \delta\mathbf{B}_3\widehat{M}_b \tag{5.15}$$

where $\mathbf{B}_1$, $\mathbf{B}_2$, and $\mathbf{B}_3$ are first, second, and third row vectors of $B$, respectively. The variations of these row vectors are given as follows:

$$\begin{aligned}\delta\mathbf{B}_1 &= \delta[-\cos\phi \quad -\sin\phi \quad 0 \quad \cos\phi \quad \sin\phi \quad 0]^T\\ &= [\sin\phi \;\; -\cos\phi \quad 0 \quad -\sin\phi \quad \cos\phi \quad 0]^T\delta\phi = \mathbf{z}\delta\phi = \frac{1}{L}\mathbf{z}\mathbf{z}^T\delta\mathbf{d}\end{aligned} \tag{5.16}$$

and

$$\delta\mathbf{B}_2 = \delta\mathbf{B}_3 = \frac{1}{L^2}(\mathbf{r}\mathbf{z}^T + \mathbf{z}\mathbf{r}^T)\delta\mathbf{d} \tag{5.17}$$

where

$$\mathbf{r} = [-\cos\phi \quad -\sin\phi \quad 0 \quad \cos\phi \quad \sin\phi \quad 0]^T, \tag{5.18}$$

$$\mathbf{z} = [\sin\phi \quad -\cos\phi \quad 0 \quad -\sin\phi \quad \cos\phi \quad 0]^T. \tag{5.19}$$

Using these expressions, Eq. (5.15) is expressed as

$$\delta\mathbf{B}^T\mathbf{f}_l = [\frac{\hat{F}}{L}\mathbf{z}\mathbf{z}^T + \frac{\widehat{M}_a + \widehat{M}_b}{L^2}(\mathbf{r}\mathbf{z}^T + \mathbf{z}\mathbf{r}^T)]\delta\mathbf{d} = \mathbf{K}_g\delta\mathbf{d} \tag{5.20}$$

where $\boldsymbol{K}_{\mathrm{g}}$ is the geometric tangent stiffness contribution. The variational consistent tangent stiffness matrix of the corotational beam element is given using Equations (5.23) and (5.29) as $\mathbf{K}_t = \mathbf{K}_l + \mathbf{K}_g$. The Newton-Raphson method with arc-length control is used to solve the forward problem.[28]

#### 5.2 Inverse problem formulation

The inverse problem is formulated as a constrained minimization problem with regularization.[29,30] The objective function is comprised of two terms, (1) error terms quantifying the difference between measured nanonet deformations and the computed deformations from the computational model, and (2) a regularization term that penalizes large values of the estimated forces. The governing force balance equations $\mathbf{R}(\mathbf{u}, \mathbf{f}) = \mathbf{0}$ are imposed as a constraint on the error minimization problem. The formulation of error terms is discussed first.

##### 5.2.1 Matching displacements at grid intersections

For the nanofiber network described with Euler-Bernoulli beams, every point of the fiber network has 3 degrees of freedom $\{u, v, \theta\}$ for in-plane deformations. In the current experimental setup, we are not tracking individual material points on the nanofibers as it would require infusing nanofibers with fiduciary markers or a speckle pattern. We circumvent this by employing grid intersections formed by nanofibers that act as natural markers to track the deformation of the grid. The measured displacements at the grid intersections provide only partial information of the full deformation field. We account for the partial access to the full deformation fields in our formulation.

The column vector collecting $n$ nodal degrees of freedom of the discrete finite elemental model is denoted by $u \in R^n$. However, nodal displacements are measured only at grid intersections. Moreover, nodal rotations are not measured as they are challenging to extract accurately from fiber images. Therefore, a linear map $\mathbf{T}_u: R^n \to R^m$ is defined to retrieve the model predicted displacements at the measurement locations $\hat{\mathbf{u}}_{\mathrm{m}} \in R^m$ from $u$ using $\hat{\mathbf{u}}_m = \mathbf{T}_u\mathbf{u}$. The experimental measurements are stored in a vector $\mathbf{u}_m \in R^m$. The error between the model data ($\mathbf{u}_m$) and the measured data ($\hat{\mathbf{u}}_m$) is defined as

$$\mathrm{E_m(u)} = \frac{1}{2}\|\mathbf{u}_m - \hat{\mathbf{u}}_m\|^2 = \frac{1}{2}\|\mathbf{u}_m - \mathbf{T}_u\mathbf{u}\|^2. \tag{5.21}$$

##### 5.2.2 Image-based matching of deformed fiber shapes

The fluorescent images of deformed fibers are obtained from phase images using the pix2pix model (see methods and Supplementary Note 1). Although these images capture deformed shapes of fibers, the available data is not in the form of displacements at specific material points *i.e.* we do not have a deformation map. Thus we introduce an image intensity based error function, inspired by the active-contour method (or snakes) used in line/edge detection,[31–33] to match the computational mesh with deformed fiber shapes. The grayscale fluorescent image is converted into a binary nanofiber mask where pixels belonging to nanofibers are assigned a value 0 and the background is set to 1. This step ensures that there are no differences between pixel intensity values for different fibers. The binary image is then smoothed using Gaussian filtering to avoid sharp gradients. The error function for fiber shape matching is given in terms of the image intensity at the current position $\boldsymbol{x}$ of a node denoted by $I(\boldsymbol{x})$,

$$\mathrm{E_l}(\mathbf{u}) = \frac{1}{2}\sum_{\mathrm{i}} I(x_i)^2 \tag{5.22}$$

where $x_i$ is the current position of the $i$ −th node and summation is over all nodes used for fiber shape matching. Bicubic interpolation scheme is used to obtain image intensity and gradients at any given position $x_i$ on the image. Consequently, this error function drives the computational mesh to overlap with the deformed fibers.

In the classical active-contour model for image segmentation,[31–33] the line energy term is accompanied by a regularizing in the form of stretching and bending energy contributions to maintain continuity and smoothness of the contour. Here we can use the corotational beam formulation to define the stretching and bending energies for the beam elements of the FE mesh. The elastic regularization energy is associated with the deformation in the local corotated frame of an element (Supplementary Figure 12) and is given as

$$E_{er}(\mathbf{u}) = \sum_{e} \frac{1}{2}\left(\hat{F}\hat{u}_l + \widehat{M}_a\hat{\theta}_a + \widehat{M}_b\hat{\theta}_b\right)_e. \tag{5.23}$$

where the sum is over all elements $e$ that participate in the fiber shape matching process. For notational convenience, we denote the combined active-contour energy contribution for the fiber shape matching as $E_{ac}(\mathbf{u}) = w_l E_l(\mathbf{u}) + w_{er} E_{er}(\mathbf{u})$. where the weights $w_l$ and $w_{er}$ are defined to adjust the relative contribution of a given term in the minimization process. Although the elastic penalty term $E_{er}(\mathbf{u})$ has a regularizing effect, more direct way of regularizing inverse problems is through the penalty on the magnitude of the unknown nodal forces, which is discussed next. Moreover, unlike the segmentation methods, we do not really need $E_{er}(\boldsymbol{u})$ to restrict the deformation of the mesh as the error minimization is constrained by the nanonet mechanics. In fact, we observe that $w_{er} = 0$ works well for solving this inverse problem. Unless stated otherwise, we use $w_{er} = 0$ for all analysis presented here and in the main text.

##### 5.2.3 Regularization

We define the vector of unknowns (parameters) $\mathbf{f} \in R^p$ that in general is the set of all model parameters to be estimated. For cNFM, it represents the set of unknown nodal forces. A linear map $\mathbf{T}_f: R^p \to R^n$ is defined to get the nodal (external) forces vector $\mathbf{f}_e = \mathbf{T}_f\mathbf{f}$ from $\mathbf{f}$. The nonlinear forward problem is given in terms of the residual force vector as $\mathbf{R}(\mathbf{u}, \mathbf{f}) = \mathbf{0}$.

The inverse problem solution is regularized by penalizing the norm of the nodal forces $\mathbf{f}$ *i.e.* using Tikhonov regularization.[34] The regularized objective function of the inverse problem is given as

$$\mathcal{W}(\mathbf{f}) = \frac{w_m}{2}\|\mathbf{u}_m - \mathbf{T}_u\mathbf{u}(\mathbf{f})\|^2 + E_{ac}(\mathbf{u}(\mathbf{f})) + \frac{1}{2}\beta^2\|\mathbf{f}\|^2 \tag{5.24}$$

where β, in the units of µm/nN, is the regularization parameter and $w_m$ is the weighing coefficient for the nodal error term. Note that $u$ is not an independent variable, but rather a function of $t$ since $u$ is obtained by solving the force balance system of equations $\mathbf{R}(\mathbf{u}, \mathbf{f}) = \mathbf{0}$. Taken together, the inverse problem is stated as

$$\mathbf{f} = \mathrm{argmin}_{\mathbf{f}}\left(\frac{w_m}{2}\|\mathbf{u}_m - \mathbf{T}_u\mathbf{u}(\mathbf{f})\|^2 + E_{ac}(\mathbf{u}(\mathbf{f})) + \frac{1}{2}\beta^2\|\mathbf{f}\|^2\right). \tag{5.25}$$

##### 5.2.4 Solving the error minimization problem

*Linear forward problem.* If the force balance equations were linear *i.e.* of the form $\mathbf{R}(\mathbf{u},\mathbf{f}) = \mathbf{K}\mathbf{u} - \mathbf{T}_f\mathbf{f}$, where $\mathbf{K}$ is a constant stiffness matrix of the system, then the solution vector is directly expressed as $\mathbf{u}(\mathbf{f}) = \mathbf{K}^{-1}\mathbf{T}_f\mathbf{f}$. The minimization problem can be easily solved by substituting $\mathbf{u}(\mathbf{f})$, with the only catch that this approach would require an explicitly storage of the dense matrix $\mathbf{K}^{-1}$ to evaluate the hessian of the objective function $\mathcal{W}(\mathbf{f})$. Alternatively, we can pose the inverse problem as a constrained minimization problem using the following Lagrangian

$$\mathcal{L}(\mathbf{u},\mathbf{f},\boldsymbol{\lambda}) = \frac{w_m}{2}\|\mathbf{u}_m - \mathbf{T}_u\mathbf{u}(\mathbf{f})\|^2 + E_{ac}(\mathbf{u}) + \frac{1}{2}\beta^2\|\mathbf{f}\|^2 - \boldsymbol{\lambda}^T\mathbf{R}(\mathbf{u},\mathbf{f}) \tag{5.26}$$

where the governing equations are imposed as a constraint through the Lagrangian multipliers $\boldsymbol{\lambda} \in R^n$. Then for a linear forward problem we have

$$\mathcal{L}(\mathbf{u},\mathbf{f},\boldsymbol{\lambda}) = \frac{w_m}{2}\|\mathbf{u}_m - \mathbf{T}_u\mathbf{u}(\mathbf{f})\|^2 + E_{ac}(\mathbf{u}) + \frac{1}{2}\beta^2\|\mathbf{f}\|^2 - \boldsymbol{\lambda}^T(\mathbf{K}\mathbf{u} - \mathbf{T}_f\mathbf{f}) \tag{5.27}$$

The solution to the inverse problem is given as

$$\{\mathbf{u},\mathbf{f},\boldsymbol{\lambda}\} = \text{argmin}_{\mathbf{u},\mathbf{f}}\text{argmax}_{\boldsymbol{\lambda}}\mathcal{L}(\mathbf{u},\mathbf{f},\boldsymbol{\lambda}). \tag{5.28}$$

The corresponding linear system of equations of the saddle-point problem are obtained as

$$\frac{\partial\mathcal{L}}{\partial\mathbf{u}} = w_m\mathbf{T}_u^T(\mathbf{T}_u\mathbf{u} - \mathbf{u}_m) + \frac{\partial E_{ac}(\mathbf{u})}{\partial\mathbf{u}} - \mathbf{K}^T\boldsymbol{\lambda} = \mathbf{0} \tag{5.29}$$

$$\frac{\partial\mathcal{L}}{\partial\boldsymbol{\lambda}} = \mathbf{K}\mathbf{u} - \mathbf{T}_f\mathbf{f} = \mathbf{0} \tag{5.30}$$

$$\frac{\partial\mathcal{L}}{\partial\mathbf{f}} = \mathbf{T}_f^T\boldsymbol{\lambda} + \beta^2\mathbf{f} = \mathbf{0}. \tag{5.31}$$

This linear system can be readily solved to obtain the inverse solution for a given regularization parameter β. Such a linear forward problem formulation can be used if the fiber deflections are sufficiently small. But otherwise, the nonlinear forward problem needs to be considered as discussed next.

*Nonlinear forward problem.* For large deflections of fibers the forward problem is nonlinear and the saddle-point problem in Eq. (5.27) can no longer be solved directly. Instead, the extremization problem is solved in the nonlinear case using an iterative gradient-based approach. Note that evaluating the gradient of the regularized error function $\mathcal{W}(\mathbf{f})$ is challenging given the implicit dependence of $\mathbf{u}$ on $\mathbf{f}$ through the nonlinear governing equations for the forward problem. This hurdle is overcome by posing the inverse problem as a constrained minimization problem using the Lagrangian defined in Eq. (5.27). Differential of the Lagrangian is given using the chain rule as

$$\mathrm{d}\mathcal{L} = \frac{\partial\mathcal{L}}{\partial\mathbf{u}}\mathrm{d}\mathbf{u} + \frac{\partial\mathcal{L}}{\partial\mathbf{f}}\mathrm{d}\mathbf{f} + \frac{\partial\mathcal{L}}{\partial\boldsymbol{\lambda}}\mathrm{d}\boldsymbol{\lambda} \tag{5.32}$$

where

$$\frac{\partial\mathcal{L}}{\partial\mathbf{u}} = w_m\mathbf{T}_u^T(\mathbf{T}_u\mathbf{u} - \mathbf{u}_m) + \frac{\partial E_{ac}(\mathbf{u})}{\partial\mathbf{u}} - \boldsymbol{\lambda}^T\frac{\partial\mathbf{R}(\mathbf{u},\mathbf{f})}{\partial\mathbf{u}} \tag{5.33}$$

$$\frac{\partial \mathcal{L}}{\partial \mathbf{f}} = -\boldsymbol{\lambda}^T \frac{\partial \mathbf{R}(\mathbf{u}, \mathbf{f})}{\partial \mathbf{f}} + \beta^2 \mathbf{f} \tag{5.34}$$

$$\frac{\partial \mathcal{L}}{\partial \boldsymbol{\lambda}} = \mathbf{R}(\mathbf{u}, \mathbf{f}) \tag{5.35}$$

For a given $\mathbf{f}$, the forward problem solution $\mathbf{u}$ can be obtained by solving $\mathbf{R}(\mathbf{u}, \mathbf{f}) = \mathbf{0}$ so that $\partial \mathcal{L}/\partial \boldsymbol{\lambda} = 0$. Then, the Lagrangian multipliers $\boldsymbol{\lambda}$ are obtained from Eq. (5.33) resulting in the following equation

$$\frac{\partial \mathcal{L}}{\partial \mathbf{u}} = w_m \mathbf{T}_u^T (\mathbf{T}_u \mathbf{u} - \mathbf{u}_m) + \frac{\partial E_{ac}(\mathbf{u})}{\partial \mathbf{u}} - \boldsymbol{\lambda}^T \frac{\partial \mathbf{R}(\mathbf{u}, \mathbf{f})}{\partial \mathbf{u}} = \mathbf{0} \tag{5.36}$$

where $\partial \mathbf{R}/ \partial \mathbf{u}$ is the consistent tangent matrix of the nonlinear equations of the forward problem. It is evaluated using the solution to the forward problem for a given $\mathbf{f}$. Under these conditions, since $\boldsymbol{\lambda}$ and $\mathbf{u}$ are given, the gradient of the Lagrangian with respect to $\mathbf{f}$ is expressed from Eq. (5.34) as

$$\frac{\mathrm{d}\mathcal{L}}{\mathrm{d}\mathbf{f}} = -\boldsymbol{\lambda}^T \frac{\partial \mathbf{R}(\mathbf{u}, \mathbf{f})}{\partial \mathbf{f}} + \beta^2 \mathbf{f} \tag{5.37}$$

When $\mathbf{R}(\mathbf{u}, \mathbf{f}) = \mathbf{0}$, $\mathcal{W}(\mathbf{f}) = \mathcal{L}(\mathbf{u}, \mathbf{f}, \boldsymbol{\lambda})$, and furthermore when $\partial \mathcal{L}/\partial \boldsymbol{u} = \mathbf{0}$ we also have that

$$\frac{\mathrm{d}\mathcal{W}}{\mathrm{d}\mathbf{f}} = \frac{\mathrm{d}\mathcal{L}}{\mathrm{d}\mathbf{f}}.$$

Therefore, gradient of the objective function can be efficiently obtained by solving the additional adjoint problem in Eq. (5.36).

In summary, we minimize the regularized error function using a gradient-based optimization approach where the gradient is evaluated using the adjoint-set method.[35,36]

#### 5.3 Effective material model of nanofiber networks

The experimental images typically capture a crosshatch network area roughly containing 20 by 20 grid intersections for square grids. The computational FE model of the nanonet is built by generating a mesh by connecting these intersecting grid points (see Supplementary Note 4). The nanonet substrate, however, is much larger in size (~ 4 mm × 4 mm) than the captured window size. The boundary conditions for the computational model need to be carefully implemented to replicate this physical setup. Fixed boundary conditions on the boundary nodes of the imaged region are clearly not appropriate as these will overly constrain the network. More importantly, as the elasticity Green's function decay is slower for planar elasticity (compared to thick gels modeled as elastic half spaces[29]), imposing fixed boundary conditions will result in overestimation of cell forces due to the increased apparent stiffness from such boundary conditions. To avoid such stiff boundary conditions, the computational grid is surrounded with a nanofiber network with similar spacing and connectivity as the imaged domain to mimic the experimental setup.

For the forces imposed by the cells, fiber deformations at a distance of 2-3 cell diameters away from the cell are sufficiently small. In this region away from the cells, due to small deflections the geometrical nonlinearities do not contribute significantly in the beam response. A linear approximation for the beam mechanics can be employed in regions surrounding the imaged network. The modeling of the linear pre-tensed beam grid network is further simplified by homogenizing the response over a unit cell of the grid. This strategy sufficiently reduces the computational resources required to represent a large nanonet surrounding the imaged region.

To this end, the periodic network of pre-tensed Euler-Bernoulli beams with a rectangular unit cell (Supplementary Figure 13) is homogenized as a Cosserat (micropolar) continuum.[27,37] The constitutive relations of the resulting orthotropic micropolar media are given as

$$
\begin{aligned}
\sigma_{11} &= E_A \varepsilon_{11} \frac{L_A}{L_B} + \frac{F_0}{L_B} \\
\sigma_{22} &= E_B \varepsilon_{22} \frac{L_B}{L_A} + \frac{F_0}{L_A} \\
\sigma_{12} &= \frac{1}{L_A L_B} \left( k_A s_A'' \varepsilon_{12} - F_0 L_A \phi \right) \\
\sigma_{21} &= \frac{1}{L_A L_B} \left( k_B s_B'' \varepsilon_{21} + F_0 L_B \phi \right) \\
m_{31} &= k_A s_A \phi_{,1} \frac{L_A}{L_B} \\
m_{32} &= k_B s_B \phi_{,2} \frac{L_A}{L_B}
\end{aligned}
\tag{5.38}
$$

where $\sigma_{ij}$ are the planar non-symmetric Cauchy stress components and $m_{ij}$ are the polar moment stress components with $i, j \in \{1, 2\}$. The subscripts $A$ and $B$ denote properties of fibers in horizontal and vertical directions, respectively. For instance, $L_A$ is the spacing between fibers in the horizontal direction and $L_B$ is the spacing in the vertical direction. The structural stiffness of beams is given as $E_A = EA/L_A$ and $E_B = EA/L_B$, where $E$ is the Young's modulus and $A$ is the cross-section area of fibers. The cross-section area of vertical and horizontal fibers remains the same. $\{s_A, c_A\}$ and $\{s_B, c_B\}$ are the stability functions defined in Eqs. (5.2) and (5.3), while $s_A'' = 2s_A(1 + c_A) + F_0 L_A / k_A$ and similarly $s_B'' = 2s_B(1 + c_B) + F_0 L_B / k_B$. The second moment of areas of the horizontal and vertical beams are given by $I_A$ and $I_B$, and furthermore, $k_A = E_A I_A / L_A$ and $k_B = E_B I_B / L_B$.

#### 5.4 Finite elemental formulation of Cosserat media

The strong form of the 3D balance of linear momentum for a micropolar media under the assumption of small strains and rotations is given as

$$
\sigma_{ji,j} + b_i = 0 \tag{5.39}
$$

where repeated indices imply summation (Einstein's summation notation) and $i, j \in \{1,2,3\}$. The balance of angular momentum is expressed as

$$
m_{ji,j} + e_{ijk} \sigma_{jk} + g_i = 0. \tag{5.40}
$$

Here $\sigma_{ij}$ are the components of the non-symmetric Cauchy stress tensor, $m_{ij}$ are the components of the moment or couple stress tensor, $e_{ijk}$ is the permutation symbol, $b_i$ is the body force, and $g_i$ is body moment contribution. The kinematics of a micropolar continuum are described by a displacement field $u$ and a rotation field $\phi$. The infinitesimal strain tensor is defined as

$$
\varepsilon_{ij} = u_{j,i} - e_{kij} \phi_k \tag{5.41}
$$

Since we intend to model a planar lattice of nanofibers, these 3D balance equations need to be expressed under planar conditions with plane-stress assumption. In this simplification, only in-plane displacements $u_1, u_2$ and rotations about the out-of-plane axis $\phi = \phi_3$ are relevant. The out-of-plane stress components are assumed to be zero, $\sigma_{33} = \sigma_{13} = \sigma_{23} = \sigma_{31} = \sigma_{32} = 0$. Furthermore, for notational convenience we use $m_1 = m_{13}$ and $m_2 = m_{23}$, which are the only relevant components for the couple stress for planar analysis and $g = g_3$ is the only relevant body moment component. The balance equations and strain definition for planar analysis are given as

$$\sigma_{ji,j} + b_i = 0, \tag{5.42}$$

$$m_{j,j} + e_{3ji}\sigma_{ji} + g = 0, \tag{5.43}$$

$$\varepsilon_{ij} = u_{j,i} - e_{3ij}\phi. \tag{5.44}$$

The weak form of the equations is obtained by multiplying the linear momentum balance equation with a vectorial admissible test function $\delta u$ and multiplying the angular momentum balance with a scalar admissible test function $\delta\phi$ and integrating over the domain. Using integration by parts followed by application of the Green-Gauss theorem we obtain

$$\int_S \sigma_{ji}\delta u_{i,j}\ dS - \int_S b_i\delta u_i\ dS - \int_{\partial S} \sigma_{ji}n_j\delta u_i\ dl = 0 \tag{5.45}$$

$$\int_S m_j\delta\phi_{,j}\ dS + \int_S e_{3ji}\sigma_{ji}\delta\phi\, dS - \int_S g\ \delta\phi\, dS - \int_{\partial S} m_j n_j\delta\phi\ dl = 0 \tag{5.46}$$

Adding the equations above and using $\delta\varepsilon_{ij} = \delta u_{j,i} - e_{3ij}\delta\phi$ we get

$$\int_S \sigma_{ij}\delta\varepsilon_{ij}\ dS + \int_S m_j\delta\phi_{,j}\ dS = \int_S b_i\delta u_i\ dS + \int_{\partial S_t} \sigma_{ji}n_j\delta u_i\ dl + \int_S g\ \delta\phi\, dS + \int_{\partial S_t} m_j n_j\delta\phi\ dl. \tag{5.47}$$

The stress and strain components are rearranged in the following matrix form for convenience

$$\varepsilon = [\varepsilon_{11} \quad \varepsilon_{22} \quad \varepsilon_{12} \quad \varepsilon_{21} \quad \phi_{,1} \quad \phi_{,2}]^T, \tag{5.48}$$

$$\sigma = [\sigma_{11} \quad \sigma_{22} \quad \sigma_{12} \quad \sigma_{21} \quad m_1 \quad m_2]^T. \tag{5.49}$$

The left-hand side of the weak form Eq. (5.47) is then written as $\int_S \sigma^T\delta\varepsilon\ dS$. The discrete formulation is obtained using the Galerkin method with piecewise linear interpolation of the displacement and rotation fields expressed as

$$u_i = \sum_{I\in<E>} N_I u_{iI} \tag{5.50}$$

$$\phi = \sum_{I\in<E>} N_I \phi_I \tag{5.51}$$

where $u_{iI}$ are the nodal displacements and $\phi_I$ represent the nodal rotation for a node $I$ belonging to an element $E$ with linear shape functions $N_I$. The constitutive relation for the homogenized rectangular nanofiber network given in Eq. (5.38) is expressed in the matrix notation as $\sigma = \mathbf{D}\varepsilon + \mathbf{E}u + \sigma_0$, where $u = [u_1 \quad u_2 \quad \phi]^T$ and $\mathbf{D}$ is the stiffness matrix, while $\mathbf{E}u$ and $\sigma_0$ are stress contributions due to the pretension in the nanofibers. The resulting system of linear equations is written as the residual vector between internal and external forces at nodes $\mathbf{R}_p(\mathbf{u}_p, \mathbf{f}_p) = \mathbf{0}$ where the subscript $p$ refers to the surrounding domain about the imaged region. This linear system of equations is coupled to the beam problem through the overlapping degrees of freedom at the interface of these two domains. The combined system of equations is solved together for the forward problem. As the consistent tangent stiffness for the linear problem remains constant, it is calculated only once during the solution process. The surrounding domain is discretized using triangular elements of the size of the grid spacing close to the discrete domain (see Supplementary Figure 9e).

#### 5.5 Validating the inverse formulation

Here we present a systematic validation of the inverse formulation using the input data generated by solving the forward problem with a known force pattern $\mathbf{f}_f$. The solution of the forward problem $\mathbf{u}_f$ provides the measurement data at the desired measurement locations using $\mathbf{u}_m = \mathbf{T}_u\mathbf{u}_f$, where $\mathbf{T}_u$ is the mapping from the full solution vector to the desired measurement locations. To simulate errors in experimental measurements, we introduce noise to the measured data such that

$$\mathbf{u}_m = \mathbf{T}_u\mathbf{u}_f + \mathbf{u}_e \tag{5.52}$$

where $\mathbf{u}_e \in R^m$ is a vector of independent identically distributed random variables following a zero-mean Gaussian distribution with standard deviation $\sigma_e$. The choice of Gaussian distribution is motivated by the actual measured errors that are observed to follow a Gaussian distribution (Supplementary Figure 8). Furthermore, the deformed shape of the network is stored as an image $I$ to be used for the active-contour regularization.

First, a simple example with only two nodal forces is considered as shown in Supplementary Figure 14a. The entire solution (nodal displacements and rotations) without any noise is provided such that $\mathbf{u}_m = \mathbf{u}_f$. In this case the applied forces are fully recovered (Supplementary Figure 14c,d) after solving the inverse problem. Even when only the displacements at the fiber intersections are provided such that $\mathbf{u}_m = \mathbf{T}_u\mathbf{u}_f$ (Supplementary Figure 14b), the inverse solver can fully recover the ground truth (Supplementary Figure 14 c and d). In this example, the minimization process quickly converges to the ground-truth due to lower number of force unknowns compared to the known information in the form of the deformation field $\mathbf{u}_m$.

Next, we performed a similar validation analysis on a more representative radially contractile force pattern as shown in Supplementary Figure 15a. In this case the applied forces are not fully recovered even when the entire solution is provided without any noise *i.e.* $\mathbf{u}_m = \mathbf{u}_f$ (Supplementary Figure 16). This emanates from the slow decay of Green's function on the nanonets which makes the problem ill-posed. Essentially, multiple statically equivalent force patterns can be obtained within the convergence tolerance of the error minimization process. Thus, an exact match between the regularized solution $\mathbf{f}$ and $\mathbf{f}_f$ is not expected when a point-wise comparison is performed as shown in Supplementary Figure 16b. Instead, we compare the sum of forces within a patch of certain radius $r_p$ about a given location to compare $\mathbf{f}$ and $\mathbf{f}_f$. Such locally aggregated comparison with $r_p$ of the order of grid spacing is appropriate when the measured data is provided only at the grid intersections. A large $r_p$ limit of this is comparing the effective contractility of both force patterns (Supplementary Figure 16b,c). See the methods section for the formula to calculate the effective contractility of a force pattern. We conclude that the proposed method can reliably estimate the sum of forces acting within a patch of the size of grid spacing and work especially well in predicting global measures such as contractility. However, full recovery of local forces is not expected even when complete nodal displacements and rotations data is provided with minimal noise.

Next, to mimic the experimental setup, we provide the measurements only at fiber intersections using $\mathbf{u}_m = \mathbf{T}_u \mathbf{u}_f$. Furthermore, we solved the inverse problem for different levels of measurement noise $\sigma_e = \{0.0\ \mu m, 0.2\ \mu m, 0.5\ \mu m\}$ added in $\mathbf{u}_m$. The force reconstruction is first performed without the active-contour regularization *i.e.* $w_l = 0$. The error between the force patterns follows a familiar trend with the regularization parameter β as shown in Supplementary Figure 17. The optimal β that minimizes the error is a function of the noise level $\sigma_e$ (Supplementary Figure 17b). The knee of the L-curve corresponds to a rapid increase in the norm of unknowns $||\mathbf{f}||$ without any appreciable decrease in the mismatch errors quantified as $(\mathbf{u} - \mathbf{u}_m)/\sqrt{N_m}$ (to resemble standard deviation $\sigma_e$) (Supplementary Figure 17a). In fact, this sharp increase on the L-curves occurs precisely when the inverse method attempts to reduce the mismatch errors below the noise level $\sigma_e$. The forces required to match the noise are typically very large resulting in a large $||\mathbf{f}||$.

The solutions obtained at the optimal $\beta$ (at the knee of the L-curve) recovers the force pattern relatively well. However, there is a clear mismatch between deformed fiber shapes since only displacements at grid intersections are provided as seen in Supplementary Figure 17c. Such mismatch between deformed fiber shapes led us to include an image intensity-based term in the inverse formulation to guide the fibers to snap into the valleys of the intensity map $I$ similar to Supplementary Figure 15c. Indeed, including the active-contour regularization with $w_l = 3$ in the minimization process provides a better match with the solution $\mathbf{f}_f$ and $\mathbf{u}_f$ as shown in Supplementary Figure 18b. The overall contractility of the force pattern also matches to within 10\% of the contractility of the target solution $\mathbf{f}_f$ (Supplementary Figure 19). The L-curves obtained for this analysis with $\{w_n = 1, w_l = 3\}$ shown in Supplementary Figure 18a lack a sharp increase in $||\mathbf{f}||$ for small $\beta$ observed before in Supplementary Figure 17a. This is an interesting consequence of using the active-contour regularization as the deformed fibers shapes are restricted to the valleys of the intensity energy landscape controlled by *I*. A sharp increase in $||\mathbf{f}||$ is not feasible as it results in highly distorted fiber shapes. The minimization process saturates after reaching the level of measurement noise.

A similar validation analysis is performed on other representative force patterns which shows qualitatively similar trends in the effectiveness of force recovery (Supplementary Figure 20, Supplementary Figure 21, Supplementary Figure 22, and Supplementary Figure 23). Overall, these validation examples demonstrate that the inverse formulation can recover forces to within 10 % accuracy for representative contractile force patterns. It is also clear from these examples that the regularization through β and $w_l$ is sufficient and an additional regularization through the elastic regularization term $E_{er}$ is not necessary, *i.e.* $w_{er} = 0$. However, elastic regularization could become a useful measure to deal with distorted and broken fiber shapes in fluorescent images.

### 6 Cell Culture, Differentiation Methodology and Additional Results

#### 6.1 Cell Culture and Imaging

The fiber scaffolds were seeded in a glass bottom 6 well plates (MatTek, MA), sterilized using 70% ethanol for 10 min and washed with PBS. Rhodamine tagged Fibronectin (Cytoskeleton Inc, CO) was used to coat the fibers for 1 hour. Bone-marrow derived human mesenchymal stem cells (hMSCs; Lonza, GA) were cultured in supplemented growth medium (PromoCell) in 5% CO2. Cells were trypsinized and added to the scaffolds. 6 well plates with the scaffolds were placed inside a temperature and CO2 controlled incubating Zeiss AxioObserver Z1 microscope. 20x objective was used to obtain phase as well as fluorescent channel images every 5min interval. Zen 3.0 and ImageJ (NIH, MD) was used to process the captured images from the microscope.

Adipogenic differentiation of hMSCs was induced by exposing to cycles of 2 days of adipogenic differentiation medium comprising of high glucose DMEM (Corning, NY), 10% FBS(ATCC), 1 µM dexamethasone (Sigma), 200 µM indomethacin (Sigma), 10 µg/ml insulin, 0.5mM methylisobutyxanthine (Sigma) followed by 1 day of adipogenic maintenance medium comprising of high glucose DMEM, 10 µg/ml insulin. Osteogenic differentiation of hMSCs was induced with osteogenic induction media composed of high glucose DMEM, 10% FBS, 50 µM ascorbic acid-2-phosphate, 5mM β-glycerophosphate and 100 nM dexamethasone with 3 days as a cycle.

#### 6.2 Oil Red O and ALP staining and Quantification

Adipocytes were confirmed through Oil Red O staining for lipids. Briefly, cells were fixed in 10% formalin, followed by DI water rinse and 60% isopropanol, stained with freshly prepared Oil Red O working solution (0.5g Oil Red O power, Sigma in 100ml isopropanol, mixed with DI water in 3:2 ratio) for 15min and rinse with DI water. Color images were obtained using Amscope MU300 camera fitted to Zeiss Axiovert inverted microscope. Semi-quantitative analysis of Oil Red O-stained images was performed as per previously reported methodology[38]. Briefly, red pixels were identified by analyzing the RGB images. A pixel was identified as red if it did not differ by more than a fixed value from the red spectrum (Supplementary Figure 24). Total area covered by the red pixels was then computed.

Alkaline Phosphatase (ALP) was stained for confirming osteogenic differentiation of hMSCs, using Sigma kit 86C as per the manufacturer instructions. Briefly, cells were fixed with citrate-acetone-formaldehyde for 30s, followed by 3X wash with DI water. Next, a solution of FBB alkaline, sodium nitrite and napthol (1:1:1) was added to the cells and stained overnight. The stain was pipetted out and rinsed with water. Color images were obtained using Amscope MU300 camera fitted to Zeiss Axiovert inverted microscope. Semi-quantitative analysis of ALP staining was performed using a previously reported methodology[39,40]. Briefly, manual outlining of cells was performed using ImageJ (NIH, Bethesda) and the mean pixel intensity was obtained from the green slice of RGB image ($I$). Mean intensity ($I_0$ ) of the background was computed using the same cell mask (Supplementary Figure 25). ALP activity was given by the expression $S_{\mathrm{cell}} = \log\frac{I_0}{I}$ where $S_{\mathrm{cell}}$ denotes the number of pixels covering the spread area of cell.

### 8 Supplementary Figures

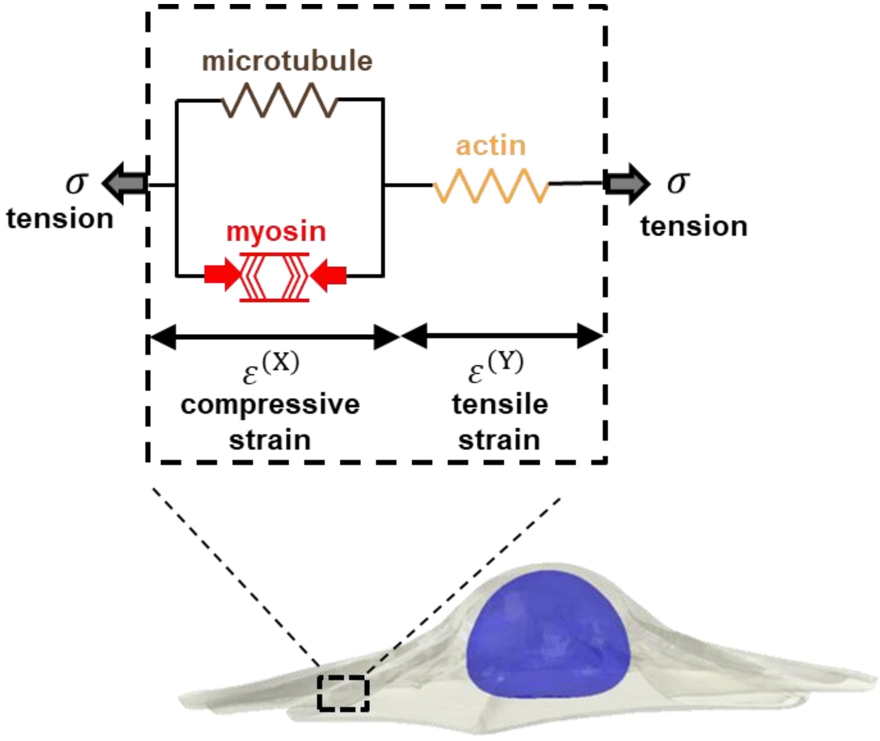

Supplementary Figure 1: Schematic representation of the three-dimensional model of the cytoskeleton. The cell cytoskeleton was modeled as a continuum of representative volume elements (RVEs), each consisting of three components: myosin molecular motors (represented by the active contractile element in red), microtubules (depicted as the passive element in parallel with the myosin element), and actin filaments (shown as the passive element in yellow, connected in series with the other two elements). The contraction force generated by the active contractile element compresses the microtubules, while the remaining force, denoted as $\sigma$, is transmitted to the extracellular matrix through the actin filament network within the cytoskeleton.

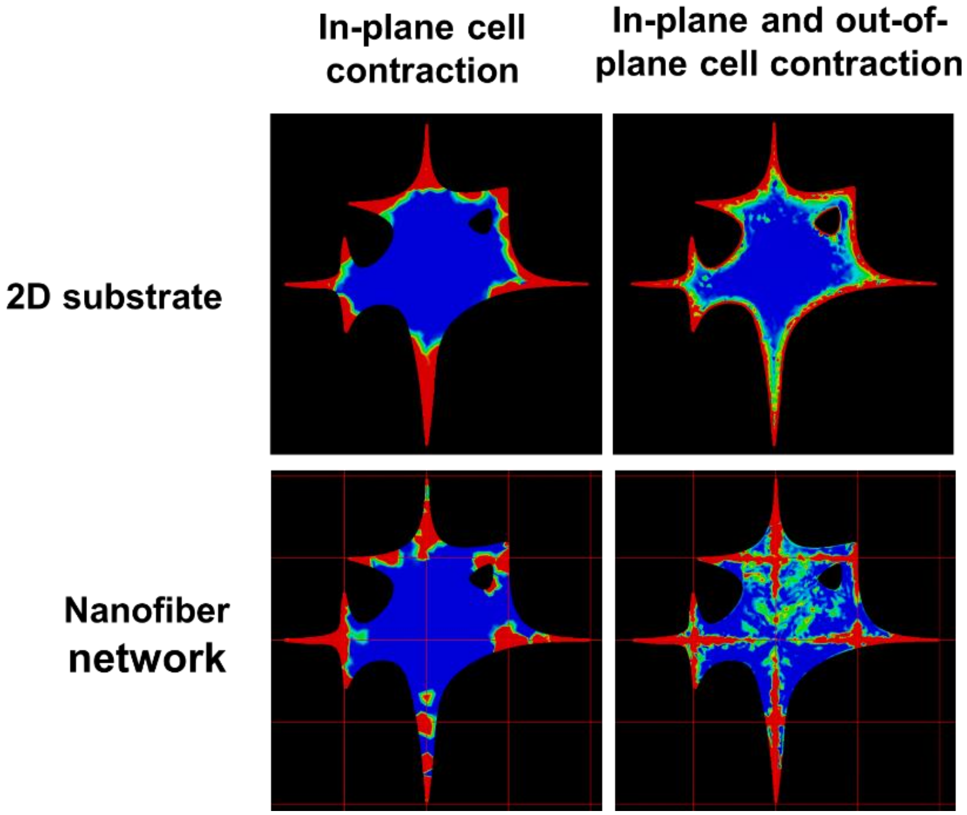

Supplementary Figure 2: Here, we conducted simulations of cell contraction on a two-dimensional (2D) nonfibrous gel and within a nanofiber network. For each scenario, we simulated both in-plane and out-of-plane contractions. In the in-plane contraction, the cell contraction only generated in-plane stress on the focal adhesion layer, which is located between the cell and the matrix. This was achieved by modeling the focal adhesion layer as a shell, a 2D layer without thickness in the z-direction. For the out-of-plane contraction, the focal adhesion layer was modeled with a small thickness, allowing stress generation in the out-of-plane direction as well. Our simulations showed that on a 2D nonfibrous matrix, focal adhesions always formed around the periphery of the cell, regardless of whether the contraction was in-plane or out-of-plane. However, for cells within the nanofiber network, allowing out-of-plane stress resulted in the formation of focal adhesions not only around the cell periphery but also farther away from the cell periphery.

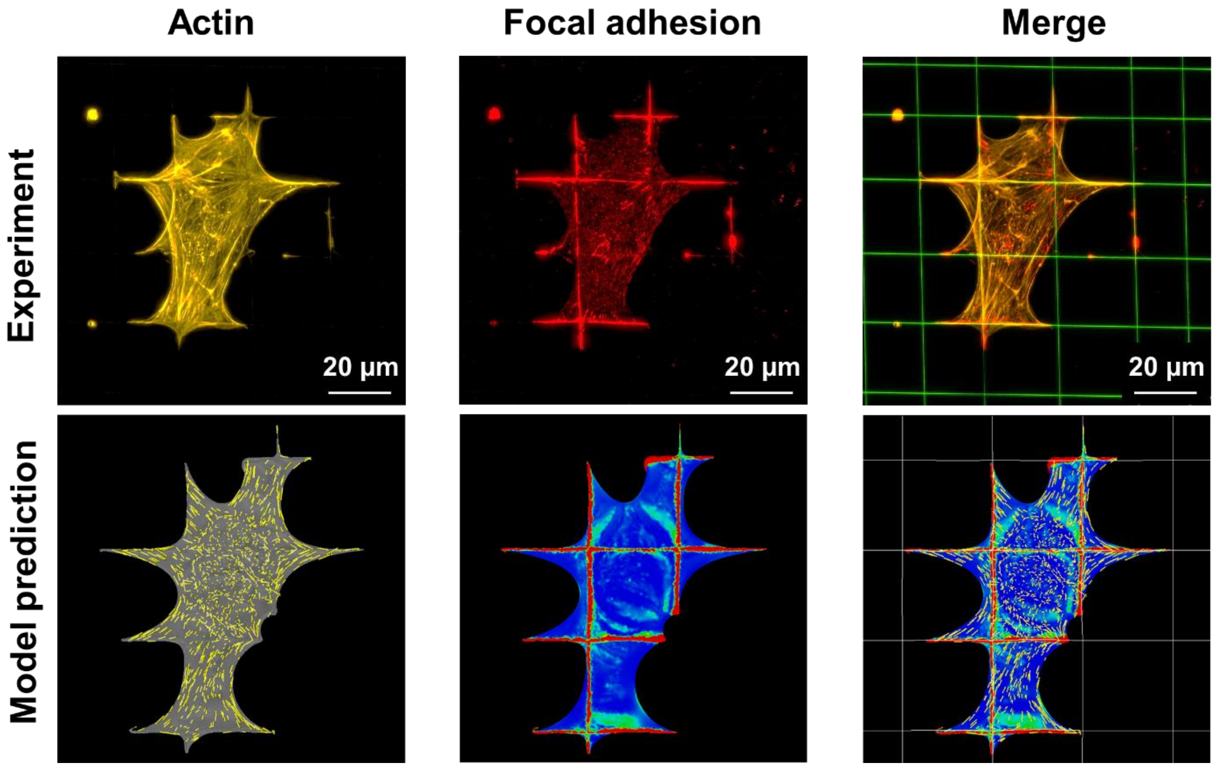

Supplementary Figure 3: Using the out-of-plane contraction described in Supplementary Figure 2, we simulated the contraction of a well-spread representative cell within our nanofiber network as detailed in Figure 1. We validated the predicted actin formation and focal adhesion formation by comparing our simulation results with experimental staining for actin (phalloidin) and paxillin.

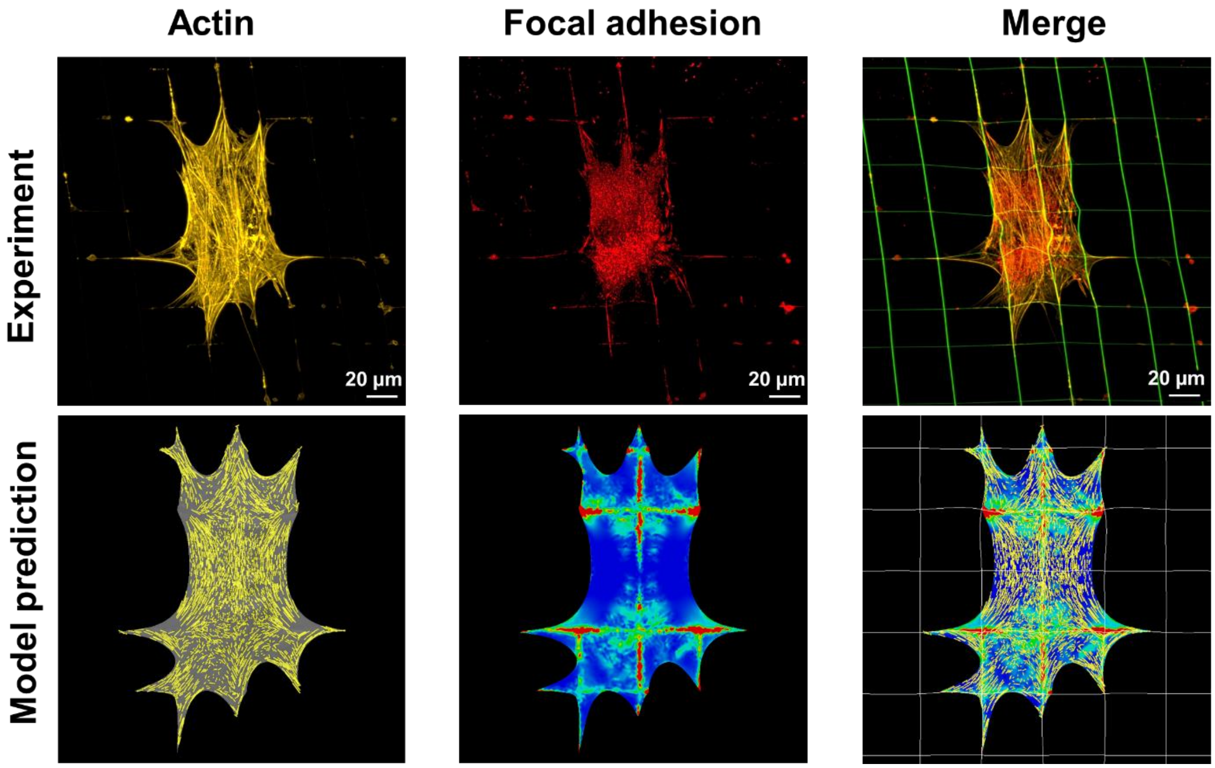

Supplementary Figure 4: Results for representative cells with (A) polarized and (B) non-polarized morphologies.

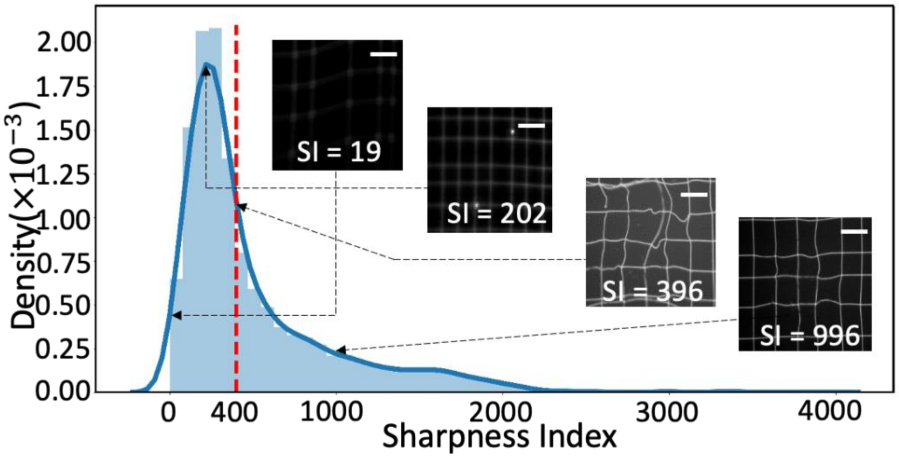

Supplementary Figure 5: Histogram of Sharpness Index of experimentally obtained fluorescent images along with representative image patches.

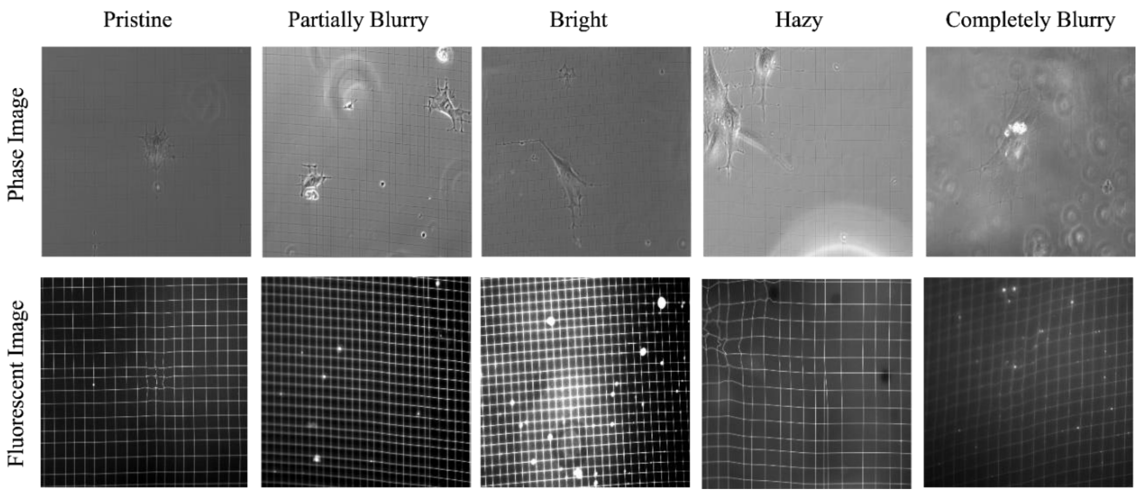

Supplementary Figure 6: An example phase and its corresponding fluorescent image are shown for each one of the five different type of training sets (Pristine, Partially Blurry, Bright, Hazy, and Completely Blurry) that were created for the ablation study.

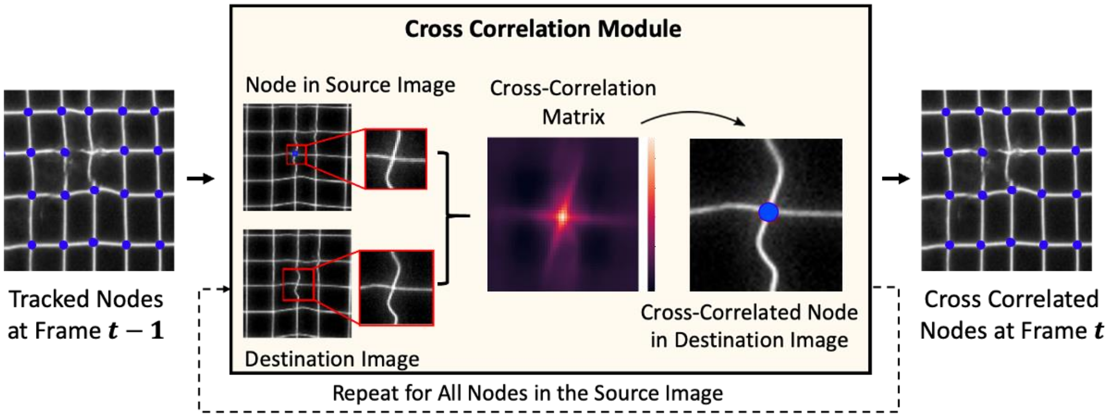

Supplementary Figure 7: A schematic representation of the Cross-Correlation Module.

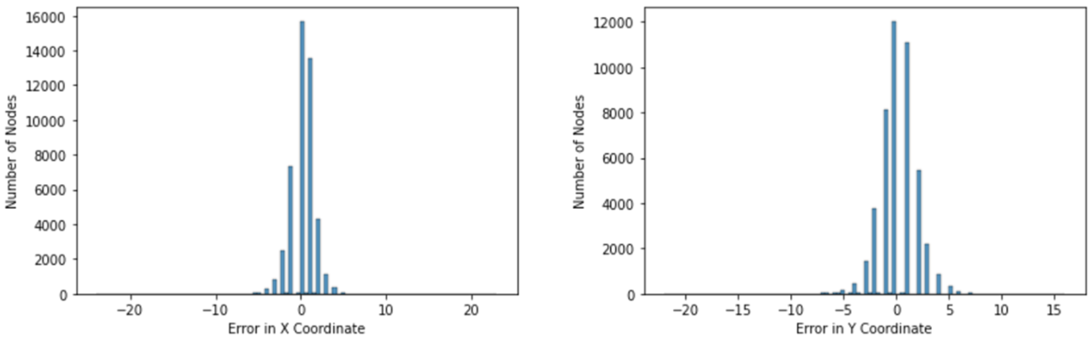

Supplementary Figure 8: Distributions of the errors in the predicted node coordinates along the X and Y dimensions across nodes of all the 17 videos.

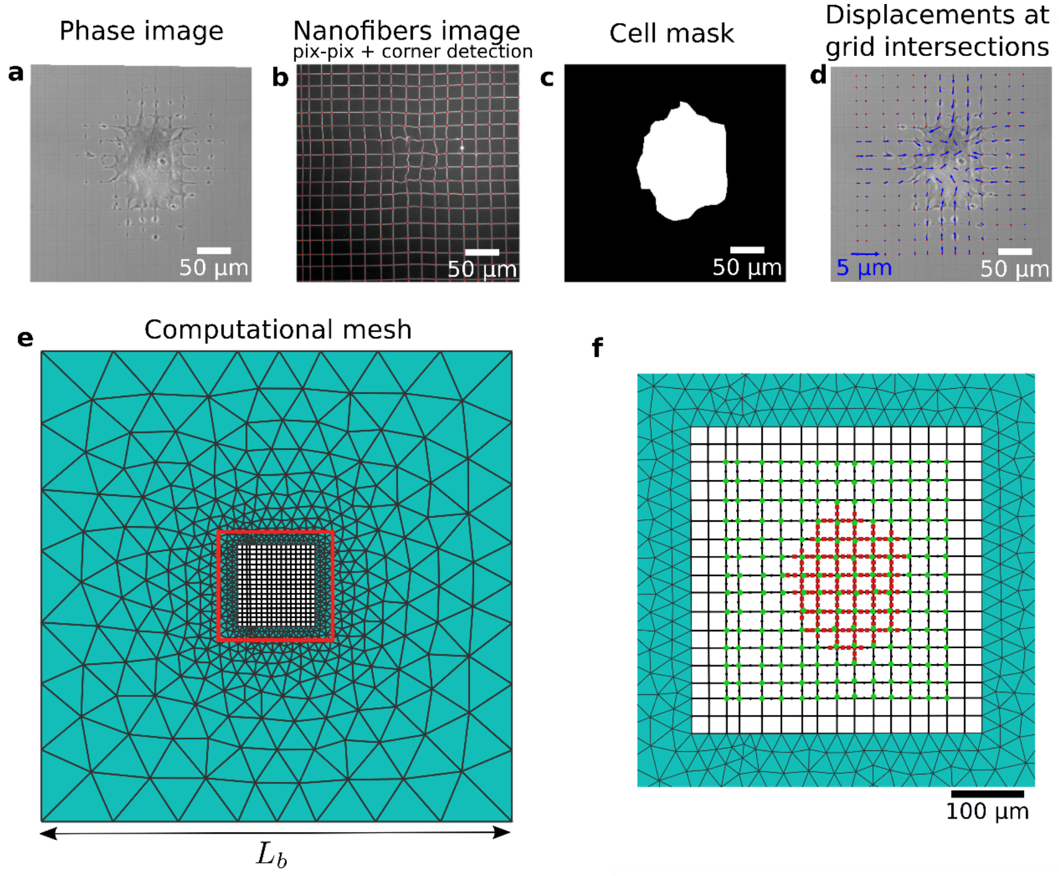

Supplementary Figure 9: Steps involved in generating the computational mesh starting from the experimental phase image of a migrating cell shown in (a). (b) The pix-pix method extracts the nanonet from the phase image, followed by corner detection algorithm to identify the grid intersection shown as red dots. (c) A cell mask is manually generated to outline the region occupied by the cell. (d) The displacements at the grid intersections are obtained using the reference grid positions obtained using the artificial reference generation method. (e) The computational mesh has two parts, the imaged nanonet region represented using corotational beam elements, while the surrounding region is meshed using triangular elements representing a homogenized Cosserat continuum. The box size $L_b$ = 2000 µm is chosen to be sufficiently larger than the cell size. (f) The red boxed region in (e) shows the mesh used for the imaged nanonet. The nodes where the displacements are known are marked in green which forms the vector of knowns $\mathbf{u}_m$. The nodes in the cell mask region are marked in red which correspond to the nodes at which the unknown force vectors are defined using $\mathbf{f}$.

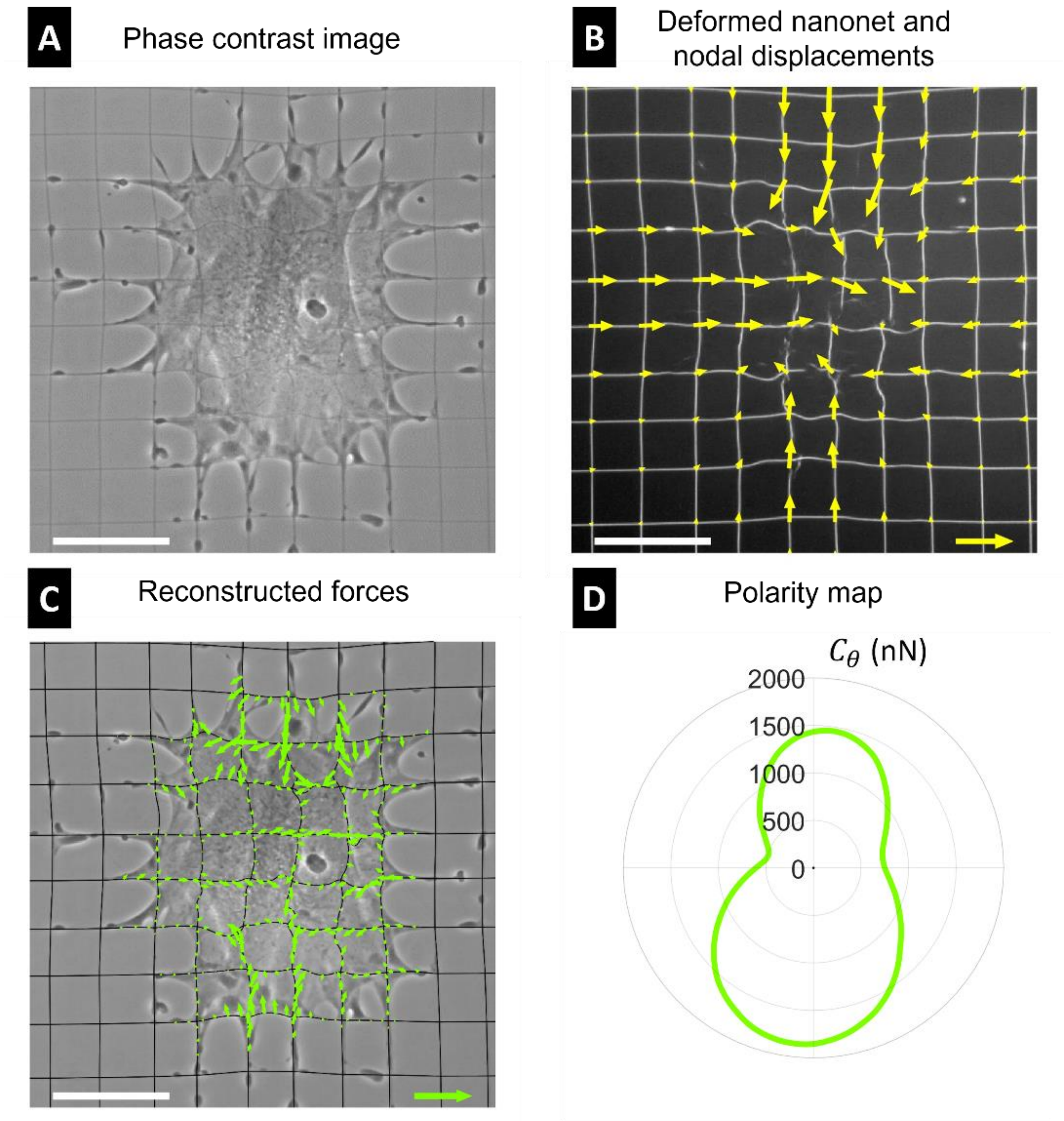

Supplementary Figure 10: Application of the inverse approach on experimental data: (A) phase image, (B) fiber shapes obtained by pix2pix and extracted displacements at fiber intersections, (C) reconstructed force pattern ($w_n$=1, $w_l$=3, $\beta$=0.01 nN/µm). (D) Polarity map showing contractility contribution $C_\theta$ along a direction specified by a unit vector an angle $\theta$ to the $x$-axis (see Methods section for the formula). White scale bar: (A-C) 50 µm. Displacement scale arrow: (B) 5 µm. Tension scale arrow: (C) 35 nN/µm.

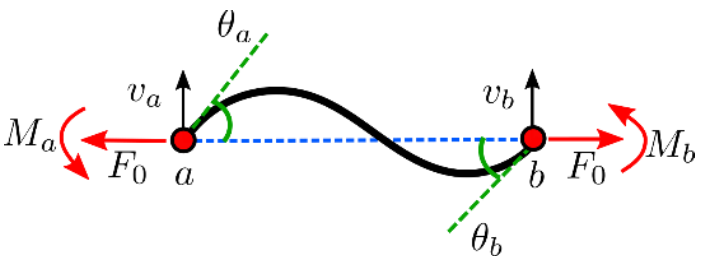

Supplementary Figure 11: Mechanical response of a pre-tensed Euler-Bernoulli beam segment *ab* of length $L$.

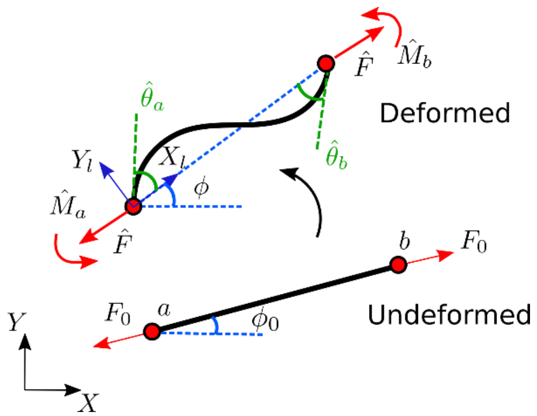

Supplementary Figure 12: Undeformed and deformed configurations of a beam segment *ab* are shown. A corotating frame $\{X_l, Y_l\}$ is attached with the deformed configuration to capture large rotations of the beam. The global fixed reference frame is denoted by the $X - Y$ axes.

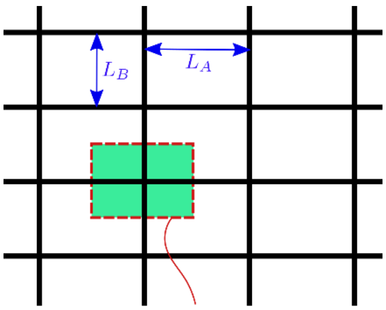

Supplementary Figure 13: Unit cell of a rectangular nanofiber grid with spacings $L_A$ and $L_B$ in horizontal and vertical directions, respectively.

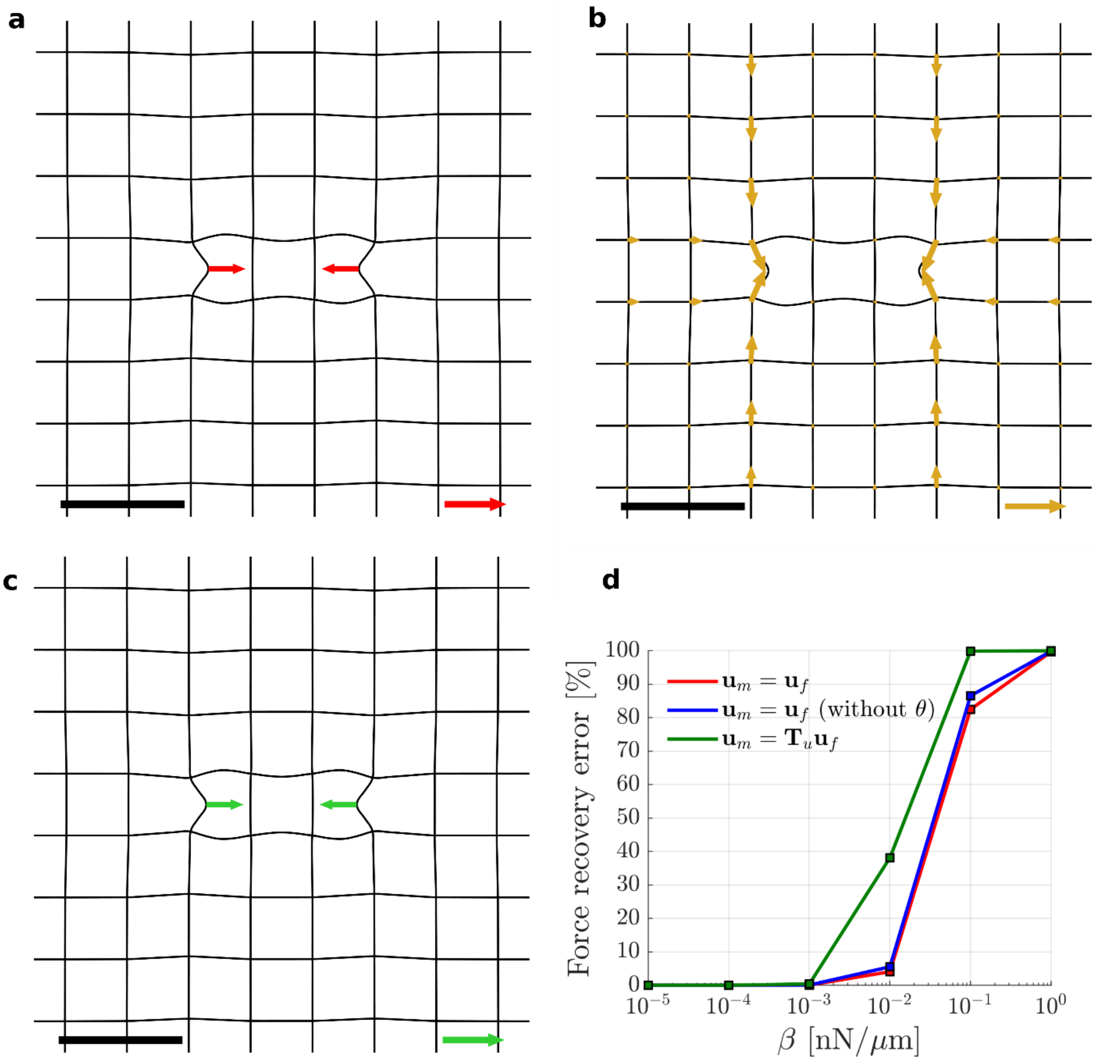

Supplementary Figure 14: (a) The force pattern $\mathbf{f}_f$ applied on the nanonet resulting in the deformation field shown in (b). (c) The recovered force pattern $\mathbf{f}$ for $\mathbf{u}_m = \mathbf{T}_u \mathbf{u}_f$ at $\beta = 10^{-4}$ nN/µm. (d) The error between recovered forces $\mathbf{f}$ and $\mathbf{f}_f$ as a function of the regularization parameter when using the full solution as measurement data ($\mathbf{u}_m = \mathbf{u}_f$), using only nodal displacements data, and using only displacements at grid intersections ($\mathbf{u}_m = \mathbf{T}_u \mathbf{u}_f$). The applied force pattern $\mathbf{f}_f$ is recovered to within 0.001% in all cases. (a)-(c) Black scale bar: 50 µm. (a) and (c): Force scale arrow: 1250 N (b) Displacement scale arrow: 3 µm.

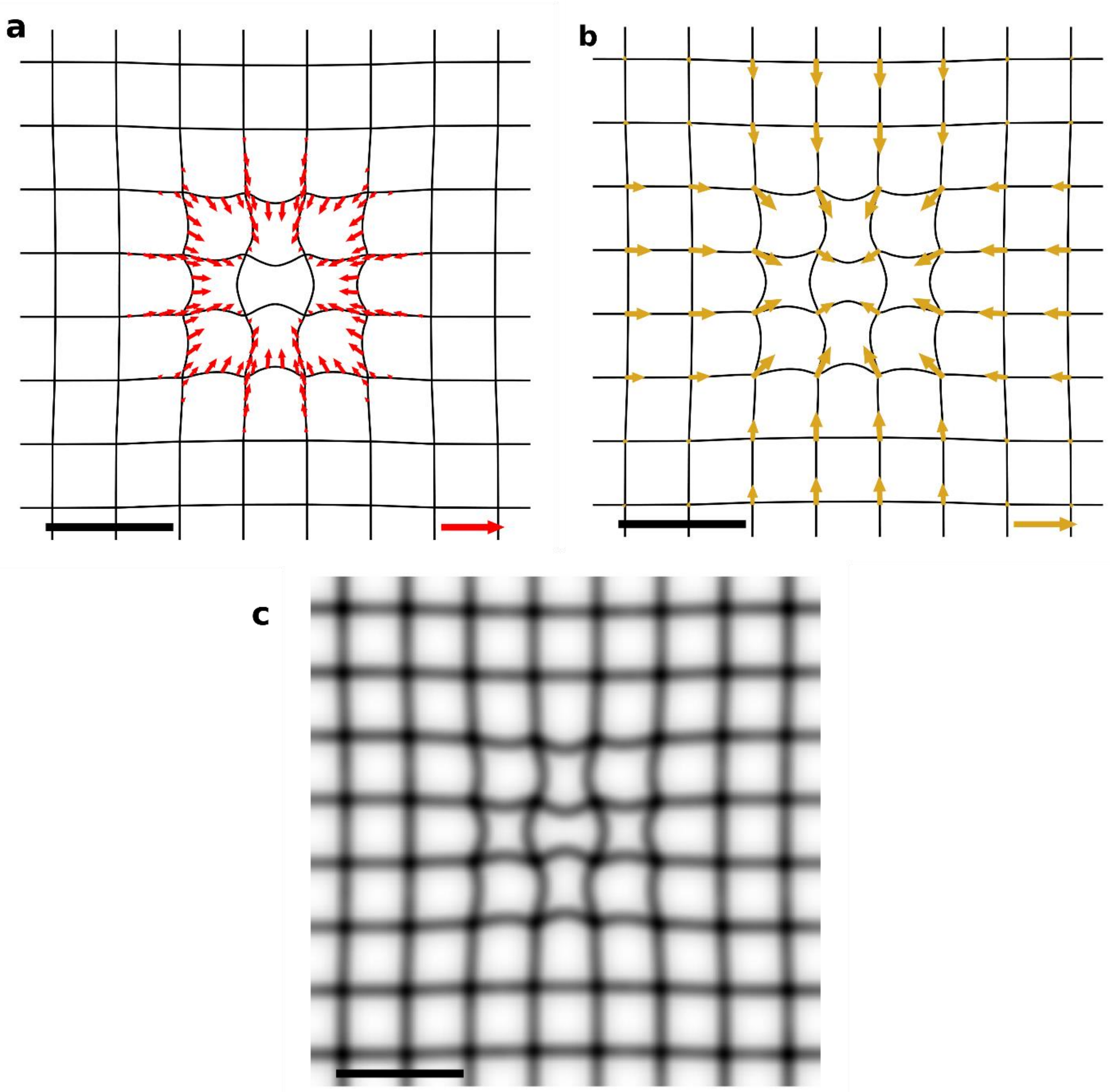

Supplementary Figure 15: (a) A radially symmetric contractile force pattern $\mathbf{f}_f$ applied on the nanonet. (b) The resulting displacements at the fiber intersections obtained using $\mathbf{u}_m = \mathbf{T}_u \mathbf{u}_f$.(c) The deformed nanonet is stored as a grayscale image to be used for active-contour regularization. (a)-(c) Black scale bar: 50 µm. (a): Line tension scale arrow: 25 mN/m. (b) Displacement scale arrow: 3 µm.

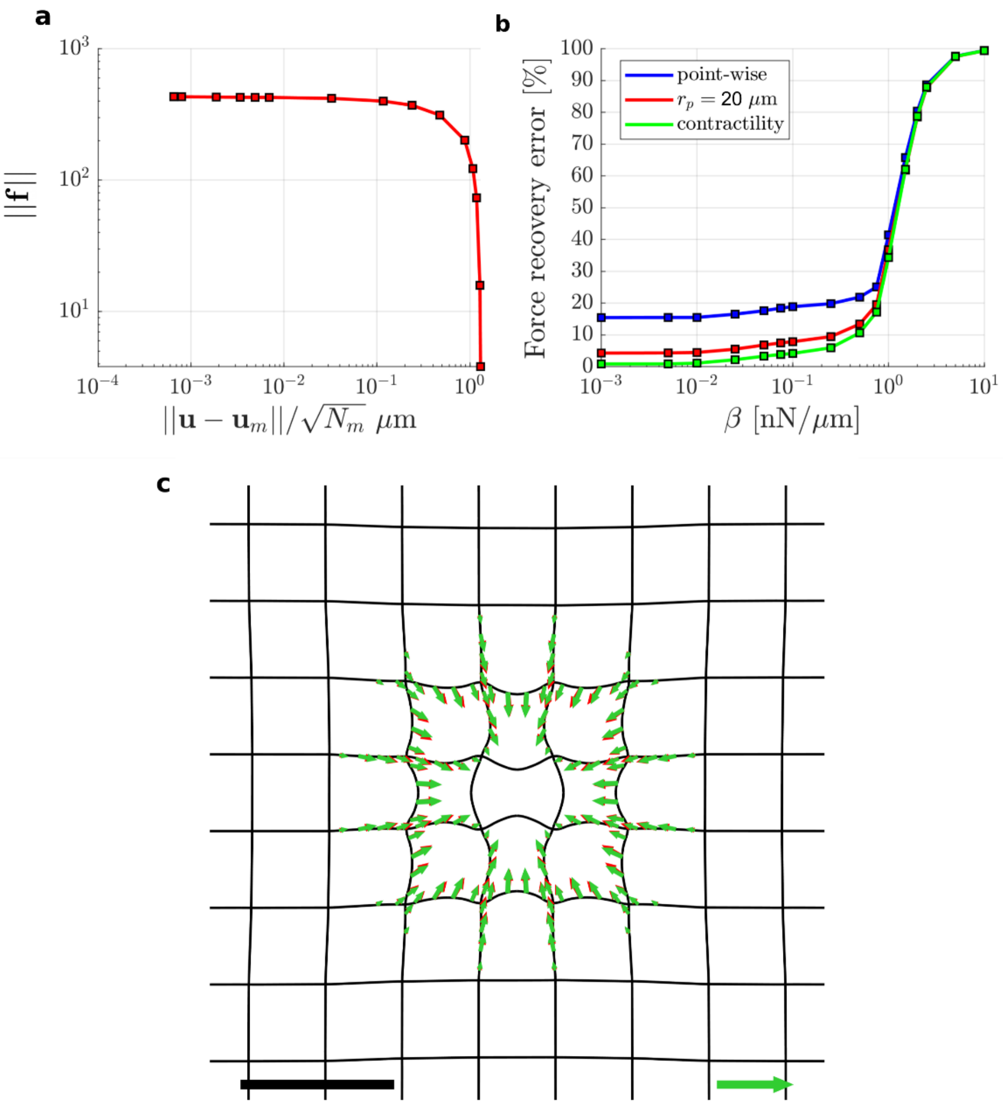

Supplementary Figure 16: Force recovery for the force pattern shown in Supplementary Figure 15 using the full solution $\mathbf{u}_m = \mathbf{u}_f$ without any noise. The inverse analysis is performed using $w_m = 1$ and $w_l = 0$. (a) L-curve obtained for the inverse analysis. Since $\sigma_e = 0$, we do not see a sharp increase in $||\mathbf{f}||$. (b) Relative error in the recovered forces $\mathbf{f}$ compared to $\mathbf{f}_f$. The error is calculated by point-wise direct comparison, by summing up forces within a radius of $r_p = 20$ μm about a given node, and by comparing the overall contractility $C$ of $\mathbf{f}$ and $\mathbf{f}_f$. (c) Comparison of the ground truth force pattern $\mathbf{f}_f$ (red) and the regularized solution $\mathbf{f}$ obtained at $\beta = 10^{-3}$ nN/μm. Black scale bar: 50 μm, Line tension scale arrow: 25 mN/m.

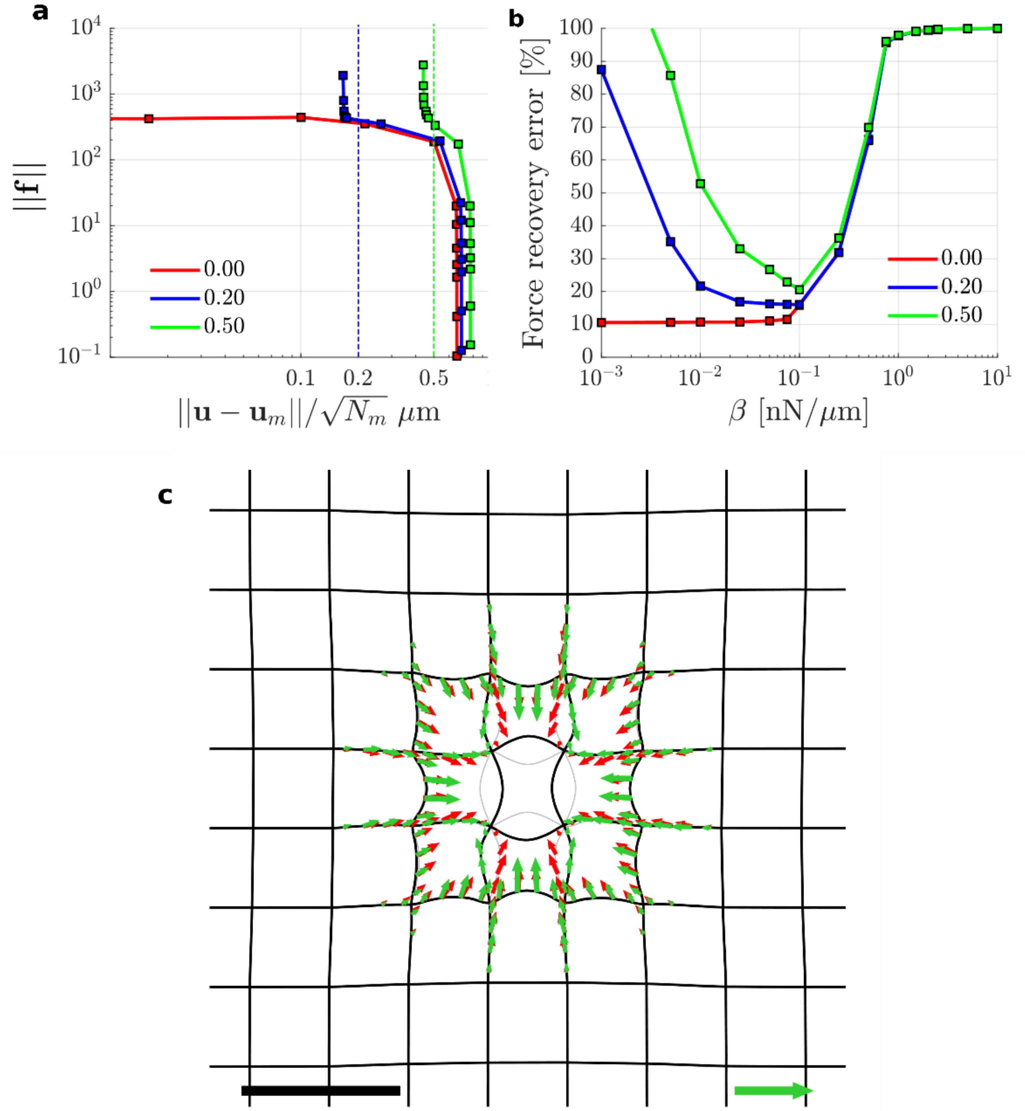

Supplementary Figure 17: Force recovery for the force pattern shown in Supplementary Figure 15 using at the fiber intersections $\mathbf{u}_m = \mathbf{T}_u \mathbf{u}_f$ with $\sigma_e$ ={0.0 µm, 0.2 µm, 0.5 µm}. The inverse analysis is performed using $w_m$ = 1 without active contour regularization *i.e.* $w_l$ = 0. (a) L-curves obtained from the inverse analysis for different $\sigma_e$. (b) Relative error in the recovered forces $\mathbf{f}$ compared to $\mathbf{f}_f$. The error is calculated by comparing the sum of forces within a radius of $r_p$ = 20 µm about a given node for $\mathbf{f}$ and $\mathbf{f}_f$. (c) Comparison of the ground truth force pattern $\mathbf{f}_f$ (red) and the regularized solution $\mathbf{f}$ obtained at $\beta$ = 0.1 nN/µm for $\sigma_e$ = 0.2. The gray lines represent the expected nanonet deformed shapes while the black lines represent the deformed nanonet obtained by solving the inverse problem. Black scale bar: 50 µm, Line tension scale arrow: 25 mN/m.

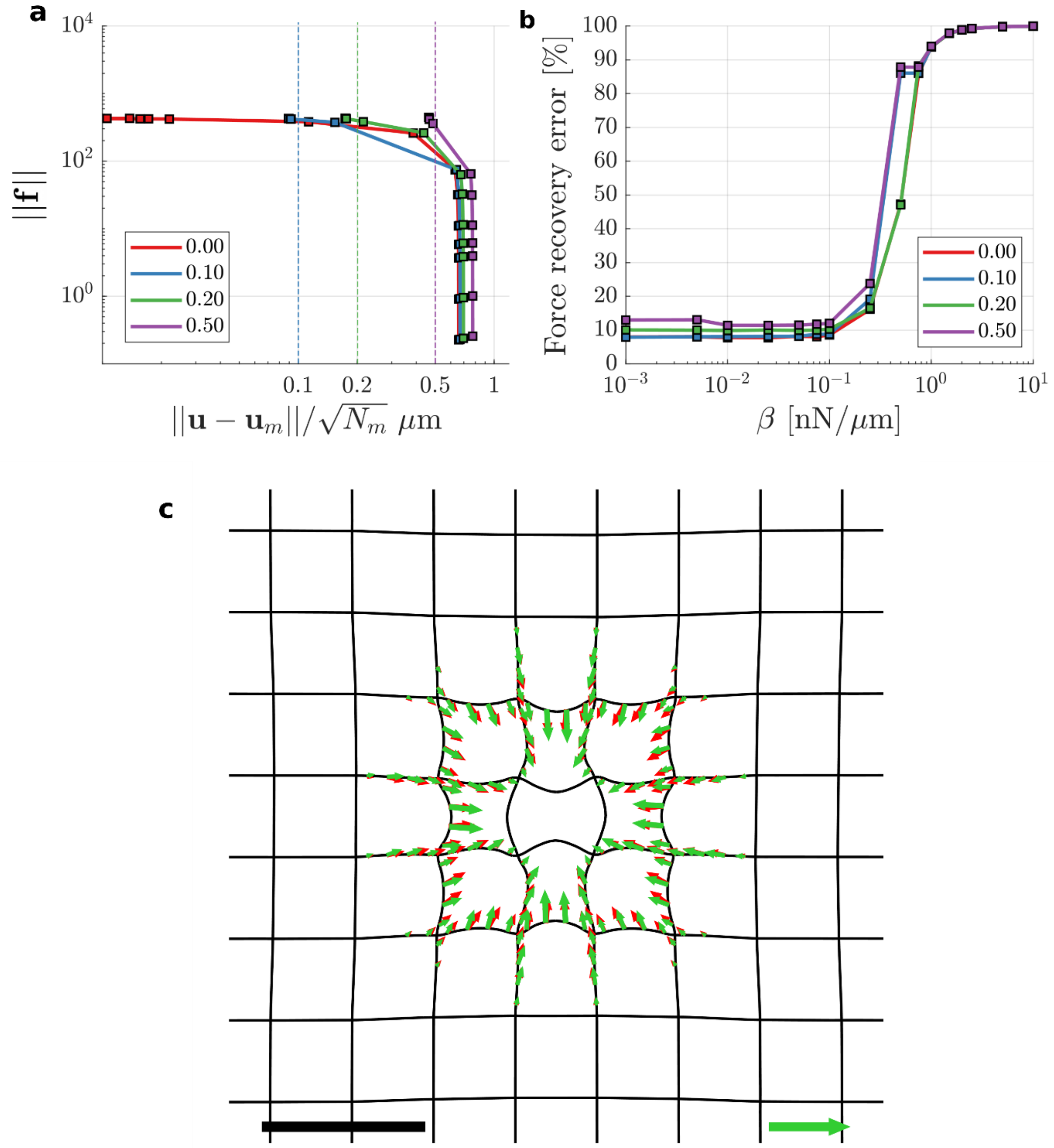

Supplementary Figure 18: Force recovery for the force pattern shown in Supplementary Figure 15 using at the fiber intersections $\mathbf{u}_m = \mathbf{T}_u\mathbf{u}_f$ with $\sigma_e$ ={0.0 μm, 0.2 μm, 0.5 μm}. The inverse analysis is performed using $w_m$=1 and with active contour regularization using $w_l$ = 3. (a) L-curves obtained from the inverse analysis for different $\sigma_e$. (b) Relative error in the recovered forces $\mathbf{f}$ compared to $\mathbf{f}_f$. The error is calculated by comparing the sum of forces within a radius of $r_p$ = 20 μm about a given node for $\mathbf{f}$ and $\mathbf{f}_f$. (c) Comparison of the ground truth force pattern $\mathbf{f}_f$ (red) and the regularized solution $\mathbf{f}$ obtained at $\beta$ = 0.1 nN/μm for $\sigma_e = 0.2$. Black scale bar: 50 μm, Line tension scale arrow: 25 mN/m.

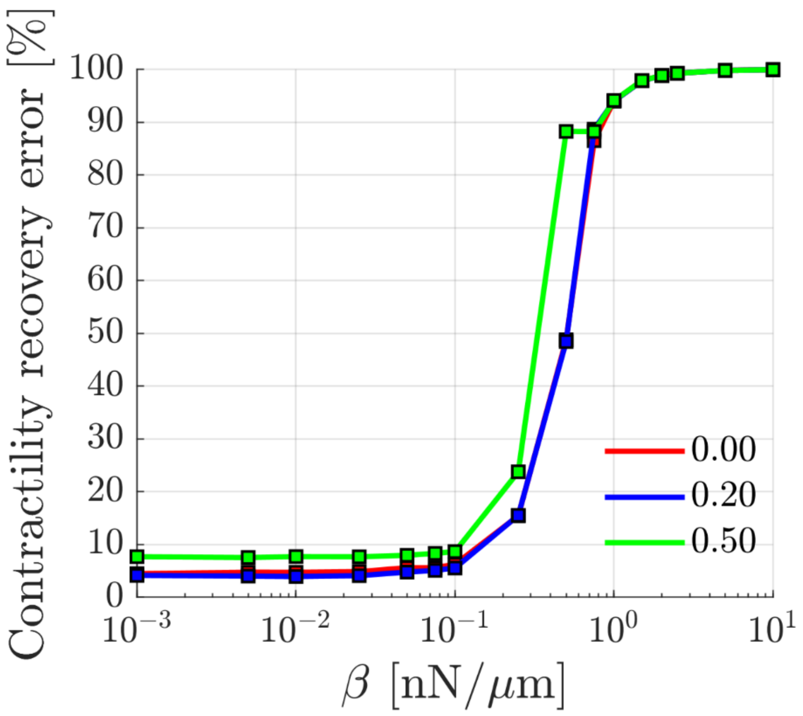

Supplementary Figure 19: Relative error in the contractility of recovered forces $\mathbf{f}$ compared to $\mathbf{f}_f$ shown in Supplementary Figure 18.

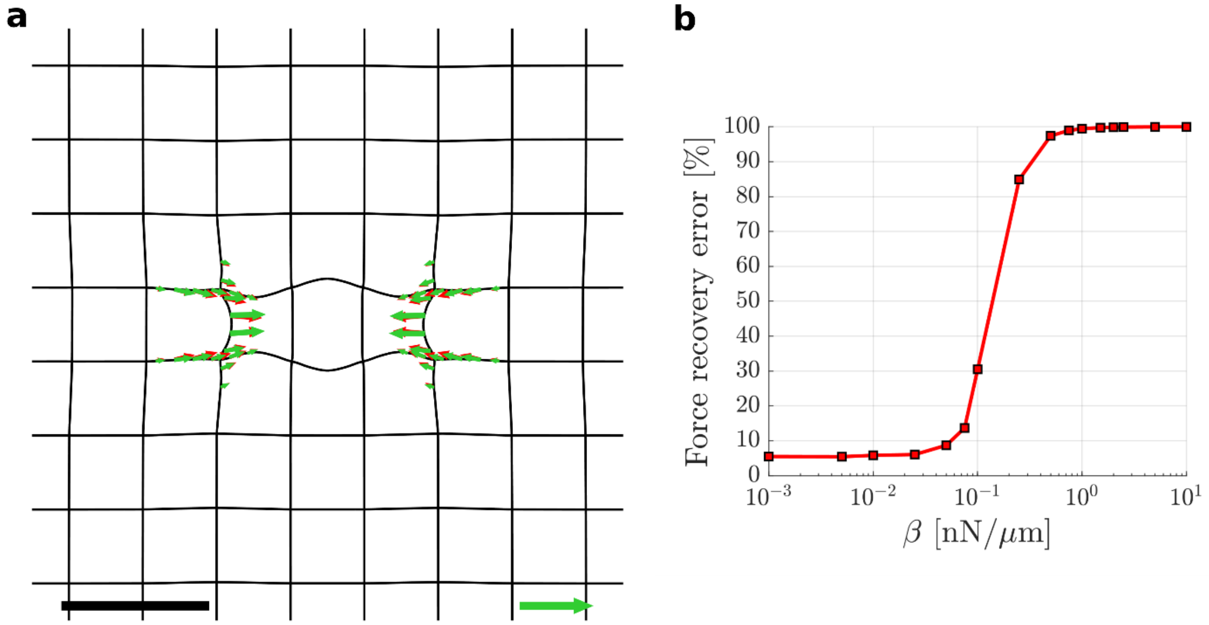

Supplementary Figure 20: (a) Comparison of the ground truth force pattern $\mathbf{f}_f$ (red) and the regularized solution $\mathbf{f}$ obtained at $\beta$ = 0.025 nN/µm for $\sigma_e$ = 0.2. (b) Relative error in the recovered forces $\mathbf{f}$ compared to $\mathbf{f}_f$ using $r_p$ = 20 µm. Black scale bar: 50 µm, Line tension scale arrow: 35 mN/m

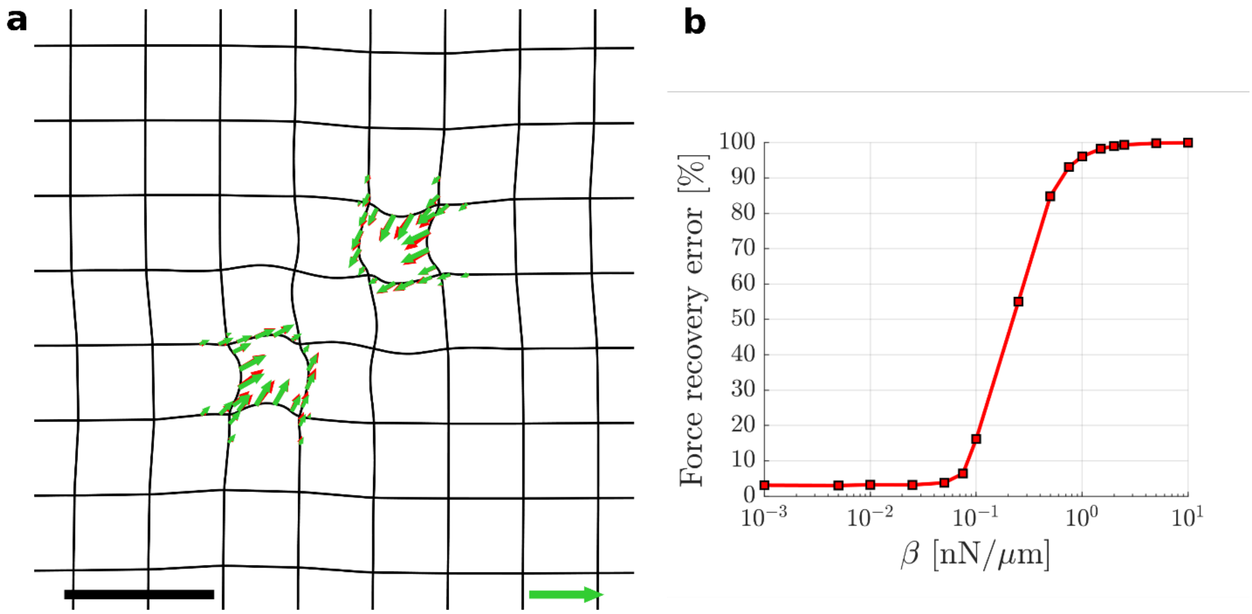

Supplementary Figure 21: (a) Comparison of the ground truth force pattern $\mathbf{f}_f$ (red) and the regularized solution $\mathbf{f}$ obtained at $\beta$ = 0.025 nN/µm for $\sigma_e$ = 0.2. (b) Relative error in the recovered forces $\mathbf{f}$ compared to $\mathbf{f}_f$ using $r_p$ = 20 µm. Black scale bar: 50 µm, Line tension scale arrow: 35 mN/m

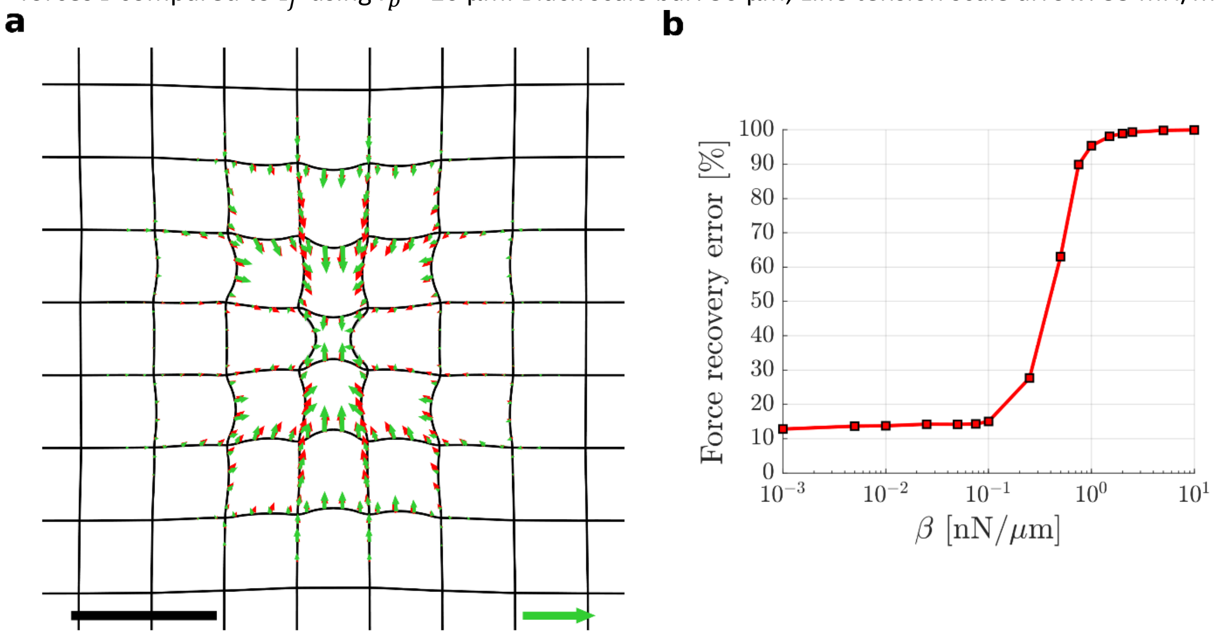

Supplementary Figure 22: (a) Comparison of the ground truth force pattern $\mathbf{f}_f$ (red) and the regularized solution $\mathbf{f}$ obtained at $\beta$ = 0.025 nN/µm for $\sigma_e$ = 0.2. (b) Relative error in the recovered forces $\mathbf{f}$ compared to $\mathbf{f}_f$ using $r_p$ = 20 µm. Black scale bar: 50 µm, Line tension scale arrow: 50 mN/m

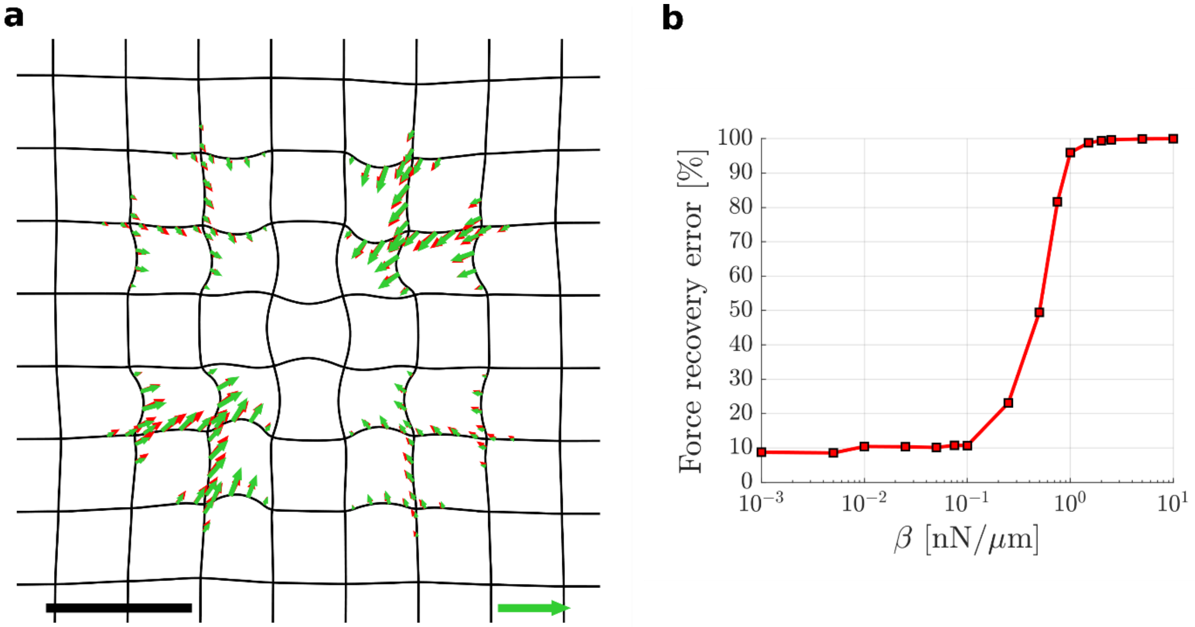

Supplementary Figure 23: (a) Comparison of the ground truth force pattern $\mathbf{f}_f$ (red) and the regularized solution $\mathbf{f}$ obtained at $\beta$=0.025 nN/µm for $\sigma_e$=0.2. (b) Relative error in the recovered forces $\mathbf{f}$ compared to $\mathbf{f}_f$ using $r_p$=20 µm. Black scale bar: 50 µm, Line tension scale arrow: 35 mN/m

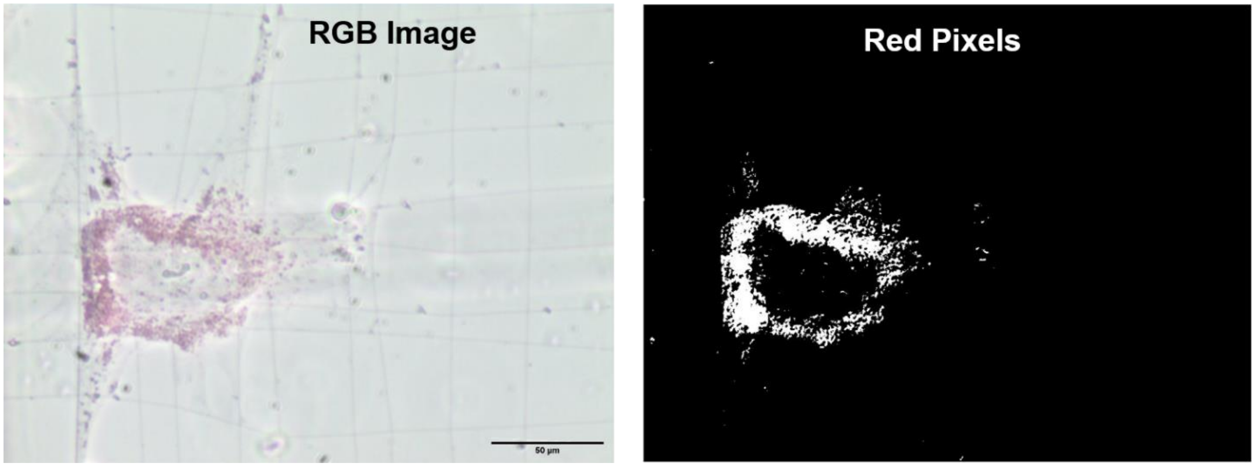

Supplementary Figure 24: RGB image obtained through a colored Amscope camera MU300 shows the Oil Red O stained lipid droplets on cell placed in adipogenic differentiation medium. Red pixels are extracted from the RGB image based on the criteria $\frac{R}{\sqrt{R^2+G^2+B^2}} > 0.6$ where $R$, $G$ and $B$ denoted the intensity value in the corresponding channel. Scale bar: 50 µm.

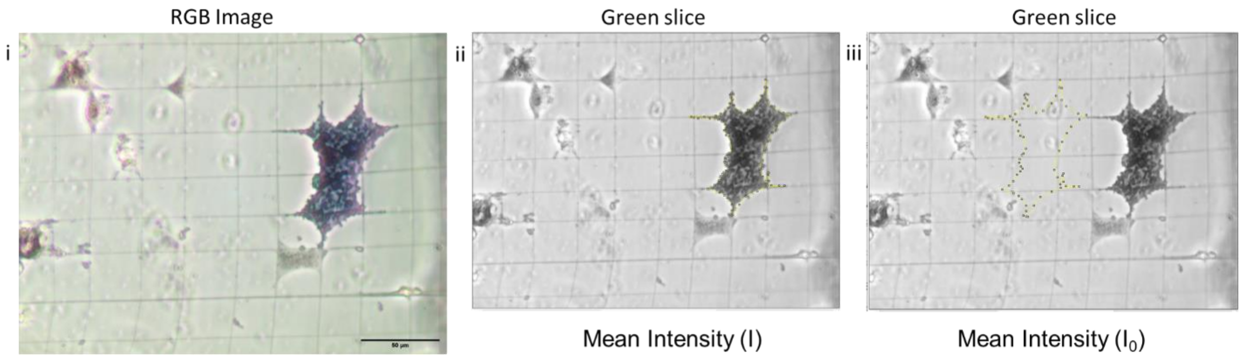

Supplementary Figure 25: (i) RGB image obtained through a colored Amscope camera MU300 shows the ALP stain on cell placed in osteogenic differentiation medium. (ii) Mean intensity (I) of area covered by cell obtained through manual outlining. (iii) Mean intensity ($I_0$) obtained by using the same cell mask for background region. ALP activity is computed using the expression $S_{\text{cell}} = \log \frac{I_0}{I}$ where $S_{\text{cell}}$ denotes the number of pixels covering the spread area of cell. Scale bar: 50 µm.
